# A deep learning approach for the discovery of tumor-targeting small organic ligands from DNA-Encoded Chemical Libraries

**DOI:** 10.1101/2023.01.25.525453

**Authors:** Wen Torng, Ilaria Biancofiore, Sebastian Oehler, Jin Xu, Jessica Xu, Ian Watson, Brenno Masina, Luca Prati, Nicholas Favalli, Gabriele Bassi, Dario Neri, Samuele Cazzamalli, Jianwen A. Feng

## Abstract

DNA-Encoded Chemical Libraries (DELs) emerged as efficient and cost-effective ligand discovery tools, which enable the generation of protein-ligand interaction data of unprecedented size. In this article, we present an approach that combines DEL screening and instance-level deep learning modeling to identify tumor-targeting ligands against Carbonic Anhydrase IX (CAIX), a clinically validated marker of hypoxia and clear cell Renal Cell Carcinoma. We present a new ligand identification and HIT-to-LEAD strategy driven by Machine Learning (ML) models trained on DELs, which expand the scope of DEL-derived chemical motifs. CAIX screening datasets obtained from three different DELs were used to train ML models for generating novel HITs, dissimilar to elements present in the original DELs. Out of the 152 novel potential HITs that were identified with our approach and screened in an *in vitro* enzymatic inhibition assay, 70% displayed submicromolar activities (IC_50_ < 1 μM). Based on the first HIT set, the model was further used to prioritize and generate LEAD compounds with nanomolar affinity for *in vivo* tumor-targeting applications. Three LEAD candidates showed accumulation on the surface of CAIX-expressing tumor cells in cellular binding assays. The best compound displayed *in vitro* K_D_ of 5.7 nM and selectively targeted tumors in mice bearing human Renal Cell Carcinoma lesions. Our results demonstrate the synergy between DEL and machine learning for the identification of novel HITs and for the successful translation of LEAD candidates for *in vivo* targeting applications.

**Graphical Abstracts:** 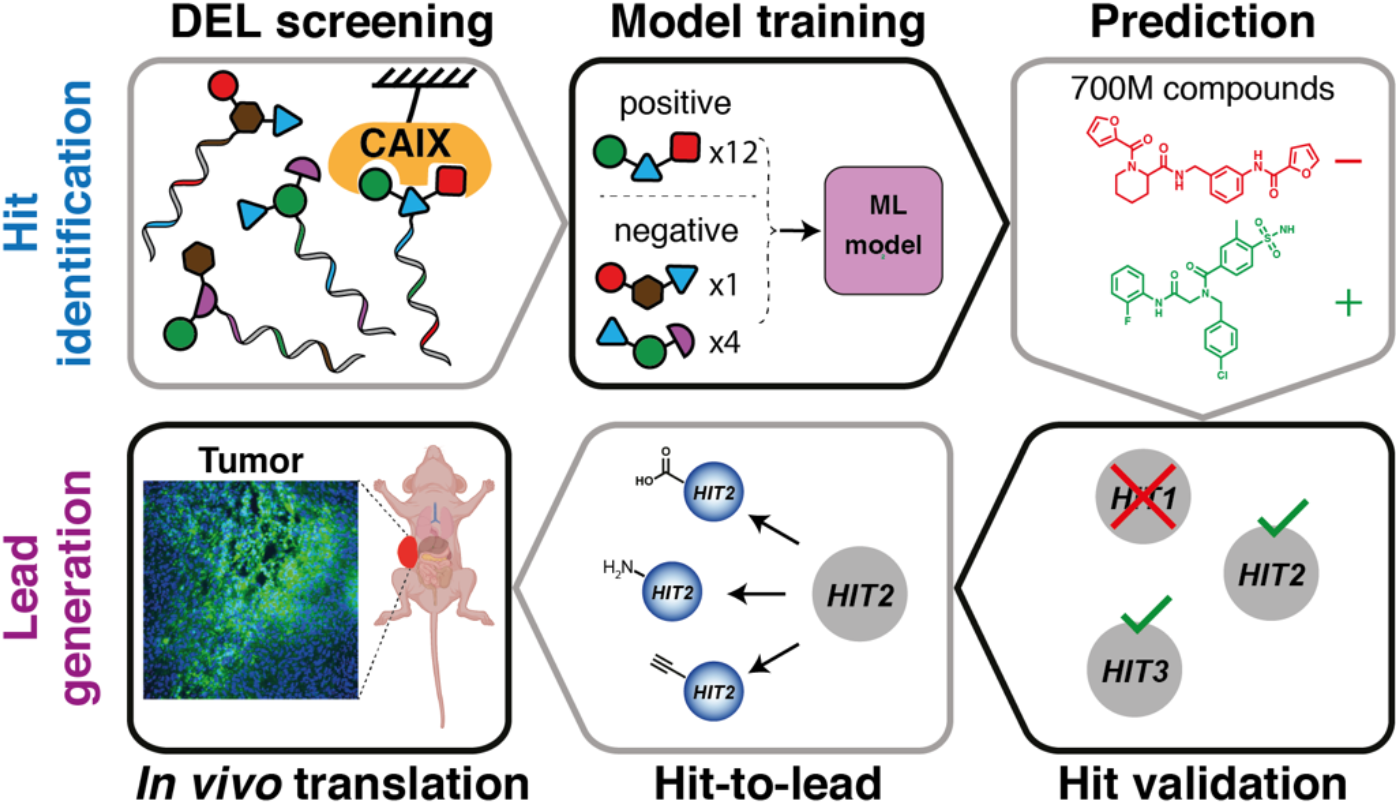

## Introduction

Small organic ligands which specifically interact with protein targets overexpressed in cancer lesions are increasingly being considered for the selective delivery of therapeutic payloads to the site of diseases.[1]–[4] Most of the ligands used for pharmacodelivery applications have been generated on the basis of natural substrates of tumor-associated antigens.[5]–[7] Lutathera® and Pluvicto®, two recently approved Radioligand Therapeutics (RLTs) for the treatment of gastroenteropancreatic Neuroendocrine Tumors (GEP-NETs) and metastatic Castration-Resistant Prostate Cancer (mCRPC), are based on derivatives of previously known binders of the respective molecular targets (i.e., Somatostatin Receptor-2 and Prostate Specific Membrane Antigen, respectively). Nature has been a productive source of binding specificities, but *de novo* ligand discovery remains challenging.[1], [8] Interrogation of chemical compound collections has been miniaturized and automatized in the form of high-throughput screening (HTS) technologies.[9] While HTS promised to deliver ligands for any protein target of interest, practical application of this technology by pharmaceutical companies is limited by high set up costs and time-consuming screening protocols.[8]–[10]

DNA-encoded chemical libraries (DELs) have evolved as efficient and cost-effective ligand discovery tools in alternative to HTS.[11]–[17] DELs are pools of organic chemical compounds which are individually linked to DNA “barcodes”. In a typical DEL selection, millions to billions of DEL members are screened against the target protein of interest, immobilized on a solid support. Large sets of data are obtained thanks to High Throughput DNA Sequencing (HTDS) technologies aimed at identifying the barcodes uniquely associated to preferentially enriched compounds.[17] With increasing library size, complexity, and sequencing capacity, it has become more challenging to interpret and exploit large data sets which result from DEL screening campaigns.[9], [18], [19] Computational methods have been developed to facilitate the identification of potential protein binders (HITs) and study Structure-Activity Relationships (SARs).[20], [21] Quantitative analysis based on negative binomial distribution,[18], [22], [23] enrichment metrics that factor in different sources of uncertainties,[24]–[26] and modeling approaches to de-noise DELs accounting for partial products[27] were successfully applied for HIT prioritization.

Building predictive models on DEL-screening datasets is challenging due to various confounding factors such as varying chemical yields of expected structures during library synthesis,[28] nonuniform baseline abundances of library members, and substantial under-sampling.[26] To cover a diverse chemical space, modern DEL experiments are often multiplexed[29], where tens to hundreds of DEL libraries are pooled together during selections and sequencing.[26], [30], [31] This allows billions of compounds to be screened against the target in a single experiment. However, due to sequencing time and cost, a typical sequencing experiment generates 10^6^ -10^8^ of reads, which is only a small fraction of the total chemical diversity that can be reached in the library pool. With a low sampling ratio, sequencing count distributions are often dominated by shot noise, resulting in low signal-to-noise ratios and poor reproducibility between experimental replicates.[26]

In order to address under-sampling while building predictive models on DEL datasets, de-noising strategies such as disynthon aggregation[9] have been employed. Sequencing reads of molecules which share common two-cycle building blocks are aggregated to generate “disynthon-level” enrichment scores and classification labels for model training.[32] However, during disynthon aggregation, structural information from individual molecules is partially lost. Additionally, aggregation over a middle-position building block can generate molecules which cannot be practically synthesized. Several groups have recently proposed probabilistic models that operate on the full DEL compounds (“instance-level”) and have demonstrated promising results in retrospective evaluation.[33]–[35] However, there have not been prospective studies to validate such models in real world HIT-finding applications.

In this article, we developed an approach that combines DEL screening and modeling to identify and generate LEAD compounds against Carbonic Anhydrase IX (CAIX), a clinically validated marker of hypoxia and clear cell Renal Cell Carcinoma.[36]–[42] We introduced an instance-level deep learning approach on screening results of three different DELs. The initial model resulted in novel HITs (not present in the original chemical collections) which were characterized by enzymatic inhibition assay against the target protein. The model was then further applied to prioritize and generate a list of LEAD candidates for *in vivo* applications. A selection of LEAD compounds was found to bind to the surface of CAIX-positive cancer cells and to selectively target tumors in mice bearing human Renal Cell Carcinoma lesions. Our results demonstrate the synergy between DEL and machine learning for the identification of novel HITs and for the successful translation of LEAD candidates for *in vivo* targeting applications.

## Material and Methods

### Datasets

Our training dataset consists of three DEL libraries with 4.2 million, 1.57 million and 53,326 members respectively, each constructed with 3 synthesis cycles. Schematic structures of the three DELs are shown in **Figure 1**. For each DEL library, affinity-mediated selections for CAIX (target selection) and selections without the presence of the target protein (no target control, NTC) were performed. During the construction of the three DEL libraries, sulfamoylbenzoic acid (SABA) derivatives were included as DEL building blocks, resulting in 1.3% SABA-containing compounds in the overall DEL dataset. To reduce noise in DEL sequencing counts, selections and sequencing were performed for each individual DEL separately, resulting in sampling ratios (defined as the ratio of sequencing read depth to the number of DEL members) that are on the order of 1 (**Table S1**), which is thousand times higher than typical multiplex DEL screenings.[26] Such setup enables high reproducibility between experimental replicates (**Figure S1** and **S2**).

**Figure 1.**
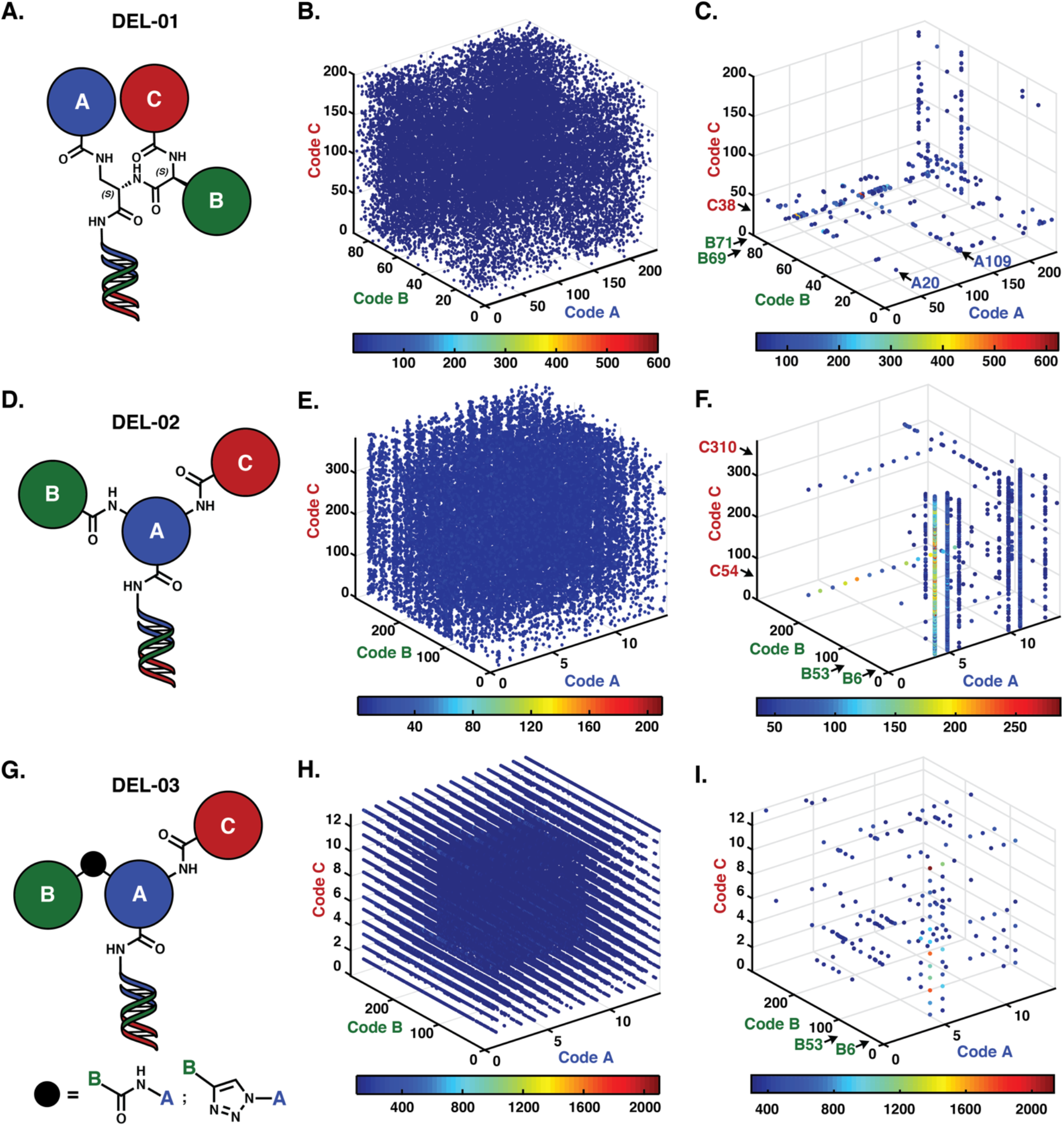
DEL training dataset. **A.,D.,G.**, Chemical structures of DELs. **B.,E.,H.** High-Throughput DNA Sequencing (HTDS) plots after library selections against unmodified streptavidin-coated beads. The x, y and z axes correspond to code A, B, and C, respectively. The colored jet scale indicates DNA sequence counts. Cut-off = 4; 4 and 100, respectively. Total counts (**B., E., H.**) = 1,551,875; 1,817,134 and 1,617,433, respectively. These results are used in data analysis as negative controls to evaluate selection results. **C.,F.,I.** HTDS plots after library selections against Carbonic Anhydrase IX (CAIX). Cut-off = 40; 30; 300, respectively. Total counts = 2,837,727; 1,964,585 and 2,533,971, respectively. Building block combinations enriched in selections against CAIX are indicated by black arrows. Selections were performed in duplicates (see **Figure S1**).

### Input featurization and processing

To train our machine learning models, the DEL compounds and screening results need to be represented in model readable formats. This requires two data preprocessing steps: (1) Computationally enumerate individual DEL compound structures from their corresponding building blocks and represent them as molecular graphs (2) Create training labels for each DEL compounds based on their sequencing counts.

#### SMILES enumeration and small molecule representation

SMARTS-based enumeration was used to generate SMILES (simplified molecular input line entry system) representations of the DEL compounds. For each synthesis cycle, RDkit was used to perform in-silico reactions to generate the products from the building blocks and the corresponding reaction SMARTS. The fully enumerated products were then represented as molecular graphs, where the nodes represent individual atoms and the edges represent bonds, with atom and bond features specified by Kearnes *et al*.[43]

#### Enrichment score computation and example labeling

For each of the target and no target control selection types, we compute normalized z-scores[24] from the raw sequencing counts. The normalized z-score calculation approximates the DNA sequencing process as random sampling with replacement using a Binomial distribution and quantifies the level of enrichment of an observed count compared to the expected count (e.g., uniform prior), factoring the sequencing read depth and library sizes. Specifically, the formulation is described in EQ 1:

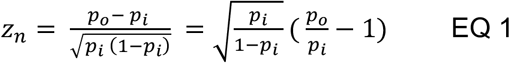

where p_o_: observed probability (p_o_ = c_o_ / n), p_i_: expected probability (p_i_ = 1/library size), c_o_: observed count, n: total sequencing reads.

From the normalized z-scores, we define classification labels to categorize each DEL compound into one of the three different classes: MATRIX_BINDER, TARGET_HIT and NON_HIT. The classes are defined as follow: (1) MATRIX_BINDER: Examples with NTC normalized zscore >= 0.004 are considered enriched in the no target control selection and are labeled as MATRIX_BINDERs. (2) TARGET_HIT: Examples with NTC normalized zscore < 0.004 AND target selection normalized zscore >= 0.004 are labeled as TARGET_HITs. (3) NON_HIT: All the other examples were labeled as NON_HITs. With this process, the TARGET_HIT class included DEL compounds that were enriched in the target condition, but not enriched in the matrix only condition, and is defined as the “positive” class for our machine learning model. From this process, 37,928 examples were labeled as TARGET_HIT.

### Model design, training and evaluation

#### Model architecture

Following the work by Mccloskey *et al*.[32], we employed a Graph Convolutional Neural Network (GCNN) with weave modules (“W2N2” variant), where the input features and hyperparameters were as specified by Kearnes *et al*.[43] The max number of atoms per compound was set to 70 to account for the larger, instance-level compounds. The final linear layer of the model makes predictions on the three mutually exclusive classes [NON_HIT, MATRIX_BINDER, TARGET_HIT], trained with softmax cross-entropy loss. GCNN was selected as our model architecture because it was demonstrated in Mccloskey *et al*.[32] to outperform Random Forest models in HIT finding.

#### Cross-validation and evaluation metrics

To evaluate model performance retrospectively, we employed a k-fold cross validation scheme, which divides the DEL data into train, validation, and test splits. The k folds were determined by affinity clustering,[44] a method to perform clustering on weighted (where the edge weights represent similarity), undirected graphs. Specifically, we created compound-similarity-based clusters by the following steps: (i) Compute pairwise molecular similarities between all DEL compounds with Extended-Connectivity Fingerprints[45] (ECFP), radius 3 (ECFP6). (ii) Construct a large graph where each compound is a single node and two nodes are connected with a weighted edge if they have fingerprint similarity >= 0.5. The weight of the edge is the molecular similarity between the two nodes. (iii) Affinity clustering is then run on the constructed graph to identify the best neighbors (prioritized by weights) of each node. The resulting connected subgraphs are then outputted as different clusters. The algorithm can be run with different levels, with higher levels corresponding to larger clusters. For this study, results at level 7 were used, resulting in 5 clusters (5 folds).

With the created fold splits, k − 2 folds were then used for training the GCNN, with one validation fold set aside for model selection and one test fold reserved. The process is repeated 5 times, with each of the folds being used as the validation fold. For each of the cross-validation fold models, two different metrics are computed for the TARGET_HIT class: (1) The Area Under the Curve (AUC) of the Receiver Operator Characteristic (ROC) curve[46], or ROC-AUC, which quantifies the overall ability of the model to classify CAIX HITs against non-HITs across different score thresholds and (2) top_100_positives, defined as the number of true TARGET_HIT class examples in the top-100-scored compounds in the validation fold. This metric quantifies the early enrichment of the model, which mimics the actual use case in a drug discovery program where only the top k candidates are experimentally validated.

### Batch sampling

The three classification classes are highly imbalance (with the NON_HIT class dominating the training data), and the affinity-cluster based cross-validation folds varied substantially in size. To ensure the model is presented with examples from different classes and chemical spaces, we follow a similar oversampling strategy outlined in Mccloskey *et al*.,[32] which over-samples examples from the under-represented classification classes and cross-validation folds. Additionally, due to the high enrichment of SABA derivatives in the TARGET_HIT class, we additionally enforced a strategy to sample evenly from SABA-containing and non-SABA-containing examples per classification class. This results in a sampling strategy where each minibatch contains equal numbers of examples from different (fold, classification class, SABA-containing or non-SABA-containing) categories. Effectively, the additional SABA-based batch creation strategy up-samples the TARGET_HITs that do not contain SABA and non-HITs that contain SABA.

### Model training and selection

Each of the GCNN models was trained to converge on 1 Tensor Processing Units (TPUs) with 8 TensorNodes. Each model comprises 8 independently randomly initialized TPU replicas. Each of the 8 TPU replicate models converged independently, and the median of the predictions from the 8 replicates is used as the overall prediction of a single model. To assess the variability of the GCNN models, we train 3 independent models with different random initializations for each fold, resulting in 15 models (5 cross-validation fold, 3 independent model replicas, each with 8 TPU replicates). Each GCNN model converged within 24 hours. For each cross-validation fold model, the model weights at the training step with the maximum TARGET_HIT class top_100_positive validation fold metric were selected. Among the selected models, the average cross-validation TARGET_HIT ROC-AUC and top_100_positive metrics are 0.88 and 71.3 respectively.

### HIT finding: Inference and diversity selection on commercial catalogs

The selected best models were used to make predictions on Enamine REAL (735.15M compounds, version 2019)[47] and Mcule Instock (9.35M compounds, version 2021).[48] For a given test compound, the median prediction across the 15 models (5 cross-validation fold, 3 replicates) was used as the final prediction. The following process was then applied to select a set of diverse, high-scored and drug-like compounds for experimental validation. (1) Top scoring compounds that received prediction scores higher than a pre-specified threshold (0.8) were considered as the candidate compound sets. (2) A set of pre-defined property filters were then applied to remove compounds that are non-drug-like or reactive. Briefly, compounds weighing >600 Da, containing more than 4 aromatic rings or more than 7 rotatable bonds were removed. A set of SMARTS patterns were further used to perform substructure search to remove compounds that may be toxic or reactive. The full set of filtering criteria is described in (**Table S2**). (3) Directed sphere exclusion[49] (DISE) was applied using ECFP6 Tanimoto similarity (cutoff = 0.7 for Enamine and cutoff = 0.65 for Mcule), ranked by the model prediction score. Following the above steps, we selected 125 compounds from Enamine and 47 compounds from Mcule. 108 and 44 compounds were delivered from Enamine and Mcule respectively.

### HIT-to-LEAD: Analog search, enumeration and prioritization

Given the potent and diverse starting HITs, to develop tumor-targeting ligands against CAIX that can be functionalized with imaging payloads, we aimed to identify analogs of the initial HITs that contain reaction handles for amidation and CuAAC click chemistry. To achieve this, we performed analog search and model-guided analog prioritization, as detailed below.

#### Analog search in Enamine REAL and enumeration of derivatives

We first performed substructure search in Enamine REAL (1.9B compounds, version 2021) to identify readily available compounds that contain the desired reaction handles (primary amines, carboxylic acids, and alkynes) or their protected variants. For each starting HIT, we then perform similarity searches within the identified Enamine subset to find analogs with ECFP6 Tanimoto similarity > 0.5 to the initial HIT. For starting HITs where reaction-handle-containing analogs were not available in Enamine, we computationally enumerated the amine/acid/alkyne derivatives. The attachment points of the reaction handles were determined based on heuristics on synthesizability and positioning away from the aromatic sulfonamides.

#### Model driven analog prioritization

Since the reaction handles are expected to undergo chemical changes after conjugation, to predict the compounds’ ability to bind to CAIX in their conjugated form, for each of the proposed analogs, we computationally generated surrogate compounds where the reaction handles were replaced with the reacted form (capped) and re-scored the surrogate compounds with our trained models. Only compounds whose surrogate form scored above 0.8 were selected for experimental validation (**Figure S3**).

#### Purchasing and custom synthesis of the analogs

Commercially available analogs were directly purchased from Enamine while computationally enumerated derivatives were custom synthesized by Enamine. Additionally, we custom synthesized the surrogate compounds (capped version) for all analogs through Enamine to allow a two-stage testing strategy: i.) Validation of the surrogate compounds in enzymatic assays (ii) Conjugation to FITC and subsequent validation of the analogs with fluorescence polarization.

### Protein expression and purification

To produce recombinant human CAIX (amino acids 120-397) a CHO stable cell line was generated. Briefly, CHO cells were transfected with a pcDNA3.1 mammalian expression vector (Invitrogen) carrying a CAIX gene where the endogenous leader sequence was replaced by a murine IgG signal peptide and an hexa-histidine-tag sequence was fused at the 3’ end of the CAIX gene to facilitate purification. Transfected cells were cultivated for three weeks in PowerCHO-2CD median (LONZA) supplemented with 4mM Ultraglutamine-1 (Lonza) and 500mg/L G418 (Millipore) to obtain a pool of stably transfected cells. Single cells were then sorted by limiting dilution and a clone showing high CAIX expression was chosen for production.

For production the selected clone was incubated at a density of 0.3×10e6 cells/mL in PowerCHO-2CD median (LONZA) supplemented with 4mM Ultraglutamine-1 (Lonza) and incubated under shaking conditions at 37°C for 4 days, followed by 5 additional days at 31°C. The cell supernatant was then collected and clarified by centrifugation and filtration before loading it on a cOmplete™ His-Tag purification resin (Roche). Following a washing step with 250 mM NaCl, 10mM imidazole, the protein was eluted with 250 mM NaCl, 250mM imidazole and finally dialyzed against PBS. Protein characterization is described in **Figure S4**.

### Enzymatic assay

Carbonic anhydrase IX was diluted in assay buffer (12.5 mM Tris-HCl, 75 mM NaCl, 1% DMSO pH 7.5) to reach a final concentration of 200 nM. 10 μL of the respective compound was transferred into a transparent flat-bottom 384-well microplate (Greiner Bio-One, #781901) to perform a 1:1 serial dilution in the assay buffer with a typical concentration range of 40 μM - 20 pM. To each well, 20 μL of 200 nM CAIX and 10 μL of 1 mM 4-nitrophenylacetate (assay buffer, 3% acetone) was added. After an incubation period of 60 minutes at 37°C in the dark, the absorption was measured at 400 nm on a Tecan Spark® Multimode Microplate Reader. Values were normalized in respect to the enzyme activity in the absence of inhibitor. The assay was performed for 152 compounds in single titration experiments (see **Table S3**) between those, 14 most potent candidates were selected. The assay was then performed in duplicates on 39 capped analogs of the previous 14 HITs and between those 8 were selected for FITC derivatization for in vivo studies (see **Figure S5**).

### Synthesis

Chemical synthesis and compound characterization are described in details in Supporting Information: Additional Methods.

### Fluorescence polarization (FP) measurements

Fluorescence polarization measurements were performed in black 384-well microplates (Greiner Bio-One, #784900). A 1:1 dilution series of the protein (i.e., CAIX or serum albumin in PBS) was prepared to reach a final volume of 5 μL per well. Subsequently, 5 μL of the fluoresceinated compound (20 nM in PBS) was added to each well. The plate was centrifuged (400 rcf, 1 min) and incubated in the dark for 15 min at room temperature. Anisotropy was recorded on a Tecan Spark® Multimode Microplate Reader (Excitation = 485 ± 20 nm, Emission = 535 ± 25 nm). For all tested compounds, FP measurements were performed in five independent replicates (see **Figure S6-S13**).

### Cell culture

The human renal cell carcinoma cell line SK-RC-52 was kindly provided by Professor E. Oosterwijk (Radbound University Nijmegen Medical Centre, Nijmegen, The Netherlands). SK-RC-52 Cells were cultured in RPMI medium (Invitrogen), supplemented with fetal calf serum (10%, FCS, Invitrogen) and Antibiotic-Antimycotic (1%, AA, Invitrogen) at 37°C, 5% CO_2_. Once confluence was reached (90-100% confluence), tumor cells were detached using Trypsin-EDTA 0.05% (Invitrogen) and re-seeded at a dilution of 1:6. Expansion of tumor cells was continued until sufficient cells to run *in vitro* and *in vivo* assays presented below were available.

### Confocal fluorescence microscopy

SK-RC-52 cells were seeded into 4-well coverslip chamber plates (Sarstedt, Inc.) at a density of 10^4^ cells per well in RPMI medium (1 mL, Invitrogen) supplemented with 10% Fetal Bovine Serum (FBS, Thermofisher), Antibiotic-Antimytotic (Gibco), and 10 mM HEPES (VWR). Cells were allowed to grow overnight under standard culture conditions. The culture medium was replaced with fresh medium containing the suitable FITC-conjugated probes (100 nM) and Hoechst 33342 nuclear dye (Invitrogen, 1 μg/mL). Colonies were randomly selected and imaged 30 min after incubation on a SP8 confocal microscope equipped with an AOBS device (Leica Microsystems).

### Animal studies

All animal experiments were conducted in accordance with Swiss animal welfare laws and regulations under the license number ZH006/2021 granted by the Veterinäramt des Kantons Zürich.

### Subcutaneous tumor implantation

SK-RC-52 cells were grown to 80-100% confluence and detached with Trypsin-EDTA 0.05% (Invitrogen). Cells were washed once with Hank’s Balanced Salt Solution (HBSS, Thermo Fisher Scientific, pH 7.4), counted and re-suspended in HBSS. Aliquots of 5-10 × 10^6^ cells were resuspended in 150 μL of HBSS and injected subcutaneously in the right flank of female athymic BALB/c nu/nu mice (8-10 weeks of age, Janvier).

### Ex vivo fluorescence analysis

Mice bearing subcutaneous SK-RC-52 tumors were injected intravenously with fluorescein-linked compounds (30 nmol dissolved in 150 μL sterile PBS, pH 7.4 5% v/v DMSO). Animals were sacrificed by CO_2_ asphyxiation 1 hour after the intravenous injection. Tumors were excised, snap-frozen in OCT medium (Thermo Scientific) and stored at −80°C. Cryostat sections (10 μm) were cut and nuclei were stained with Fluorescence Mounting Medium (Dako Omnis, Agilent). Images were obtained using an Axioskop2 mot plus microscope (Zeiss) and analyzed by ImageJ 1.53 software.

## Results

### Model-based HIT identification and characterization of potency by enzymatic assay

**Figure 1** presents the chemical structures of three building block libraries, named DEL01, DEL02 and DEL03, which were screened against streptavidin beads (no target control, NTC) and against beads coated with CAIX. Selection results show a homogenous count distribution for the no protein control selections. DEL screening against CAIX led to the preferential enrichment of several different combinations of building blocks which are indicated by the arrows. All screening experiments were performed in duplicates and gave consistent and reproducible fingerprints. (see **Figure S1**). Screening data from NTC and from CAIX were used as input for machine learning procedures, as presented in **Figure S2** and **S3**.

From the raw sequencing counts, enrichment scores were computed to assign classification labels to each of the DEL compounds for model training. The Instance-level GCNN models were trained in a 5-fold cross-validation setup, with the average cross-validation metrics summarized in *Material and Methods*. The best models were used to select a list of 152 high-scored and diverse compounds from Enamine (108 compounds) and Mcule Instock (44 compounds), which were experimentally validated for CAIX binding in enzymatic assays. Out of the 152 tested compounds, 108 compounds (71%) achieved a better half maximal inhibitory concentration (IC_50_) than sulfamoylbenzoic acid (SABA, IC_50_ = 1.2 μM), while 43 compounds (28%) revealed an IC_50_ below 50 nM. Furthermore, our model led to the identification of 12 aromatic sulfonamide comprising compounds with higher potency in comparison to a highly potent reference, acetazolamide (AAZ, IC_50_ = 24 nM). Representative structures of the HITs are shown in **Figure 2**. The 152 structures and respective IC_50_ values are summarized in **Table S3**.

**Figure 2.**
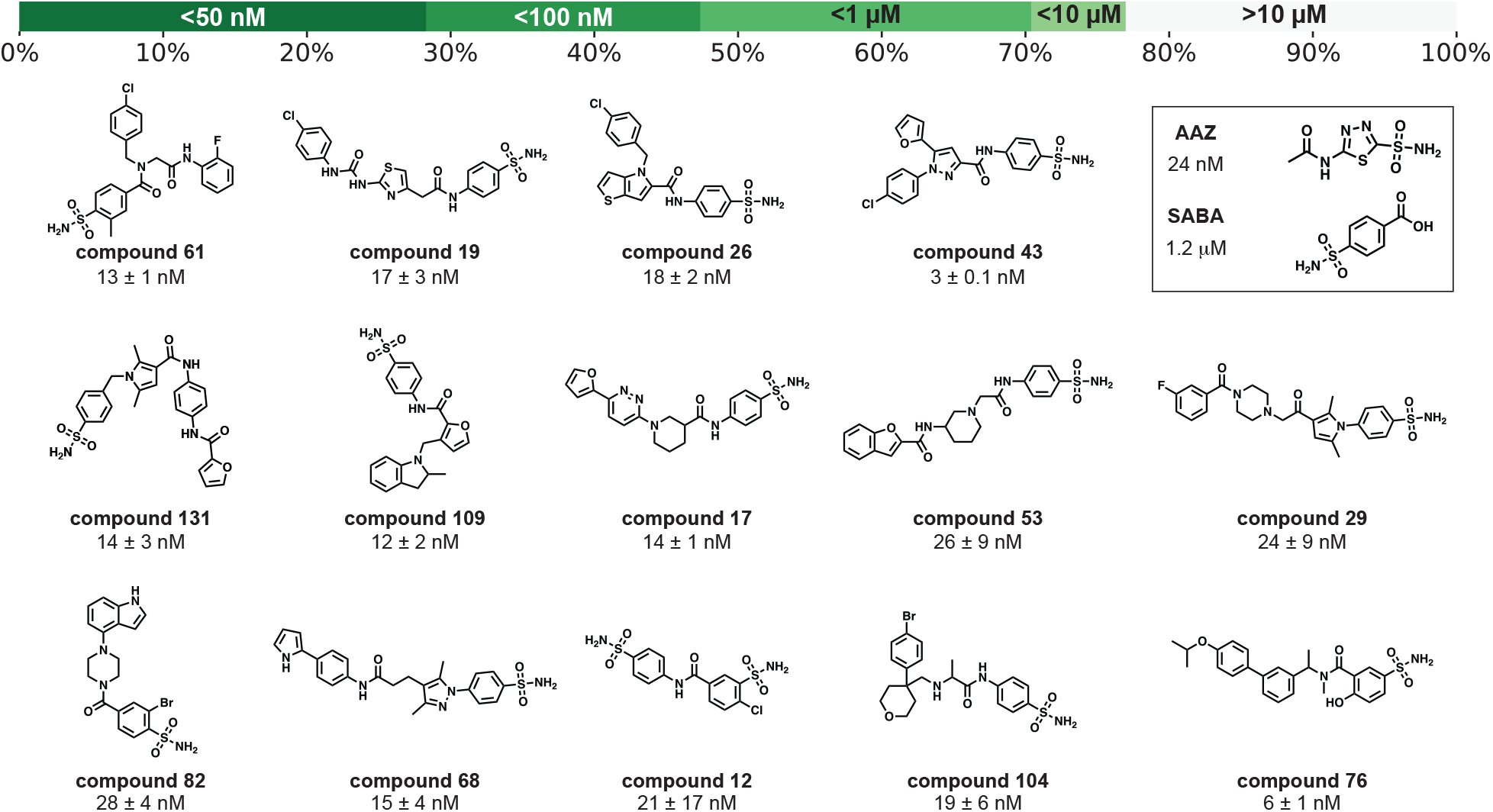
Representative structures of the identified HITs in HIT finding. The top panel shows the cumulative % hit rates of the 152 compounds that were identified with our approach and screened in an in vitro enzymatic inhibition assay, where the darker the color, the more stringent the potency cutoff. Out of the 152 compounds, 70% displayed submicromolar activities (IC_50_ < 1 μM) and 28% achieved IC_50_ < 50 nM. The GCNN model discovered 12 diverse and potent SABA-derived compounds that are more potent than AAZ (24 nM), improving the potency of SABA (1.2 μM) by 2-3 orders of magnitude.

### HIT optimization and LEAD generation

In order to develop tumor-targeting ligands against CAIX that can be functionalized with anti-cancer payloads, we aimed at identifying “portable” versions of the initial HITs displaying amine, alkyne or carboxylic acid functions (reaction handles for amidation and CuAAC click chemistry). Among the most potent HITs identified by our approach (IC_50_ < 50 nM, 28% of total HITs), we selected 14 compounds for further development based on the commercial availability of amine, alkyne and carboxylic acid analogs. A set of functionalized analogs were identified by fingerprint similarity (ECFP6 Tanimioto similarity ≥ 0.5) and by computationally installing amine, alkyne and carboxylic acid groups on sites distal from the sulfonamide group. (See Materials and Methods). A total of 46 analogs were prioritized by the GCNN model, purchased and experimentally validated by inhibition enzymatic assay. Analogs prioritized by the model retained the potency of the original starting points (67% of the surrogate compounds showed IC_50_ ≤ 50 nM). Mean pIC_50_ of the starting points and the surrogate compounds are 7.841 and 7.458, respectively. Mean LLE (pIC_50_ - clogP) of the starting points and the surrogate compounds are 4.7 and 5.0, respectively (**Figure S14**, **Table S4**).

Among the 46 portable analogs identified, eight analogs were selected and conjugated to fluorescein to enable affinity measurements by fluorescence polarization (FP) and further *in vivo* characterization. The fluorescein-conjugated compounds were screened against CAIX, human serum albumin and mouse serum albumin. All compounds bound to human recombinant CAIX in the nanomolar range (**Table S5**). Six of the tested compounds revealed higher affinity compared to acetazolamide [AAZ, K_D_ = (17.5 ± 1.4) nM] (**Figure 3**), a sulfonamide derivative which has already been successfully used for tumor-targeting applications in mice and in men.[50], [51] Moderate binding (K_D_ > 1 μM) was observed for human and mouse serum albumins.

**Figure 3.**
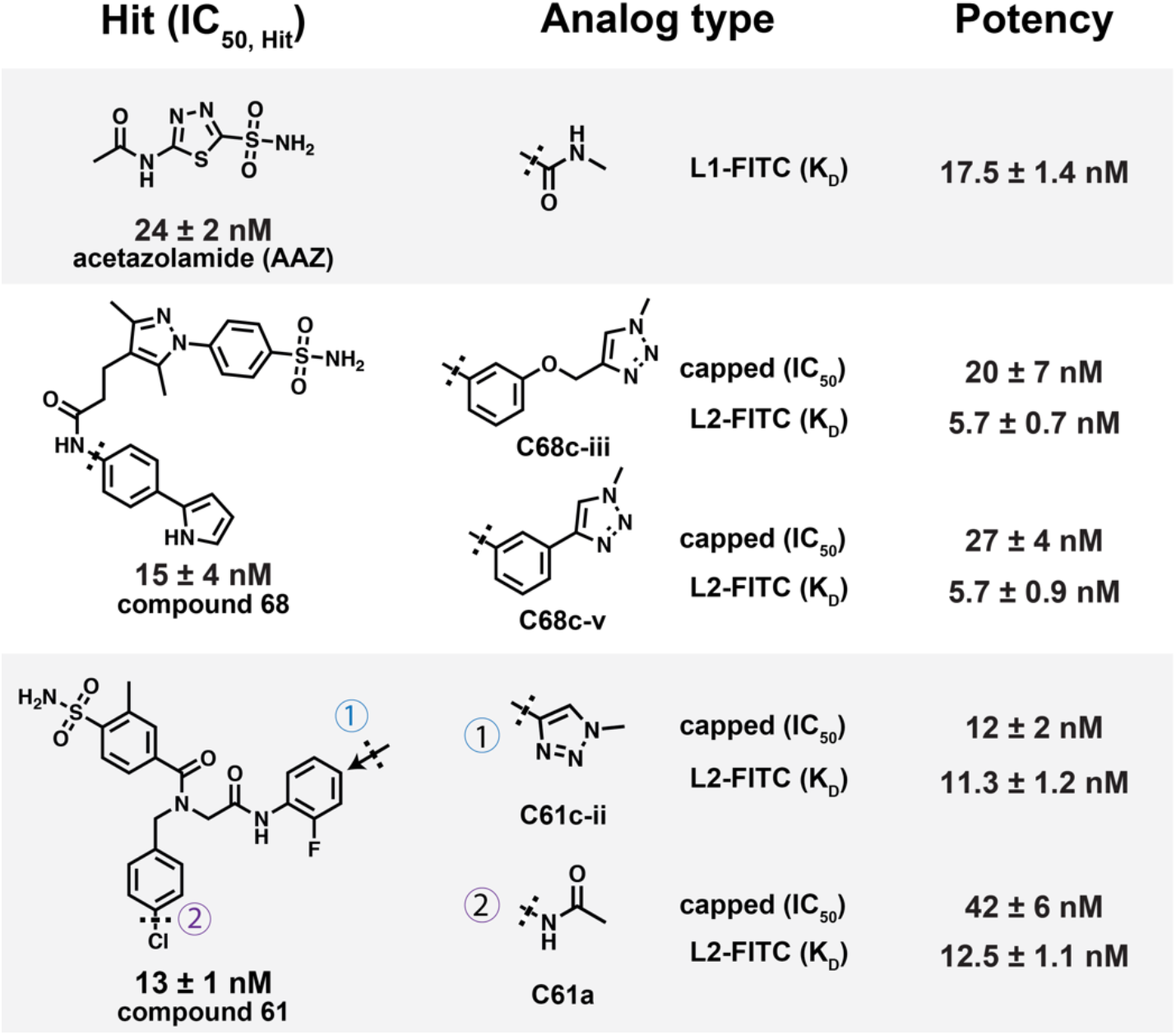
Potency of representative starting HITs, capped compounds and FITC conjugates. Starting HITs and capped analog compounds (surrogates) were validated in enzymatic assays (IC_50_). The confirmed analogs were subsequently conjugated with FITC and further validated with Fluorescence polarization (K_D_). Among the eight FITC conjugated compounds (data reported in **Table S5**), six compounds obtained better K_D_ than the positive control AAZ (K_D_ = 17.5 nM). L1 corresponds to the βAla-Asp-Lys tripeptide linker, while L2 corresponds to the PEG2 linker (see synthesis schemes presented in the **Supplementary Information**). Compound suffixes a, b, and c represent analog type amine, carboxylic acid and alkyne, respectively.

### Evaluation of similarities between DEL training set and ML-derived HITs and LEAD compounds

One of the advantages of applying machine learning for the valorization of DEL screening datasets is the ability to discover compounds outside of the original libraries, expanding the scope of DEL-derived chemical motifs. To inspect the ability of our model to identify HITs that are dissimilar to the DEL training set, we evaluated the nearest neighbor similarity of the diversity selected 152 HITs to the DEL training dataset (all training data and positive training examples (PTE) only, respectively) and computed their correlation to experimental potency. As shown in **Figure S15**, there is no meaningful correlation between similarity to the original DELs and experimental potency. Spearman correlation between experimental potency and similarity to the nearest DEL neighbor is 0.0994, while correlation to the nearest PTE is 0.1616. Furthermore, for the 46 tested surrogate compounds, Spearman correlation between experimental potency and similarity to the nearest DEL neighbor is −0.335, while correlation to the nearest PTE is −0.287.

### Cellular binding

Fluorescent derivatives of the compounds (compounds 61a, 68c-iii and 68c-v) selectively bound to CAIX on the surface of SK-RC-52 renal cell carcinoma cells in flow cytometry and confocal microscopy assays, while no binding was observed on the negative control cell line HEK-293 (CAIX negative cells) (**Figure 4A-C**, and **Figure S16**). AAZ* and untargeted fluorescein were included in the experiments as positive and negative controls, respectively.

**Figure 4.**
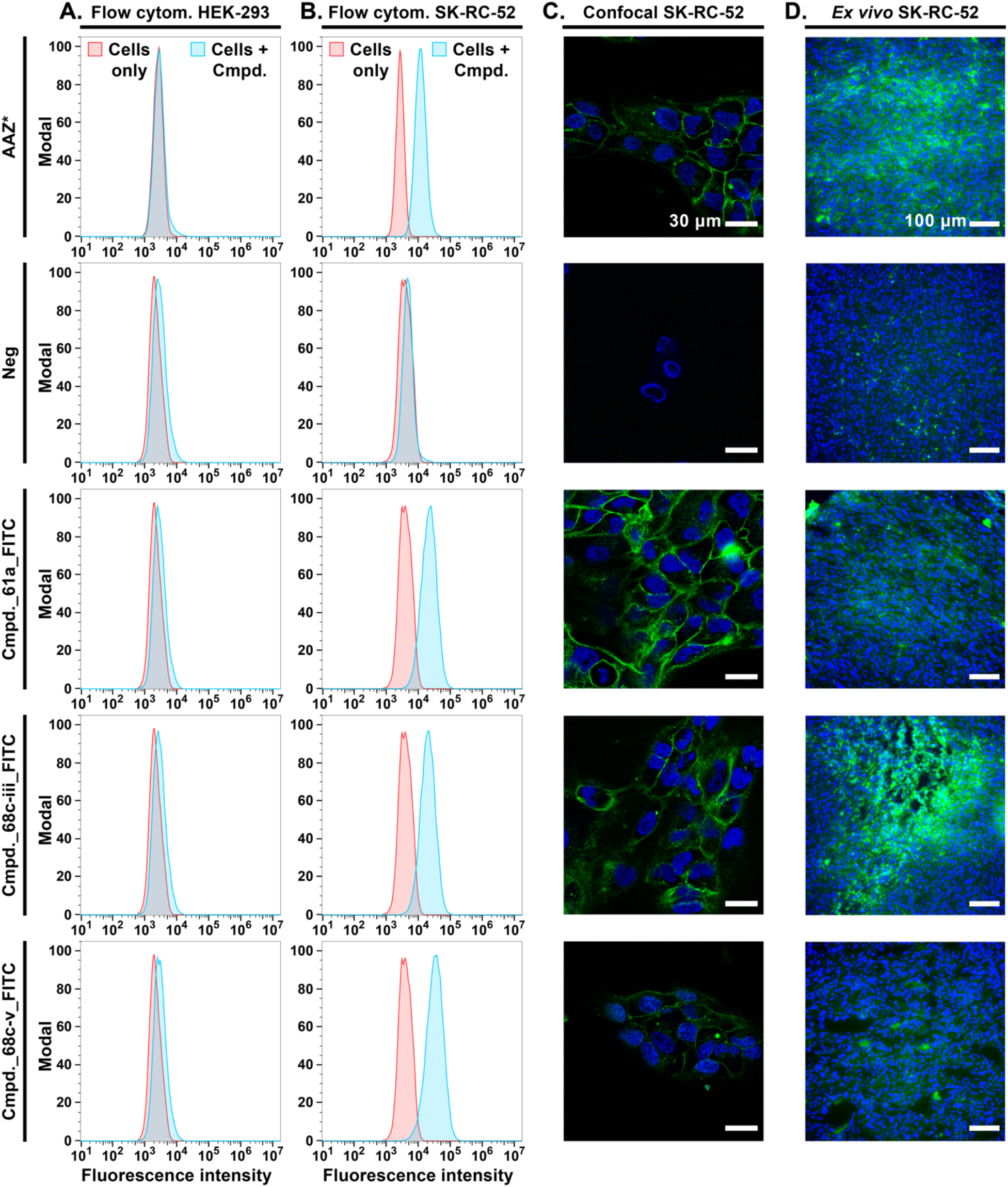
Cellular binding and ex-vivo biodistribution of CAIX-LEAD compounds. Flow cytometry analysis on (**A.**) CAIX-negative HEK-293 cells and (**B.**) CAIX-positive SK-RC-52 cells. **C.** Confocal fluorescence microscopy images on SK-RC-52 cancer cells. AAZ* and compounds 61a, 68c-iii and 68c-v accumulated on the surface of CAIX-positive SK-RC-52 cancer cells. No binding on CAIX-negative cells was observed in flow cytometry and confocal experiments. **D.** Results of ex vivo biodistribution experiments in SK-RC-52 tumor bearing mice. Microscopic pictures of cancer lesions collected 1-hour after systemic administration of compounds are presented. All compounds were injected intravenously (tail vein injection, dose of 30 nmol/mouse). Compound 68c-iii shows strong accumulation in CAIX positive tumors after systemic administration. Acetazolamide-Fluorescein (AAZ*) and a non-targeted fluorescein conjugate (Neg) were included in the in vitro and ex vivo experiments as positive and negative controls, respectively. GREEN = compounds (Fluorescein), BLUE = cancer cell nuclei (DAPI staining). Scale bar (confocal) = 30 μm. Scale bar (ex vivo biodistribution) = 100 μm.

### *Ex vivo* studies in tumor-bearing mice

In order to assess *in vivo* tumor targeting performance, the fluorescent derivatives compounds (compounds 61a, 68c-iii and 68c-v), AAZ* (positive control) and untargeted fluorescein (negative control) were intravenously administered to athymic balb/c nude mice bearing subcutaneous SK-RC-52 tumors. Microscopic analysis of the fluorescence signal associated to small molecules revealed a selective tumor accumulation of compound compound 68c-iii, similar to what was achieved with AAZ* (positive control). Untargeted fluorescein (negative control) and compounds (compounds 61a and 68c-v) did not accumulate to SK-RC-52 tumors (**Figure 4D**).

## Discussion

In this article, we presented the development of a novel drug discovery approach based on DEL screening data, processed and expanded by machine learning (ML). Our approach enabled the discovery of new HITs against CAIX for tumor targeting applications. During HIT finding and HIT-to-LEAD procedures, we applied our model on commercially available catalogs to discover novel and diverse HITs with structural features which fall outside of the original DEL chemical space. Our approach efficiently generated structurally diverse HITs, as demonstrated by *in vitro* biochemical characterization. No correlation between HIT potency and chemical similarity to the DEL screening training dataset was observed. Our method enabled successful translation of DEL selection data into LEAD compounds with promising *in vivo* biodistribution and excellent accumulation in CAIX-positive tumors.

Performance of machine learning models highly relies on the quantity and quality of the underlying training data. Compared to DELs, other screening methodologies included in the general term “high-throughput screening”[8], [19], [52] generate relatively small dataset which can hardly be used to train machine learning models. The drug discovery approach presented in this article can interrogate large libraries of commercially available compounds with limited costs and high productivity.

Even though multiplexed DEL screening experiments can generate large datasets covering broad chemical space, the low signal-to-noise ratios often hinder the ability of machine learning models to reliably capture the relationship between sequencing counts and affinity for the biological target. Synthon-based aggregations are often required to enhance the signals at the expense of discarding structural information. In this work, we performed selections and sequencing of individual libraries, achieving sampling ratios ~1000-fold higher in comparison to conventional multiplexed DEL screenings [26]. This approach led to high quality DEL datasets, which enabled the generation of a productive GCNN machine learning model.

The GCNN model presented was used to identify a panel of high affinity ligands of a tumor-associated antigen. As presented in **Figure 4**, additional hit-validation experiments that test the ability of the novel compounds to bind to the target in its natural cellular environment (e.g., antigen expressed on the surface of cancer cells) are still required to select LEAD candidates for *in vivo* applications. The availability of large datasets based on *in vitro* cellular binding could become crucial to further expand the scope of drug-discovery machine learning models and enhance their success rate.

To further expand from the instance-level classification model presented in this paper, it will be desirable to build probabilistic regression models, based on DEL screening results, to relate biological activity of compounds to sequencing reads. This approach may allow the *in silico* predictions of binding affinities [33]–[35]. Methods to accurately estimate synthetic yields during library construction, correct for PCR and HTS bias and normalize screening results across DELs could address inaccuracies which lower model productivity.

## Conclusions

We have developed a novel approach that combines DEL screening and instance-level deep learning modeling to identify tumor targeting agents against Carbonic Anhydrase IX (CAIX), a clinically validated marker of Renal Cell Carcinoma and hypoxia. The trained model enabled the discovery of diverse CAIX HITs which were not present in the original DEL chemical space. Furthermore, our method was applied to HIT-to-LEAD procedures to generate candidates that accumulated on the surface of CAIX-expressing tumor cells. The successful translation of LEAD candidates for *in vivo* tumor targeting applications demonstrates the potential of machine learning on DEL methods to advance real world drug discovery. The powerful discovery approach presented here can be generalized and will be applied to additional targets of pharmaceutical interest for the discovery of novel drug prototypes.

## Supporting Information

Additional experimental details for chemical synthesis and compound characterization, additional methods on compound selection, additional analysis, and experimental results for all tested compounds (DOCX). A Colab for running inference with the trained models is available at https://github.com/google-research/google-research/tree/master/gigamol.

## Acknowledgements

The authors would like to thank Amina Menhour for the help on the synthesis of HIT compounds, Marco Ruckstuhl for his work on the implementation of CAIX inhibition assays, Dr. Mattia Matasci for help with the protein production, Dr. Ettore Gilardoni for the MS-characterization of the presented compounds and Dr. Andrea Galbiati and Matilde Bocci for the assistance with *in vivo* experiments.

