## Supporting Information for "A deep learning approach for the discovery of tumor-targeting small organic ligands from DNA-Encoded Chemical Libraries"

Authors and affiliation:

Wen Torng*[1], Ilaria Biancofiore*[2], Sebastian Oehler*[2], Jin Xu[1], Jessica Xu[1], Ian Watson[1], Brenno Masina[2], Luca Prati[2], Nicholas Favalli[2], Gabriele Bassi[2], Dario Neri[2,3,4], Samuele Cazzamalli[#2], Jianwen A. Feng[#1]

[1]Google Research, 1600 Amphitheatre Parkway Mountain View, CA 94043

[2]Philochem AG, R&D Department, Otelfingen, 8112 Zürich, Switzerland

[3]Philogen S.p.A., Siena 53100, Italy

[4]Department of Chemistry and Applied Biosciences, Swiss Federal Institute of Technology (ETH Zürich), Zürich, Switzerland

*These authors contributed equally.

### Additional Methods

#### Chemical synthesis and compound characterization

##### Compound purification and characterization

Liquid Chromatography-Mass Spectrometry (LC-MS) spectra were measured on an Agilent 6100 Series Single Quadrupole MS system coupled with an Agilent 1200 Series LC, equipped with an InfinityLab Poroshell 120 EC-C18 Column (2.7 µm, 4.6 × 50 mm). Measurements were performed at a flow rate of 0.4 mL/min, with a linear gradient of eluent A (0.1% aqueous formic acid) and eluent B (acetonitrile with 0.1% formic acid): 0-10 min 90% - 0% A.

High-Resolution mass spectrometry (HR-MS) were performed on a Q Exactive Mass Spectrometer (Thermo Fisher Scientific) via direct injection at a flow rate of 4 µL/min. Recorded spectra had a resolution of 70000 FWHM (Full Width at Half Maximum) at 200 m/z.

Reversed Phase-Medium Performance Liquid Chromatography (RP-MPLC) was performed with a BÜCHI Sepacore Chromatography line (SepacoreRecord 1.4 software) coupled with a BÜCHI UV Photometer C-635 as a detector using a C18 40 µM irregular 12 g column (BUCHI, #145152103). Purifications were performed at a flow rate of 30 mL/min, with the following gradient of eluent A (0.1% aqueous formic acid) and eluent B (acetonitrile with 0.1% formic acid): 0-5 min 98% A, 5-45 min 98% to 0% A, 45-50 min 0% A, 50-50.1 min 0% to 98% A and 50.1-55 min 98% A.

Reversed Phase-High Performance Liquid Chromatography (RP-HPLC) was performed on an Agilent 1200 Series RP-HPLC which was connected to a PDA UV detector and equipped with a Synergi 4µm, Polar-RP 80Å 10 × 150 mm C18 column. The flow rate was set to 5 mL/min applying the following gradient of eluent A (0.1% aqueous trifluoroacetic acid) and eluent B (acetonitrile with 0.1% trifluoroacetic acid): 0-15 min 90% to 0% A, 15-16 min 0% A, 16-17 min 0% to 90% A, 17-18 min 90% A.

**Synthesis of intermediate 1 (negative control)**

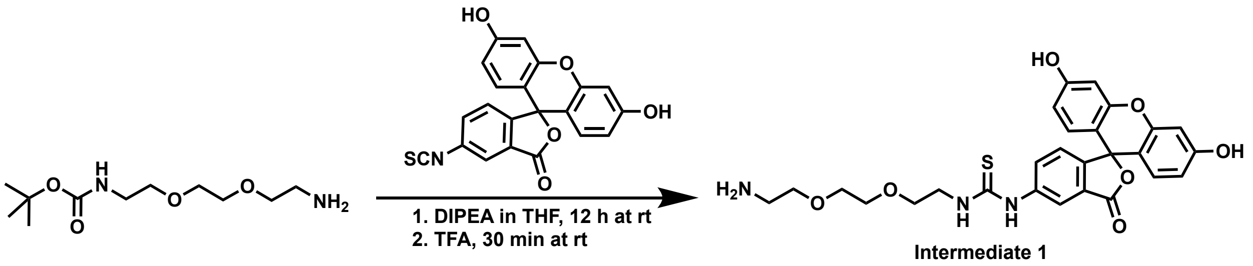


*Synthesis scheme of intermediate 1*

Tert-butyl *N*-[2-[2-(2-aminoethoxy)ethoxy]ethyl]carbamate (96 mg, 0.39 mmol, 1.5 eq) was dissolved in 15 mL Oxolan (THF) to add 3',6'-dihydroxy-6-isothiocyanatospiro[2-benzofuran-3,9'-xanthene]-1-one (FITC, 100 mg, 0.26, 1 eq.) and N-Ethyl-*N*-(propan-2-yl)propan-2-amine (DIPEA, 70 µL, 0.40 mmol, 1 eq.). After 12 h at room temperature, the solvent was removed under reduced pressure, the residue re-dissolved in 2 mL 2,2,2-Trifluoroacetic acid (TFA) to proceed with Boc-deprotection for 30 min. TFA was removed and the crude was dissolved in dichloromethane (DCM) for purification by direct-phase medium pressure liquid chromatography (MPLC). Intermediate 1 was obtained as yellow solid (94 mg, 47% yield). m/z calculated for $C_{27}H_{28}N_3O_7S$ $[M+H]^+$: 538.1642, detected (TOF MS ES+): 538.1622.

**Synthesis of intermediate 2**

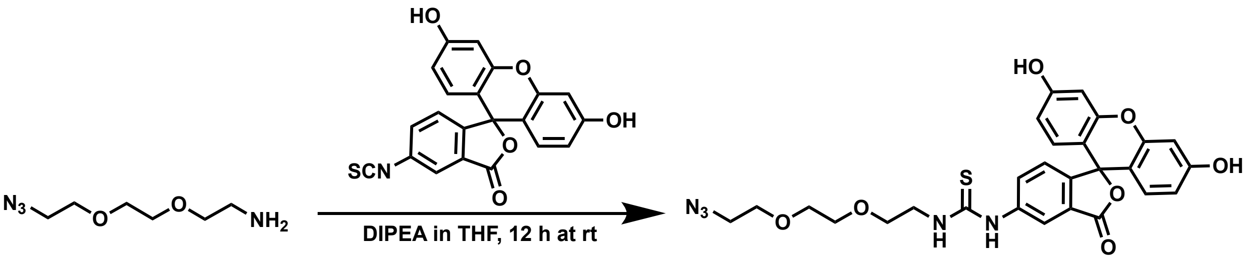


*Synthesis scheme of intermediate 2*

2-[2-(2-azidoethoxy)ethoxy]ethanamine (67 mg, 0.39 mmol, 1.5 eq.) was dissolved in 15 mL THF to add fluorescein-5-isothiocyanate (100 mg, 0.26, 1 eq.) and DIPEA (70 µL, 0.40 mmol, 1 eq.). After 12 h at room temperature, the solvent was removed under reduced pressure, the residue re-dissolved in water:acetonitrile 1:1 and purified by reversed phase (RP-) MPLC. Intermediate 2 was afforded as yellow solid (82 mg, 57% yield). m/z calculated for $C_{27}H_{26}N_5O_7S$ $[M+H]^+$: 564.1547, detected (TOF MS ES+): 564.1553.

**Synthesis of intermediate 3**

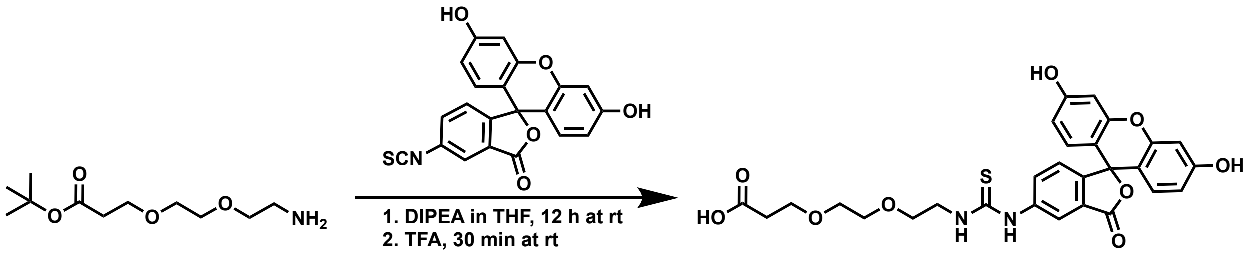


*Synthesis scheme of intermediate 3*

Tert-butyl 3-[2-(2-aminoethoxy)ethoxy]propanoate (90 mg, 0.39 mmol, 1.5 eq.) was dissolved in 15 mL THF. The reaction was initiated by addition of fluorescein-5-isothiocyanate (100 mg, 0.26, 1 eq.) and DIPEA (70 µL, 0.40 mmol, 1 eq.) and proceeded for 12 h at room temperature in the dark. Subsequently, solvent was removed under reduced pressure to perform Boc-deprotection in 2 mL TFA for 30 min at room temperature. TFA was evaporated and the residue re-dissolved in water:acetonitrile 1:1 for purification via RP-MPLC. Intermediate 3 was obtained as yellow solid (70 mg, 32% yield). m/z calculated for $C_{28}H_{27}N_2O_9S$ $[M+H]^+$: 567.1432, detected (TOF MS ES+): 567.1444.

**Synthesis of compound_26c_FITC**

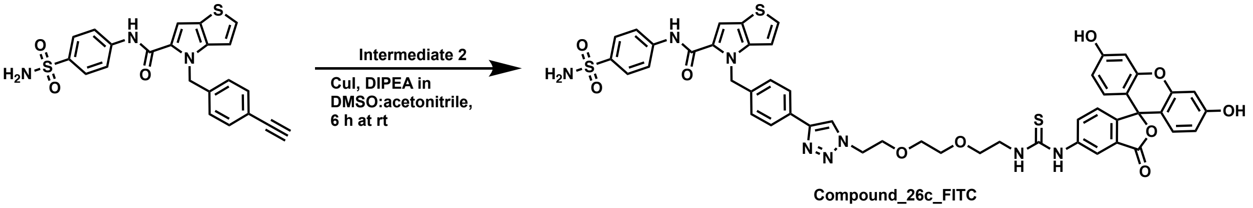


*Synthesis scheme of compound_26c_FITC*

4-[(4-ethynylphenyl)methyl]-*N*-(4-sulfamoylphenyl)-4*H*-thieno[3,2-b]pyrrole-5-carboxamide (4.7 mg, 11 µmol, 1 eq.) was dissolved in 500 µL dimethyl sulfoxide (DMSO):acetonitrile (1:1). DIPEA (1.9 µL, 11 µmol, 1 eq.), copper(I) iodide (CuI, 0.41 mg, 2.2 µmol, 0.2 eq.) and intermediate 2 (6.1 mg, 11 µmol, 1 eq.) were added to the mixture. The reaction proceeded for 6 h at room temperature to directly purify the crude by RP-HPLC, affording compound_26c_FITC as yellow solid (4.2 mg, 39% yield). m/z calculated for $C_{49}H_{43}N_8O_{10}S_3$ $[M+H]^+$: 999.2259, detected (TOF MS ES+): 999.2253.

**Synthesis of compound_68c-v_FITC**

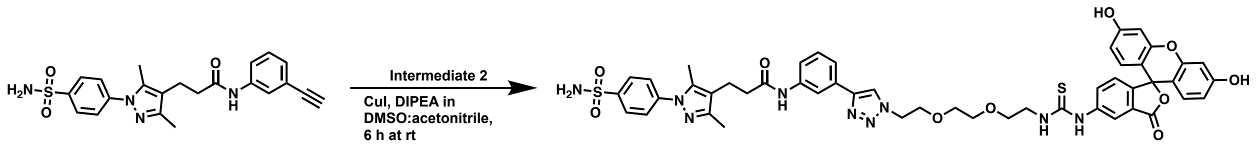

Compound_68c-v_FITC

*Synthesis scheme of compound_68c-v_FITC*

3-[3,5-dimethyl-1-(4-sulfamoylphenyl)-1*H*-pyrazol-4-yl]-*N*-(3-ethynylphenyl)propenamide (5.2 mg, 12 µmol, 1 eq.) was dissolved in 500 µL DMSO:acetonitrile (1:1). DIPEA (2.1 µL, 12 µmol, 1 eq.), CuI (0.47 mg, 2.5 µmol, 0.2 eq.) and intermediate 2 (6.9 mg, 12 µmol, 1 eq.) were added to the mixture. After 6 h at room temperature, the crude was purified by RP-HPLC, yielding compound_68c-v_FITC as yellow solid (3.1 mg, 25% yield). m/z calculated for $C_{49}H_{48}N_9O_{10}S_2$ $[M+H]^+$: 986.2960, detected (TOF MS ES+): 986.2976.

**Synthesis of compound_61c-ii_FITC**

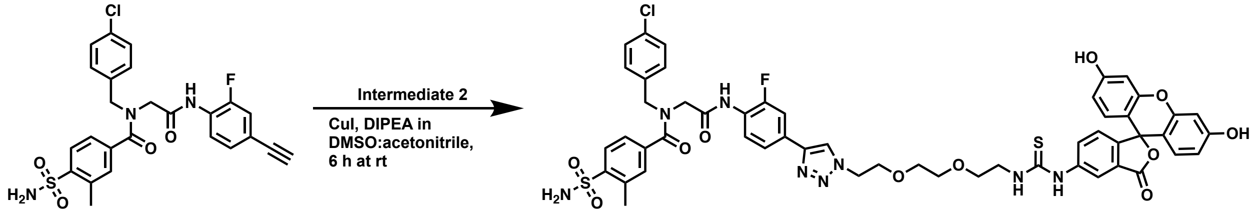

Compound_61c-ii_FITC

*Synthesis scheme of compound_61c-ii_FITC*

2-{*N*-[(4-chlorophenyl)methyl]-1-(3-methyl-4-sulfamoylphenyl)formamido}-*N*-(4-ethynyl-2-fluorophenyl)acetamide (5.0 mg, 10 µmol, 1 eq.) was dissolved in 500 µL DMSO:acetonitrile (1:1). DIPEA (1.7 µL, 10 µmol, 1 eq.), CuI (0.37 mg, 1.9 µmol, 0.2 eq.) and intermediate 2 (5.5 mg, 10 µmol, 1 eq.) were added to the mixture. The reaction proceeded for 6 h at room temperature followed by RP-HPLC purification resulting in compound_61c-ii_FITC as yellow solid (5.0 mg, 48% yield). m/z calculated for $C_{52}H_{47}ClFN_8O_{11}S_2$ $[M+H]^+$: 1077.2473, detected (TOF MS ES+): 1077.2473.

**Synthesis of compound_68c-iii_FITC**

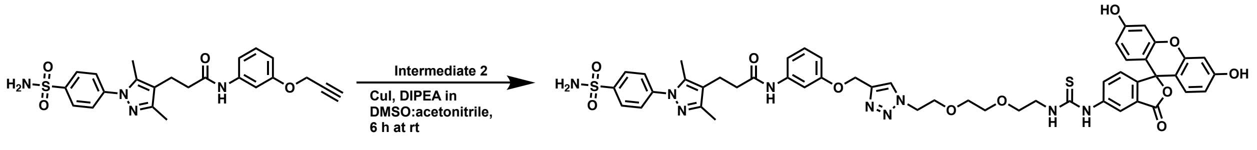

Compound_68c-iii_FITC

*Synthesis scheme of compound_68c-iii_FITC*

3-[3,5-dimethyl-1-(4-sulfamoylphenyl)-1*H*-pyrazol-4-yl]-*N*-[3-(prop-2-yn-1-yloxy)phenyl]propenamide (5.1 mg, 11 µmol, 1 eq.) was dissolved in 500 µL DMSO:acetonitrile (1:1). DIPEA (2.0 µL, 11 µmol, 1 eq.), CuI (0.43 mg, 2.2 µmol, 0.2 eq.) and intermediate 2 (6.4 mg, 11 µmol, 1 eq.) were added to the mixture. After 6 h at room temperature, the crude was purified by RP-HPLC, yielding compound_68c-iii_FITC as yellow solid (4.8 mg, 42% yield). m/z calculated for $C_{50}H_{50}N_9O_{11}S_2$ $[M+H]^+$: 1016.3066, detected (TOF MS ES+): 1016.3045.

**Synthesis of compound_43b_FITC**

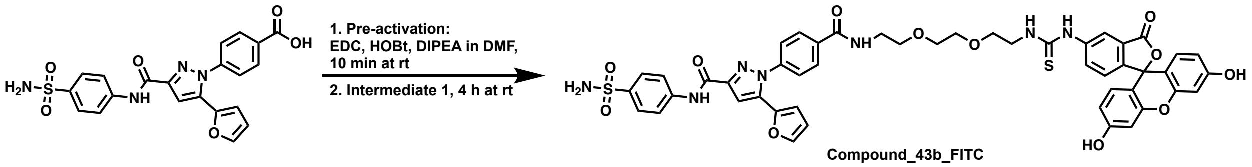


*Synthesis scheme of compound_43b_FITC*

4-[5-(furan-2-yl)-3-[(4-sulfamoylphenyl)carbamoyl]-1*H*-pyrazol-1-yl]benzoic acid (5.3 mg, 12 µmol, 1 eq.) was pre-activated with 1-ethyl-3-(3-dimethylaminopropyl)carbodiimide (EDC, 2 µL, 12 µmol, 1 eq.), 1-hydroxybenzotriazole (HOBt, 1.6 mg, 12 µmol, 1 eq.) and DIPEA (8.1 µL, 47 µmol, 4 eq.) in 500 µL *N*,*N*-dimethylformamide (DMF) for 10 min at room temperature. The pre-activation solution was added to intermediate 1 (6.3 mg, 12 µmol, 1 eq.) and the coupling proceeded for 4 h at room temperature. The reaction mixture was purified by RP-HPLC to obtain a yellow solid (compound_43b_FITC, 6.1 mg, 54% yield). m/z calculated for $C_{48}H_{42}N_7O_{12}S_2$ $[M+H]^+$: 972.2327, detected (TOF MS ES+): 972.2311.

**Synthesis of compound_53a_FITC**

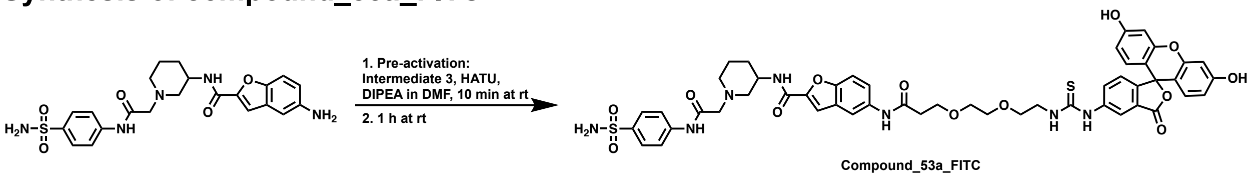


*Synthesis scheme of compound_53a_FITC*

Acid pre-activation of intermediate 3 (5.9 mg, 10 μmol, 1 eq.) was performed with *O*-(7-azabenzotriazol-1-yl)-*N*,*N*,*N'*,*N'*-tetramethyluronium hexafluorophosphate (HATU, 3.9 mg, 10 μmol, 1 eq.) and DIPEA (7.2 μL, 42 μmol, 4 eq.) in 500 μL DMF. Upon addition of 5-amino-*N*-(1-{[(4-sulfamoylphenyl)carbamoyl]methyl}piperidin-3-yl)-1-benzofuran-2-carboxamide (4.9 mg, 10 μmol, 1 eq.), the coupling proceeded for 1 h at room temperature. RP-HPLC purification yielded compound_53a_FITC as yellow solid (4.3 mg, 41% yield). m/z calculated for $C_{50}H_{50}N_7O_{13}S_2$ $[M+H]^+$: 1020.2903, detected (TOF MS ES+): 1020.2885.

**Synthesis of compound_61a_FITC**

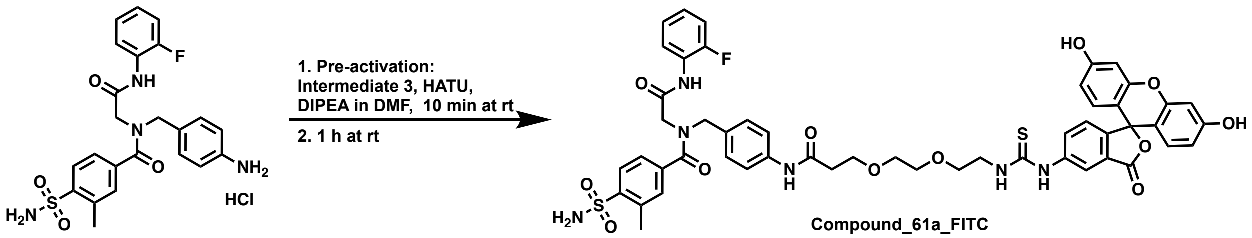


*Synthesis scheme of compound_61a_FITC*

Intermediate 3 (5.6 mg, 10 μmol, 1 eq.) was pre-activated with HATU (3.9 mg, 10 μmol, 1 eq.) and DIPEA (6.9 μL, 39 μmol, 4 eq.) in 500 μL DMF. 2-{*N*-[(4-aminophenyl)methyl]-1-(3-methyl-4-sulfamoylphenyl)formamido}-*N*-(2-fluorophenyl)acetamide hydrochloride (5.0 mg, 10 μmol, 1 eq.) was added and the reaction proceeded for 1 h at room temperature. RP-HPLC purification resulted in compound_61a_FITC as yellow solid (5.5 mg, 55% yield). m/z calculated for $C_{51}H_{48}FN_6O_{12}S_2$ $[M+H]^+$: 1019.2750, detected (TOF MS ES+): 1019.2733.

#### Synthesis of compound_109b-ii_FITC

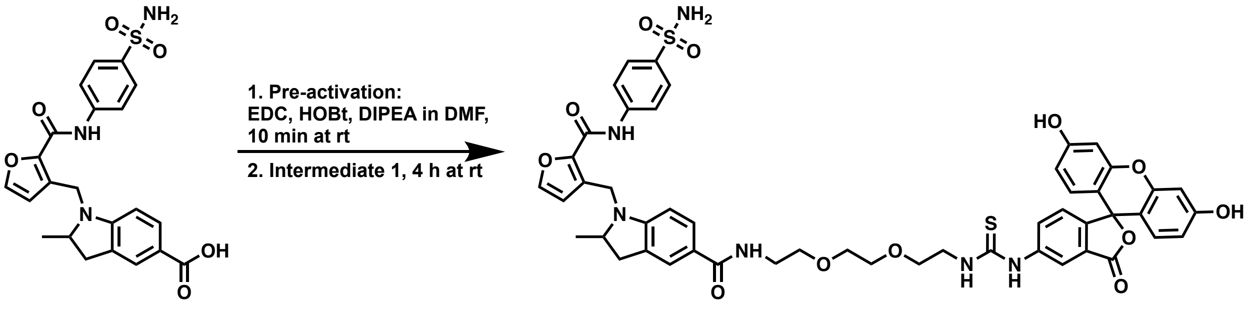


**Compound_109b-ii_FITC**

*Synthesis scheme of compound_109b-ii_FITC*

2-Methyl-1-({2-[(4-sulfamoylphenyl)carbamoyl]furan-3-yl}methyl)-2,3-dihydro-1*H*-indole-5-carboxylic acid (4.5 mg, 10 µmol, 1 eq.) was pre-activated with EDC (1.7 µL, 10 µmol, 1 eq.), HOBt (1.3 mg, 10 µmol, 1 eq.) and DIPEA (6.9 µL, 40 µmol, 4 eq.) in 500 µL DMF for 10 min at room temperature. Intermediate 1 was added to the reaction mixture (5.3 mg, 10 µmol, 1 eq.) and incubated for 4 h at room temperature. The crude was purified by RP-HPLC to obtain compound_109b-ii_FITC as a yellow solid (3.9 mg, 40% yield). m/z calculated for $C_{49}H_{47}N_6O_{12}S_2$ $[M+H]^+$: 975.2688, detected (TOF MS ES+): 975.2699.

#### Synthesis of AAZ*

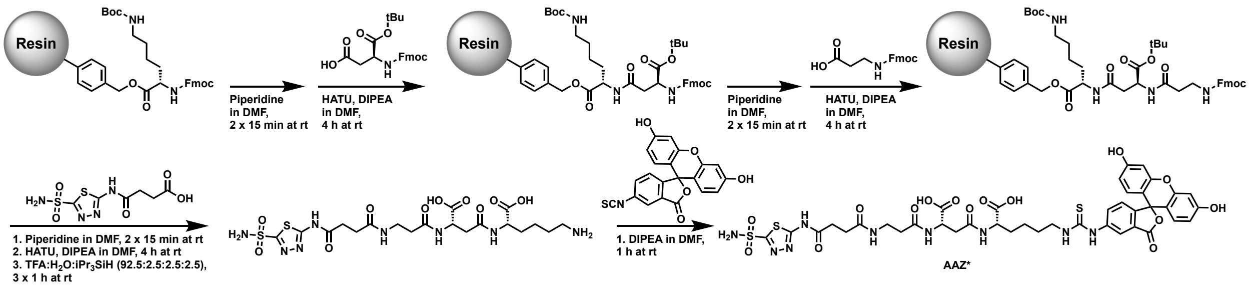


*Synthesis scheme of AAZ**

Preloaded Fmoc-Lys(Boc)-Wang resin (125 mg scale, 0.06 mmol loading capacity) was swollen in DMF. After Fmoc deprotection by two incubation cycles in DMF with 20% piperidine for 15 min at room temperature, the tripeptide Lys-Asp-βAla was generated by iterative amide coupling and Fmoc deprotection. Coupling were performed with 4 equivalents acid, 3.9 equivalents HATU and 8 equivalents DIPEA at following loading order: (2*S*)-2-(9*H*-fluoren-9-ylmethoxycarbonylamino)-6-[(2-methylpropan-2-yl)oxycarbonylamino]hexanoic acid (preloaded), (3*S*)-3-(9*H*-fluoren-9-

ylmethoxycarbonylamino)-4-[(2-methylpropan-2-yl)oxy]-4-oxobutanoic acid, 3-(9*H*-fluoren-9-ylmethoxycarbonylamino)propanoic acid. Subsequently, an acetazolamide derivative (4-oxo-4-[(5-sulfamoyl-1,3,4-thiadiazol-2-yl)amino]butanoic acid, 70 mg, 0.25 mmol, 4 eq.) in the presence of HATU (95 mg, 0.25 mmol, 4 eq.) and DIPEA (87 µL, 0.5 mmol, 8 eq.) in DMF. The coupling proceeded for 4 h at room temperature. The resin was washed with DMF and DCM and cleavage solution was added to the resin. The intermediate was purified by RP-HPLC and lyophilized overnight, yielding a white solid (3.7 mg, 10% yield). The intermediate was dissolved in 200 µL DMF to add 5-FITC (2.4 mg, 0.006 mmol, 1 eq.) and DIPEA (4.3 µL, 0.024 mmol, 4 eq.), followed by 1 h incubation at room temperature in the dark. The crude was directly purified by RP-HPLC and product was obtained after lyophilization as yellow solid (2.1 mg, 34% yield). m/z calculated for $C_{40}H_{42}N_9O_{15}S_3$ $[M+H]^+$: 984.1957, detected (TOF MS ES+): 984.1965.

### Supporting Information Tables

**Table S1. Sampling ratios of the three DEL libraries in CAIX screenings.** Sampling ratios of the three DEL libraries used in this work are computed with Eq S1.

$$Sampling\ ratio = \frac{total\ number\ of\ reads}{total\ diversity} \qquad \text{Eq S1}$$[1]

| Library | Number of compounds | Total sequencing reads (n=2) | Sampling ratio (n=2) |
|---|---|---|---|
| DEL01 | 4266000 | 2837745 / 2071139 | 0.67 / 0.49 |
| DEL02 | 1575168 | 1964585 / 2159325 | 1.25 / 1.37 |
| DEL03 | 53326 | 2669343 / 2533971 | 50.06 / 47.52 |

**Table S2. Pre-defined property filters applied to remove compounds that are non-drug-like or reactive.**

| Description | SMARTS |
|---|---|
| R1 Reactive alkyl halides | [Br,Cl,I][CX4;CH,CH2] |
| R3 Carbazides | O=CN=[N+]=[N-] |
| R4 Sulphate esters | COS(=O)O[C,c] |
| R6 Acid anhydrides | C(=O)OC(=O) |
| R10 esters of HOBT | C(=O)Onnn |
| R11 Isocyanates & Isothiocyanates | N=C=[S,O] |
| R15 Aromatic azides | cN=[N+]=[N-] |
| R16 beta carbonyl quaternary Nitrogen | C(=O)C[N+,n+] |
| R17 acylhydrazide | [N;R0][N;R0]C(=O) |
| R18 Quaternary C, Cl, I, P or S | [C+,Cl+,I+,P+,S+] |
| R19 Phosphoranes | C=P |
| R21 Nitroso | [N&D2](=O) |
| R23 Carbodiimide | N=C=N |
| R25 Triacyloximes | C(=O)N(C(=O))OC(=O) |
| I3 Crown ethers | [O;R1][C;R1][C;R1][O;R1][C;R1][C;R1][O;R1] |
| I4 Disulphides | SS |
| I5 Thiols | [SH] |
| I7 2,4,5 trihydroxyphenyl | c([OH])c([OH])c([OH]) |
| I8 2,3,4 trihydroxyphenyl | c([OH])c([OH])cc([OH]) |
| I9 Hydrazothiourea | N=NC(=S)N |
| I11 Benzylic quaternary Nitrogen | cC[N+] |
| I12 Thioesters | C[O,S;R0][C;R0](=S) |
| I13 Cyanamides | N[CH2]C#N |
| I16 Betalactams | N1CCC1=O |
| N1 Quinones | O=C1[#6]~[#6]C(=O)[#6]~[#6]1 |
| N3 Saponin derivatives | O1CCCCC1OC2CCC3CCCCC3C2 |
| N5 Cycloheximide derivatives | O=C1CCCC(N1)=O |
| N6 Monensin derivatives | O1CCCCC1C2CCCO2 |
| imine | C=[N!R] |
| iodine | I |
| 2halo_pyrazine_3EWG | [#7;R1]1[#6]([F,Cl,Br,I])[#6]([$(S(=O)(=O)),$(C(F)(F)(F)),$(C#N),$(N(=O)(=O)),$([N+](=O)[O-]),$(C=O)])[#7][#6][#6]1 |
| 2halo_pyrazine_5EWG | [#7;R1]1[#6]([F,Cl,Br,I])[#6;!$(c-N)][#7][#6]([$(S(=O)(=O)),$(C(F)(F)(F)),$(C#N),$(N(=O)(=O)),$([N+](=O)[O-]),$(C=O)])[#6;!$(c-N)]1 |

| Description | SMARTS |
|---|---|
| 2halo_pyridazine_3EWG | [#7;R1]1[#6]([F,Cl,Br,I])[#6]([$(S(=O)(=O)),$(C(F)(F)(F)),$(C#N),$(N(=O)(=O)),$([N+](=O)[O-]),$(C=O)])[#6][#6][#7]1 |
| 2halo_pyridazine_5EWG | [#7;R1]1[#6]([F,Cl,Br,I])[#6][#6][#6]([$(S(=O)(=O)),$(C(F)(F)(F)),$(C#N),$(N(=O)(=O)),$([N+](=O)[O-]),$(C=O)])[#7]1 |
| 2halo_pyridine_3EWG | [#7;R1]1[#6;!$(c=O)]([F,Cl,Br,I])[#6]([$(S(=O)(=O)),$(C(F)(F)(F)),$(C#N),$(N(=O)(=O)),$([N+](=O)[O-]),$(C=O)])[#6;!$(c-N)][#6][#6;!$(c-N)]1 |
| 2halo_pyridine_5EWG | [#7;R1]1[#6;!$(c=O)]([F,Cl,Br,I])[#6][#6;!$(c-N)][#6]([$(S(=O)(=O)),$(C(F)(F)(F)),$(C#N),$(N(=O)(=O)),$([N+](=O)[O-]),$(C=O)])[#6;!$(c=O);!$(c-N)]1 |
| 2halo_pyrimidine_5EWG | [#7;R1]1[#6]([F,Cl,Br,I])[#7][#6][#6]([$(S(=O)(=O)),$(C(F)(F)(F)),$(C#N),$(N(=O)(=O)),$([N+](=O)[O-]),$(C=O)])[#6]1 |
| acrylate | [CH2]=[C;!$(C-N);!$(C-O)]C(=O) |
| activated_acetylene | [$(S(=O)(=O)),$(C(F)(F)(F)),$(C#N),$(N(=O)(=O)),$([N+](=O)[O-]),$(C(=O))]C#[C;!$(C-N);!$(C-n)] |
| activated_diazo | [N;!R]([$(S(=O)(=O)),$(C(F)(F)(F)),$(C#N),$(N(=O)(=O)),$([N+](=O)[O-]),$(C(=O))])=[N;!R]([$(S(=O)(=O)),$(C(F)(F)(F)),$(C#N),$(N(=O)(=O)),$([N+](=O)[O-]),$(C(=O))]) |
| activated_vinyl_ester | O=COC=[$(C(S(=O)(=O))),$(C(C(F)(F)(F))),$(C(C#N)),$(C(N(=O)(=O))),$(C([N+](=O)[O-])),$(C(C(=O)));!$(C(N))] |
| activated_vinyl_sulfonate | O(-S(=O)(=O))C=[$(C(S(=O)(=O))),$(C(C(F)(F)(F))),$(C(C#N)),$(C(N(=O)(=O))),$(C([N+](=O)[O-])),$(C(C(=O)));!$(C(N))] |
| acyl_activated_NO | O=C(-[!N])O[$([#7;+]),$(N(C=[O,S,N])(C=[O,S,N]))] |
| alpha_dicarbonyl | C(=O)!@C(=O) |
| alpha_halo_amine | [F,Cl,Br,I,$(O(S(=O)(=O)))]-[CH,CH2;!$(C(F)F)]-[N,n] |
| alpha_halo_heteroatom | [N,n,O,S;!$(S(=O)(=O))]-[CH,CH2;!$(C(F)(F))][F,Cl,Br,I,$(O(S(=O)(=O)))] |
| alpha_halo_heteroatom_tert | [N,n,O,S;!$(S(=O)(=O))]-C([Cl,Br,I,$(O(S(=O)(=O)))])(C)(C) |
| anhydride | [$(C(=O)),$(C(=S))]-[O,S]-[$(C(=O)),$(C(=S)),$(C(=[N;!R])),$(C(=N(-[C;X4])))] |
| aryl_phosphonate | P(=O)-[O;!R]-a |
| aryl_thiocarbonyl | a-[S;X2;!R]-[C;!R](=O) |
| azide | [$(N#[N+]-[N-]),$([N-]=[N+]=N)] |
| aziridine_diazirine | [C,N]1~[C,N]~N~1 |
| azo_amino | [N]=[N;!R]-[N] |
| azo_aryl | c[N;!R;!+]=[N;!R;!+]-c |
| azo_filter2 | [N;!$(N-S(=O)(=O));!$(N-C=O)]-[N;!r3;!$(N-S(=O)(=O));!$(N-C=O)]-[N;!$(N-S(=O)(=O));!$(N-C=O)] |
| beta_lactone | [#6,#15,#16]1(=O)~[#6]~[#6]~[#8,#16]1 |
| betalactam | C1(=O)~[#6]~[#6]N1 |

| Description | SMARTS |
| --- | --- |
| crown_ether | [$([O,S,#7;R1;r9,r10,r11,r12,r13,r14,r15,r16,r17,r18][CH,CH2;r9,r10,r11,r12,r13,r14,r15,r16,r17,r18][CH,CH2;r9,r10,r11,r12,r13,r14,r15,r16,r17,r18][O,S,#7;R1;r9,r10,r11,r12,r13,r14,r15,r16,r17,r18][CH,CH2;r9,r10,r11,r12,r13,r14,r15,r16,r17,r18][CH,CH2;r9,r10,r11,r12,r13,r14,r15,r16,r17,r18][O,S,#7;R1;r9,r10,r11,r12,r13,r14,r15,r16,r17,r18]),$([O,S,#7;R1;r9,r10,r11,r12,r13,r14,r15,r16,r17,r18][CH,CH2;r9,r10,r11,r12,r13,r14,r15,r16,r17,r18][CH,CH2;r9,r10,r11,r12,r13,r14,r15,r16,r17,r18][CH,CH2;r9,r10,r11,r12,r13,r14,r15,r16,r17,r18][O,S,#7;R1;r9,r10,r11,r12,r13,r14,r15,r16,r17,r18][CH,CH2;r9,r10,r11,r12,r13,r14,r15,r16,r17,r18][CH,CH2;r9,r10,r11,r12,r13,r14,r15,r16,r17,r18][CH,CH2;r9,r10,r11,r12,r13,r14,r15,r16,r17,r18][O,S,#7;R1;r9,r10,r11,r12,r13,r14,r15,r16,r17,r18]),$([O,S,#7;R1;r9,r10,r11,r12,r13,r14,r15,r16,r17,r18][CH,CH2;r9,r10,r11,r12,r13,r14,r15,r16,r17,r18][CH,CH2;r9,r10,r11,r12,r13,r14,r15,r16,r17,r18][O,S,#7;R1;r9,r10,r11,r12,r13,r14,r15,r16,r17,r18][CH,CH2;r9,r10,r11,r12,r13,r14,r15,r16,r17,r18][CH,CH2;r9,r10,r11,r12,r13,r14,r15,r16,r17,r18][CH,CH2;r9,r10,r11,r12,r13,r14,r15,r16,r17,r18][O,S,#7;R1;r9,r10,r11,r12,r13,r14,r15,r16,r17,r18])] |
| cyanohydrin | [C;X4](-[OH,NH1,NH2,SH])(-C#N) |
| diamino_sulfide | [N,n]~[S;!R;D2]~[N,n] |
| diazonium | a[N+]#N |
| flavanoid | O=C2CC(a3aaaaa3)Oa1aaaaa12 |
| gte_2_free_phos | P([O;D1])=O.P([O;D1])=O |
| gte_3_COOH | C(=O)[O;D1].C(=O)[O;D1].C(=O)[O;D1] |
| gte_3_iodine | [#53].[#53].[#53] |
| gte_4_basic_N | [N;!$(N(=[N,O,S,C]));!$(N(S(=O)(=O)));!$(N(C(F)(F)(F)));!$(N(C#N));!$(N(C(=O)));!$(N(C(=S)));!$(N(C(=N)));!$(N(#C));!$(Nc)].[N;!$(N(=[N,O,S,C]));!$(N(S(=O)(=O)));!$(N(C(F)(F)(F)));!$(N(C#N));!$(N(C(=O)));!$(N(C(=S)));!$(N(C(=N)));!$(N(#C));!$(Nc)].[N;!$(N(=[N,O,S,C]));!$(N(S(=O)(=O)));!$(N(C(F)(F)(F)));!$(N(C#N));!$(N(C(=O)));!$(N(C(=S)));!$(N(C(=N)));!$(N(#C));!$(Nc)].[N;!$(N(=[N,O,S,C]));!$(N(S(=O)(=O)));!$(N(C(F)(F)(F)));!$(N(C#N));!$(N(C(=O)));!$(N(C(=S)));!$(N(C(=N)));!$(N(#C));!$(N-c)] |
| gte_4_nitro | [$([N+](=O)[O-]),$(N(=O)=O)].[$([N+](=O)[O-]),$(N(=O)=O)].[$([N+](=O)[O-]),$(N(=O)=O)].[$([N+](=O)[O-]),$(N(=O)=O)] |
| gte_5_phenolic_OH | a[O;D1].a[O;D1].a[O;D1].a[O;D1].a[O;D1] |
| gte_7_aliphatic_OH | C[O;D1].C[O;D1].C[O;D1].C[O;D1].C[O;D1].C[O;D1].C[O;D1] |
| gte_7_total_hal | [Cl,Br,I].[Cl,Br,I].[Cl,Br,I].[Cl,Br,I].[Cl,Br,I].[Cl,Br,I].[Cl,Br,I] |

| Description | SMARTS |
| --- | --- |
| gte_8_CF2_or_CH2 | [CH2,$(C(F)(F));R0][CH2,$(C(F)(F));R0][CH2,$(C(F)(F));R0][CH2,$(C(F)(F));R0][CH2,$(C(F)(F));R0][CH2,$(C(F)(F));R0][CH2,$(C(F)(F));R0][CH2,$(C(F)(F));R0] |
| halo_5heterocycle_bis_EWG | [#7,#8,#16]1[#6]([$(S(=O)(=O)),$([F,Cl]),$(C(F)(F)(F)),$(C#N),$(N(=O)(=O)),$([N+](=O)[O-]),$(C(=O))])[#6]([$(S(=O)(=O)),$([F,Cl]),$(C(F)(F)(F)),$(C#N),$(N(=O)(=O)),$([N+](=O)[O-]),$(C(=O))])[#7][#6]1([Cl,Br,I]) |
| halogen_heteroatom | [!C;!c;!H][F,Cl,Br,I] |
| hydrazine | [N;X3;!$(N-S(=O)(=O));!$(N-C(F)(F)(F));!$(N-C#N);!$(N-C(=O));!$(N-C(=S));!$(N-C(=N))]-[N;X3;!$(N-S(=O)(=O));!$(N-C(F)(F)(F));!$(N-C#N);!$(N-C(=O));!$(N-C(=S));!$(N-C(=N))] |
| hyperval_sulfur | [$([#16&D3]),$([#16&D4])]=,:[#6] |
| keto_def_heterocycle | [$(c([C;!R;!$(C-[N,O,S]);!$(C-[H])](=O))1naaaa1),$(c([C;!R;!$(C-[N,O,S]);!$(C-[H])](=O))1naa[n,s,o]1)] |
| maleimide_etc | [$([C;H1]),$(C(-[F,Cl,Br,I]))]1=[$([C;H1]),$(C(-[F,Cl,Br,I]))]C(=O)[N,O,S]C(=O)1 |
| meldrums_acid_deriv | O=C1OC(C)(C)OC(C1)=O |
| non_ring_CH2O_acetal | [O,N,S;!$(S~O)]!@[CH2]!@[O,S,N;!$(S~O)] |
| non_ring_acetal | [O,N,S;!$(S~O)]!@[C;H1;X4]!@[O,N,S;!$(S~O)] |
| non_ring_ketal | [O,N,S;!$(S~O)]!@[C;H0;X4](!@[O,N,S;!$(S~O)])(C) |
| oxonium | [o+,O+] |
| peroxide | [#8]~[#8] |
| phosphite | [c,C]-[P;v3] |
| phosphorous_nitrogen_bond | [#15]~[N,n] |
| phosphorus_phosphorus_bond | P~P |
| phosphorus_sulfur_bond | P~S |
| polyene | C=[C;!R][C;!R]=[C;!R][C;!R]=[C;!R] |
| polyhalo_phenol_a | c1c([O;D1])c(-[Cl,Br,I])c(-[Cl,Br,I])cc1.c1c([O;D1])c(-[Cl,Br,I])c(-[Cl,Br,I])cc1 |
| polyhalo_phenol_b | c1c([O;D1])c(-[Cl,Br,I])cc(-[Cl,Br,I])c1.c1c([O;D1])c(-[Cl,Br,I])cc(-[Cl,Br,I])c1 |
| polyhalo_phenol_c | c1c([O;D1])ccc(-[Cl,Br,I])c(-[Cl,Br,I])1.c1c([O;D1])ccc(-[Cl,Br,I])c(-[Cl,Br,I])1 |
| polyhalo_phenol_d | c(-[Cl,Br,I])1c([O;D1])c(-[Cl,Br,I])ccc1.c(-[Cl,Br,I])1c([O;D1])c(-[Cl,Br,I])ccc1 |
| polyhalo_phenol_e | c1c([O;D1])ccc(-[Cl,Br,I])c(-[Cl,Br,I])1.c1c([O;D1])ccc(-[Cl,Br,I])c(-[Cl,Br,I])1 |
| porphyrin | [#6;r16,r17,r18]~[#6]1~[#6]~[#6]~[#6](~[#6])~[#7]1 |
| quat_N_N | [N,n;R;+]!@[N,n] |

| Description | SMARTS |
| --- | --- |
| quat_N_acyl | [N,n;+]!@C(=O) |
| quinone_methide | [#6;!$([#6](-[N,O,S]))]1=[#6;!$([#6](-[N,O,S]))][#6](=[#6])[#6;!$([#6](-[N,O,S]))]=[#6;!$([#6](-[N,O,S]))][#6]1(=[O,N,S]) |
| sulf_D2_nitrogen | [S;D2](-[N;!$(N(=C));!$(N(-S(=O)(=O)));!$(N(-C(=O)))]) |
| sulf_D2_oxygen_D2 | [S;D2][O;D2] |
| sulfite_sulfate_ester | [C,c]OS(=O)O[C,c] |
| sulfonyl_heteroatom | [!#6;!#1;!#11;!#19]O(S(=O)(=O)(-[C,c])) |
| thio_hydroxamate | [S;D2]([$(N(=C)),$(N(-S(=O)(=O))),$(N(-C(=O)))]) |
| thiocarbonate | SC(=O)[O,S] |
| thioester | [S;!R;H0]C(=[S,O;!R])([!O;!S;!N]) |
| thiopyrylium | c1[S,s;+]cccc1 |
| trinitro_aromatic | [$(a1aaa([$(N(=O)(=O)),$([N+](=O)[O-])])a([$(N(=O)(=O)),$([N+](=O)[O-])])a1([$(N(=O)(=O)),$([N+](=O)[O-])]))),$(a1aa([$(N(=O)(=O)),$([N+](=O)[O-])])a([$(N(=O)(=O)),$([N+](=O)[O-])])aa1([$(N(=O)(=O)),$([N+](=O)[O-])]))),$(a1a([$(N(=O)(=O)),$([N+](=O)[O-])])aa([$(N(=O)(=O)),$([N+](=O)[O-])])aa1([$(N(=O)(=O)),$([N+](=O)[O-])]))] |
| trisub_bis_act_olefin | [CH;!R;!$(C-N)]=C([$(S(=O)(=O)),$(C(F)(F)(F)),$(C#N),$(N(=O)(=O)),$([N+](=O)[O-]),$(C(=O))])([$(S(=O)(=O)),$(C(F)(F)(F)),$(C#N),$(N(=O)(=O)),$([N+](=O)[O-]),$(C(=O))]) |
| vinyl_carbonyl_EWG | [C;!R]([$(S(=O)(=O)),$(C(F)(F)(F)),$(C#N),$(N(=O)(=O)),$([N+](=O)[O-]),$(C=O)])([$(S(=O)(=O)),$(C(F)(F)(F)),$(C#N),$(N(=O)(=O)),$([N+](=O)[O-]),$(C=O)])=[C;!R]([C;!R](=O))([!$([#8]);!$([#7])]) |
| long chain hydrocarbon | [CD2;R0][CD2;R0][CD2;R0][CD2;R0][CD2;R0][CD2;R0] |
| Filter18_oxime_ester | [!#7,#1]C=NO[C,S,P](=O)* |
| acetal_alkyne | C[OD2H0][CD2H2][OD2H0]C#C |
| acid_halide | [S,C](=[O,S])[Cl,Br,I,F] |
| acyclic_NCN | [NR0][CR0X4][$([NR0]#C),$([NR0]=C)] |
| acyclic_NS | N(S)=[S,N,O] |
| acyl_cyanide | N#CC(=O)[A] |
| acylhydrazide | C[NR0][NR0]C(=O)A=C |
| allenic_alcohol | [CX3]=[CX3][OD1H1] |
| aldehyde | [C;H1](=O)[#6] |
| alkyl_halide | [Br,Cl,I][CX4H2] |

| Description | SMARTS |
|---|---|
| alkyl_phosphate | P(O[C,c])(O[C,c])(O[C,c])=O |
| sulfonic_acid | [A]S(~O)(~O)[OH] |
| alkyne_halogenated_or_Si | C#C[F,Cl,Br,I,Si] |
| allyl_halide | [Br,Cl,F,I][CH]C=C |
| allyl_halide | [Br,Cl,F,I]C=C=C |
| sulfonyl_halide | [#6][SD4](~O)(~O)[Cl,Br,F,I] |
| alpha_beta_ket_or_ald | [$([CX3]);!$(CC=[O,S,N]);!$(C[O,S,N])]=C-[$([CH]),$(C([CX4,c]))]=O |
| alpha_beta_unsat_ca | C=CC(=O)[OH] |
| alpha_halocarbonyl_or_alpha_haloalkene | [F,Cl,Br,I]CC=[O,C] |
| alpha_halo_amine_1 | N1C([Br,Cl,F,I])CC=C1 |
| alpha_halo_ket_or_ald | [Cl,Br,I][$([CX4][CH]=O),$([CX4]C(=O)[CX4,c])] |
| amide_alkyl_or_alkene | O=C([$([CX2](#C)),$([CX3]=[CX3])])[$([NH2]),$([NH][c,CX4]),$(N([c,CX4])[c,CX4])] |
| hemiaminal_ether | [CX4]N[CH2]O[CX4] |
| amine_alkene/alkyne | [NH2][$(C=C),$(CC#C)] |
| good_leaving_group_urea_1b | [CD3H0]=CNC(=O)NC=[CD2H1] |
| 5_3_aniline | c12cc1ccc2[NX3] |
| 3_5_aniline | c21cccc1c2[NX3] |
| arene_sulfonyl_imidazole | O=S(c1ccccc1)(n2ccnc2)=O |
| aryl_halide_specific | [I,Br,Cl,F][#7] |
| azide_halide | [Cl,Br,F,I]N=[N+]=[N-] |
| aziridine | C1C[N;!H0]1 |
| azocyanamides | [a,A][N;R0]=[N;R0]C#N |
| azo_pyrrazole | [C,N]1=[C,N][CD3H0]([C,N]=[C,N]1)=[CD1H2] |
| azo_pyrrazole | [CD1H2]=[CD3H0]1[CD2H2][ND2H0]=[ND2H0][CD2H2]1 |
| benzyl_ether_alkyl | c1ccccc1[CX2][O,OX1][c,C] |
| good_leaving_group_urea_1a | NC(=O)N=[CD3H0] |
| benzyloxycarbonyl_CBZ | c1ccccc1COC(=O)[N,n] |
| beta_azo_carbonyl | [a,A]C(=O)CN=N |
| beta_carbonyl_quat_nitrogen2 | C1[N+,n+]C(=O)C1 |
| beta_carbonyl_quat_nitrogen3 | C1[N,n;R1;D3]C(C1)=O |
| terminal_vinyl | [CH2]=[CH]-[Br,Cl,I,F] |
| cyctochalasin_derivs | c1ccccc1CC2[NH]C(=O)C3[CH]2CCC(O)C3 |
| cytochalaisn_dervs_2 | O=C1NCC2CCCCC21 |
| di_tri_phosphates | P(=O)([OH])OP(=O)[OH] |
| dichloramine | [NX3](Cl)Cl |
| disulfide | [S;D2]-[S;D2] |
| dithioacetal | [$([S;D2]([CX4,c])!@[CH,CH2]!@[S;D2][CX4,c])] |
| electrophiles_for_cyc | [$(N#CSc1sc(nc1)N),$([S,Se]1C(N)C(=O)[#6][#6]1)] |
| end_vinyl | [CH2]=[CH]-[N,O,S] |

| Description | SMARTS |
|---|---|
| enedione | [$([CX3]C(=O)[CX4,c]);!$(CC=[S,N]);!$(C[O,S,N])]=[$([CX3]C(=O)[CX4,c]);!$(CC=[S,N]);!$(C[N,S])] |
| enol_ether | [$([CX3]);!$(CC=[O,S,N]);!$(C[O,S,N])]=[$([CX3]O[CX4]);!$(CC=[O,S,N]);!$(C[N,S])] |
| epoxides_thioepoxides_aziridines | C1[O,S,N]C1 |
| halo_alkene | [Cl,Br,I]C=C |
| halo_phosphane | P[Cl,Br,F,I] |
| trimethylsilyl_tms | [CH3][#14;X4]([CH3])[CH3] |
| hemiacetal | [OH][CH,CH2]O[CX4,c] |
| hemiketal | O([#6])-C([#6])([#6])-[OH] |
| HOBT_esters | n1nn(OC(=O))c2ccccc12 |
| hydrazone | [N;R0][N;R0]=C |
| hydroxy_amine | [OH][$([NX3]([C;!$(C=[O,S,N])])[C;!$(C=[O,S,N])]),$([NH][CX4])] |
| imidoyl_halide | [Cl,Br,F,I]C=N |
| iodoso | [IX2]=O |
| iodoxy | O=I=O |
| isocyanate | [#6]N=C=O |
| isocyanide | C#N-[#6] |
| isonitrile | [N+]#[C-] |
| isothiocyanate | [#6]N=C=S |
| ketal | O([CX4,c])-C([CX4,c])([CX4,c])-O([CX4,c]) |
| sulfonic_ester | [#6]S(~O)(~O)O[#6] |
| labile_ester_2 | O=COn1cncc1 |
| labile_ester_3 | Fc1c(OC=O)c(F)c(F)c(F)c1F |
| lawesson_s_reagent | P(=S)(S)S |
| malonic_acid_cBu | C1(=O)CC(=O)C1 |
| michael_acceptor | [$(N#C-C=[CH][C,c]),$([CH](=[C;R0]-[CH]=O)),$([CH](=[C,R]-C(=O)-C));!$([CH]1=CC(=O)C=CC1=*);!$([CH]1=CC(=O)C(=[N,O])C=C1);!$([CH](=C-C=O)-C=O)] |
| michael_acceptor_enyne | [$(N#C-C#C[C,c]),$(C#C-[CH]=O),$(C(#C-C(=O)-[C,c]))] |
| thiocarbamate_1 | [C,c]OC(=S)N([C,c])([C,c]) |
| n_methoyl | OC[N+X4]C |
| n_oxide | [!$([C,c]-N(=O)~O);$([!O]~[N;R0]=O)] |
| n_p_s_halides | [S,P][Br,F,I,Cl] |
| nitroso | [$([A,a]);!$([O,o])][N&D2](=O) |
| thiocarbamate_2 | [C,c]SC(=O)N([C,c])([C,c]) |
| oxalyl_omega | CC(=O)C(=O)C |
| oxalyl_rlewis | OC(=O)C(=O)O |
| oxaziridine | C1NO1 |

| Description | SMARTS |
|---|---|
| oxime | C=[N;R0]-[OH] |
| paranitrophenyl_ester_1 | [O-][N+](=O)c1ccc(OC=O)cc1 |
| paranitrophenyl_ester_2 | ON(=O)c1ccc(OC=O)cc1 |
| paranitrophenyl_ester_3 | [O-][N+](=O)c1ccc(OC=O)cc1 |
| paranitrophenyl_esters | C(=O)Oc1ccc([$([N;X2]=O),$([N+;X2]=O)][O,O-])cc1 |
| thiocyanate | SC#N |
| thiolating_agent_q | [$([S,s]~[S,s]~[C,c]=S),$([S,s]~[C,c](=S)~[S,s,N]),$([S;D2;R0]-S~O)] |
| peroxide | O~O |
| phosphate | O=P(~O)(~O)(~O) |
| phosphinate | P(=O)(O[H,C])O[H,C] |
| phosphinic_acid | [PX4](=O)(C)(C)~[OH,O-] |
| phosphonamide | [PX4](=O)(N)(*)~[OH,O-] |
| phosphonic_acid | [PX3](=O)(~O)~[OH] |
| phosphonic_ester | [PX3](=O)(~O)~O-[#6] |
| phosphonylnitrile | [PX4](=O)(~O)(~O)C#N |
| phosphoramides | NP(=O)(N)N |
| phosphorane | C=[PD4] |
| phosphorane | [PX5] |
| phosphoric_acid | [PX4](=O)(~O)(~[OH,O-])~[OH,O-] |
| phosphoric_ester | [PX4](=O)(~O)(~O)~O-[#6] |
| phosphoryl | [PX4](=O)(~O)(~O)~O-[A,a] |
| phthalimides_pht | O=C2c1ccccc1C(=O)N2 |
| polyenes | C=CC=CC=CC=C |
| thiourea_good_LG | N(C(=S)N)-,=C#,=C |
| propiolactones | C1(=O)OCC1 |
| quinone | [c,C]1(~[O;D1])~*!-*~[c,C](~[O;D1])~*!-*~1 |
| tricarbo_phosphine | [C,c]P([C,c])[C,c] |
| squalestatin_derivatives | C12OCCC(O1)CC2 |
| sulfinimine | CS(=N)C |
| sulfonimine | [C,c]S(=N)(=O)[C,c] |
| sulfinylthio | CS(=O)SC |
| sulfonyl_cyanide | S(=O)(=O)C#N |
| sulfonamide_halogenated | [#6][SD4](~O)(~O)N[Cl,Br,I] |
| sulfone | [#6]([SD4](=O)(=O))[$([C;H1]=[C;H1]);$([C;H0]#[C;H0])] |
| dihydroquinoxalin-2-one | [#6]NC(=O)c1[c;H1][c;H1]c2N[C;H2]C(=O)[N;H1]c2[c;H1]1 |

**Table S3. Series of 152 commercially available compounds selected for initial characterization in single titration experiments.**

| Compound_id | Example_id | SMILES | $pIC_{50}$ | $IC_{50}$ Value [nM] |
|---|---|---|---|---|
| compound_1 | MCULE-1061907925 | CC1SC(C(NC2=CC=C(C(NC3=CC(S(N)(=O)=O)=CC=C3)=O)C=C2)=O)=CC=1 | 6.39 | 409 |
| compound_2 | MCULE-1463311986 | NS(C1=C(O)C=CC(NC(CCC2OC(C3=CC=C(Cl)C=C3)=NN=2)=O)=C1)(=O)=O | 7.52 | 30 |
| compound_3 | MCULE-1730923659 | C1(=CC2=CC=C(C=C2OC1=NC1=CC=C(C=C1)S(N)(=O)=O)O)C(=O)NC1C=CC=C(Br)C=1 | 4.52 | 30000 |
| compound_4 | MCULE-1881485455 | S(=O)(=O)(N1CCCCC1)C1C(C)=CC=C(C2=CC=C(N=N2)NC2C=CC(=CC=2)S(N)(=O)=O)C=1 | 8.10 | 8 |
| compound_5 | MCULE-1907142958 | S(N)(=O)(=O)C1=C(C)C=CC(=C1)C1C2=CC=CC=C2C(=NN=1)NC1=CC=C(C=C1)C(=O)N1CCCCC1 | 5.73 | 1866 |
| compound_6 | MCULE-2123354796 | NS(C1=CC=C(S(N2CCN(C(C3=CC=C(N4C5=C(C=CC=C5)N=C4)C=C3)=O)CC2)(=O)=O)C=C1)(=O)=O | 6.35 | 442 |
| compound_7 | MCULE-2281142794 | COC([C@H](C(C)(C)C)NC(C1=CC(F)=C(S(N)(=O)=O)C=C1)=O)=O | 7.35 | 45 |
| compound_8 | MCULE-2668444653 | O=C(C1OC=CC=1)N1CC2(CCCCC2)N(C(C2OC=CC=2)=O)CC1 | 4.52 | 30000 |
| compound_9 | MCULE-2692204760 | CC(C1=CC=C(S(N)(=O)=O)C=C1)NC(C1=C(NC(C2OC=CC=2)=O)C=CC=C1)=O | 7.30 | 50 |
| compound_10 | MCULE-2780956388 | C12=C(CCCC1C(=O)NC1C=CC(=CC=1)S(N)(=O)=O)SC(NC(=O)C1=CC=C(C=C1)Cl)=N2 | 6.77 | 168 |

| Compound_id | Example_id | SMILES | $pIC_{50}$ | $IC_{50}$ Value [nM] |
|---|---|---|---|---|
| compound_11 | MCULE-3102976174 | S(N)(=O)(=O)C1=CC=C(C=C1)CCNC(=S)NC1=CC=C(C=C1)S(N)(=O)=O | 7.82 | 15 |
| compound_12 | MCULE-3186254642 | NS(C1=CC=C(NC(C2=CC(S(N)(=O)=O)=C(Cl)C=C2)=O)C=C1)(=O)=O | 7.68 | 21 |
| compound_13 | MCULE-3388953576 | CC(C(NC1=CC=C(S(N)(=O)=O)C=C1)=O)SC1N(C2=CC=C(Cl)C=C2)C(=C2C3=C(C=CC=C3)N=C2)NN=1 | 6.77 | 169 |
| compound_14 | MCULE-3486122804 | CC(C(NC1=CC=C(S(N)(=O)=O)C=C1)=O)OC(C1CN(C(C2=CC=C(Cl)C=C2)=O)CCC1)=O | 7.37 | 43 |
| compound_15 | MCULE-3507440037 | N12C(CCC(=O)N3CCN(CC3)C(=O)C3=CC=CO3)=NN=C1C=CC(N1CCOCC1)=N2 | 4.52 | 30000 |
| compound_16 | MCULE-3642921701 | CC(C(COC(C=CC1=CC=C(NC2=CC(S(N)(=O)=O)=CC=C2)C=C1)=O)=O)(C)C | 6.82 | 150 |
| compound_17 | MCULE-3945027457 | S(N)(=O)(=O)C1C=CC(=CC=1)NC(=O)C1CCCN(C2=CC=C(N=N2)C2=CC=CO2)C1 | 7.85 | 14 |
| compound_18 | MCULE-4296129401 | S(N)(=O)(=O)C1C=CC(=CC=1)NC(=S)N1CCN(CC1)C(=O)C1=CC=CO1 | 7.43 | 37 |
| compound_19 | MCULE-4361519810 | S(N)(=O)(=O)C1=CC=C(C=C1)NC(=O)CC1=CSC(NC(=O)NC2C=CC(=CC=2)Cl)=N1 | 7.77 | 17 |

| Compound_id | Example_id | SMILES | $pIC_{50}$ | $IC_{50}$ Value [nM] |
|---|---|---|---|---|
| compound_20 | MCULE-4399748894 | NS(C1=CC=C(NC(NCC2=CC(NC(C3CC3)=O)=CC=C2)=O)C=C1)(=O)=O | 7.26 | 55 |
| compound_21 | MCULE-4907519609 | CC1=NN(C2=CC=C(C(N3CCC(C(NC4=CC=C(S(N)(=O)=O)C=C4)=O)CC3)=O)C=C2)C(C)=C1 | 7.43 | 37 |
| compound_22 | MCULE-5232776745 | S(N)(=O)(=O)C1C=CC(=CC=1)NC(=O)C(C)SC1=CC=C(C=C1)NC(=O)C1=CC=CC=C1C(O)=O | 7.51 | 31 |
| compound_23 | MCULE-5340873886 | NS(C1=CC=C(NC(COC(C2=C(NC3=CC(C(F)(F)F)=CC=C3)C=CC=C2)=O)=O)C=C1)(=O)=O | 7.06 | 87 |
| compound_24 | MCULE-5422132115 | C1(O)C=CC(CC2NC(SCC(=O)NC3C=CC(S(=O)(N)=O)=CC=3)=NC=2C)=CC=1 | 7.20 | 63 |
| compound_25 | MCULE-6609719606 | N1([C@@H](C(C)C)C(=O)NCC2=CC=C(C=C2)S(N)(=O)=O)CC2=C(C=CC=C2)C1=O | 7.01 | 97 |
| compound_26 | MCULE-6650112466 | N1(CC2=CC=C(C=C2)Cl)C2C=CSC=2C=C1C(=O)NC1=CC=C(C=C1)S(N)(=O)=O | 7.74 | 18 |
| compound_27 | MCULE-6750190981 | S(N)(=O)(=O)C1C=CC(=CC=1)NC(=O)C1C=C(NN=1)C1C(O)=CC(=C(C)C=1)C | 7.64 | 23 |
| compound_28 | MCULE-6956067696 | NS(C1=CC=C(NC(CCN2C(C(NC3=CC(Cl)=C(Cl)C=C3)=O)CCCC2)=O)C=C1)(=O)=O | 7.11 | 77 |
| compound_29 | MCULE-7420279083 | CC1N(C2=CC=C(S(N)(=O)=O)C=C2)C(C)=C(C(CN2CCN(C(C3=CC(F)=CC=C3)=O)CC2)=O)C=1 | 7.62 | 24 |

| Compound_id | Example_id | SMILES | $pIC_{50}$ | $IC_{50}$ Value [nM] |
|---|---|---|---|---|
| compound_30 | MCULE-7980450476 | NS(C1=CC(C(NC2=CC(NC(C3OC=CC=3)=O)=CC=C2)=O)=C(Br)C=C1)(=O)=O | 7.04 | 92 |
| compound_31 | MCULE-8021959114 | CC1N(CC2SC=CC=2)C(C)=C(C(COC(C2=CC(S(N)(=O)=O)=C(Cl)C=C2)=O)=O)C=1 | 6.36 | 437 |
| compound_32 | MCULE-8136696319 | NS(C1=CC(NC(COC(C2=CC=C(C3=CC=C(O)C=C3)C=C2)=O)=O)=CC=C1)(=O)=O | 6.59 | 256 |
| compound_33 | MCULE-8300814993 | CN(C(C1=CC=C(NC(C2=C(N3CCOCC3)C=CC(S(N)(=O)=O)=C2)=O)C=C1)=O)C | 7.08 | 84 |
| compound_34 | MCULE-8500392890 | N1(C2=CC=CC=C2)C(=O)C(N=C1SCC(=O)NC1=CC=C(C=C1)S(N)(=O)=O)=CC1=CC=CO1 | 6.75 | 176 |
| compound_35 | MCULE-8794960135 | NS(C1=CC=C(NC(COC(C2=C3C(C=CC=C3)=NC3=C2CCCC3=CC2OC=CC=2)=O)=O)C=C1)(=O)=O | 7.02 | 96 |
| compound_36 | MCULE-8816143072 | CC1=C(C(NC2=CC=C(NC(NCC3OC=CC=3)=O)C=C2)=O)C=C(S(N)(=O)=O)C=C1 | 7.01 | 97 |
| compound_37 | MCULE-8853006382 | NS(C1=CC(C(N2CCN(C(C3=CC(Cl)=CC=C3)=O)CC2)=O)=C(Cl)C=C1)(=O)=O | 6.44 | 360 |
| compound_38 | MCULE-8878007518 | NS(C1=CC=C(NC(COC(C(C2SC=CC=2)=CC2SC=CC=2)=O)=O)C=C1)(=O)=O | 6.49 | 322 |
| compound_39 | MCULE-8957726090 | CC(C(NC1=CC=C(S(N)(=O)=O)C=C1)=O)NC1=CC=C(NC2=CC=CC=C2)C=C1 | 7.40 | 40 |

| Compound_id | Example_id | SMILES | $pIC_{50}$ | $IC_{50}$ Value [nM] |
|---|---|---|---|---|
| compound_40 | MCULE-9096508877 | CC(C(NC1=CC=C(S(N)(=O)=O)C=C1)=O)OC(C=CC1OC=CC=1)=O | 7.40 | 40 |
| compound_41 | MCULE-9267754685 | CC(C(NC1=CC=C(S(N)(=O)=O)C=C1)=O)OC(CCC(C1=CC=C(N)C=C1)=O)=O | 7.82 | 15 |
| compound_42 | MCULE-9296368247 | NS(C1=CC=C(NC(C(NC(C2SC=CC=2)=O)CC2C3=C(C=CC=C3)NC=2)=O)C=C1)(=O)=O | 7.43 | 37 |
| compound_43 | MCULE-9438418188 | NS(C1=CC=C(NC(C2=NN(C3=CC=C(Cl)C=C3)C(C3OC=CC=3)=C2)=O)C=C1)(=O)=O | 8.52 | 3 |
| compound_44 | MCULE-9901244439 | N1C(=O)C(SC=1NC1C=CC(=CC=1)S(=O)(N)=O)CC(=O)NC1C=CC(=CC=1)Br | 7.28 | 52 |
| compound_45 | PV-001794340822 | O=C(NCC=1C=CC=C(NC(=O)C2=CC=CO2)C1)C3CCCCN3C(=O)C4=CC=CO4 | 4.52 | 30000 |
| compound_46 | PV-001820085764 | NS(=O)(=O)C=1C=CC(=CC1Cl)C(=O)NC=2C=CC(Cl)=CC2NC(=O)C3=CC=CO3 | 6.55 | 282 |
| compound_47 | PV-001830026270 | COC=1C=CC=2C=CC=CC2C1CN(C3CCCC3)C(=O)C=4C=C(C=CC4O)S(=O)(=O)N | 4.52 | 30000 |
| compound_48 | PV-001833025516 | CN(C1CCCN(C)C1)C(=O)C=2C=CC(NC(=O)C=3C=C(C=CC3O)S(=O)(=O)N)=CC2 | 4.52 | 30000 |
| compound_49 | PV-001840466155 | CC(C)(C)[C@H](NC(=O)C=1C=C(C=CC1O)S(=O)(=O)N)C(=O)OCC=2C=CC=CC2 | 4.52 | 30000 |

| Compound_id | Example_id | SMILES | pIC$_{50}$ | IC$_{50}$ Value [nM] |
|---|---|---|---|---|
| compound_50 | PV-001844387982 | CN(C(CNC(=O)OC(C)(C)C)C1CC1)C(=O)C=2C=CC(Cl)=C(C2)S(=O)(=O)N | 7.17 | 68 |
| compound_51 | PV-001852451390 | NS(=O)(=O)C=1C=CC(O)=C(C1)C(=O)N(CC2=CC=CO2)CC=3C=CC(O)=CC3 | 6.82 | 152 |
| compound_52 | PV-001918000390 | CC1CC=2C=C(C=CC2N1C(=O)CN3CCN(CC3)C(=O)C=4C=CC(O)=CC4)S(=O)(=O)N | 7.02 | 95 |
| compound_53 | PV-001918140173 | NS(=O)(=O)C=1C=CC(NC(=O)CN2CCCC(C2)NC(=O)C3=CC=4C=CC=CC4O3)=CC1 | 7.59 | 26 |
| compound_54 | PV-001961080300 | CC1=CC=C(S1)C(=O)N2CCN(CC2)C(=O)NCC=3C=CC=C(C3)C(=O)N4CCOCC4 | 4.52 | 30000 |
| compound_55 | PV-002087722760 | CC1CCC(CC1)NC(=O)CN2CCN(CC2)C(=O)NCC=3C=CC(=CC3C)S(=O)(=O)N | 5.62 | 2392 |
| compound_56 | PV-002096979560 | CC(N(C)C(=O)C=1C=CC(NC2=NS(=O)(=O)C=3C=CC=CC23)=CC1)C=4C=CC=C(C4)S(=O)(=O)N | 7.12 | 75 |
| compound_57 | PV-002102577649 | NS(=O)(=O)C=1C=CC(O)=C(C1)C(=O)NCC=2C=CN=C(C2)N3CCN(CC3)C=4C=CC(F)=CC4 | 4.52 | 30000 |
| compound_58 | PV-002105154988 | NS(=O)(=O)C=1C=CC(F)=C(CNC(=O)NC=2C=CC(=CC2)C(=O)N3CCCC3C=4C=CC=CC4)C1 | 4.91 | 12432 |
| compound_59 | PV-002112803468 | CC=1C=C(C=CC1S(=O)(=O)N)C(=O)N2CCN(CC3=NC(=NO3)C=4C=CC(Cl)=CC4)CC2 | 7.09 | 81 |

| Compound_id | Example_id | SMILES | $pIC_{50}$ | $IC_{50}$ Value [nM] |
|---|---|---|---|---|
| compound_60 | PV-002137030657 | CC1=NN(C(C)=C1C2COCCN2C(=O)C=3C=CC(Cl)=C(C3)S(=O)(=O)N)C=4C=CC=CC4 | 6.81 | 155 |
| compound_61 | PV-002143468768 | CC=1C=C(C=CC1S(=O)(=O)N)C(=O)N(CC(=O)NC=2C=CC=CC2F)CC=3C=CC(Cl)=CC3 | 7.89 | 13 |
| compound_62 | PV-002147702825 | CC(C)(C)OC(=O)N1CCN(C(C1)C=2C=CC=C(Cl)C2)C(=O)C=3C=CN=C(C3)S(=O)(=O)N | 7.30 | 50 |
| compound_63 | PV-002168728631 | CC(N(C)C(=O)C=1C=CC(=CC1)N2C=C(C=N2)C=3C=CC(Cl)=CC3)C=4C=CC(=CC4)S(=O)(=O)N | 7.21 | 62 |
| compound_64 | PV-002173372704 | CC=1C=NN(C1)C=2C=CC(=CC2NCC=3C=CC(=CC3)C(=O)N4CCCCC4)S(=O)(=O)N | 4.61 | 24573 |
| compound_65 | PV-002181194802 | NS(=O)(=O)C=1C=C(NC(=O)C=2OC=3C=CC=CC3C2CSC4=CC=CS4)C=CC1O | 4.52 | 30000 |
| compound_66 | PV-002187000791 | CCN1C(SCC=2C=C(Cl)C=C(C2)S(=O)(=O)N)=NC=3C=C(C=CC13)S(=O)(=O)N | 7.36 | 44 |
| compound_67 | PV-002190974130 | COC=1C=CC(CNC(=O)C=2N=NN(C2C)C=3C=CC(=CC3)C=4C=CC=CC4)=CC1S(=O)(=O)N | 4.52 | 30000 |
| compound_68 | PV-002194470557 | CC1=NN(C(C)=C1CCC(=O)NC=2C=CC(=CC2)C3=CC=CN3)C=4C=CC(=CC4)S(=O)(=O)N | 7.82 | 15 |
| compound_69 | PV-002200745773 | CC(C)[C@H](N1CC=2C=CC=CC2C1=O)C(=O)NC=3C=C(C=CC3O)S(=O)(=O)N | 5.55 | 2838 |

| Compound_id | Example_id | SMILES | $pIC_{50}$ | $IC_{50}$ Value [nM] |
|---|---|---|---|---|
| compound_70 | PV-002200991204 | COC=1C=CC(C(=O)N2CCCC(C2)C(=O)NC=3C=C(C=CC3O)S(=O)(=O)N)=C4C=CC=CC14 | 5.58 | 2610 |
| compound_71 | PV-002228990905 | CC=1C=CC(NC(=O)NC=2C=CC=C3CN(CCC23)C(=O)C=4C=CC(=C(C)C4)S(=O)(=O)N)=CC1 | 7.42 | 38 |
| compound_72 | PV-002257851860 | COC(=O)C1CCCCN1C(=O)C=2C=CC=C(NC(=O)CSC=3C=CC(=CC3)S(=O)(=O)N)C2 | 7.04 | 91 |
| compound_73 | PV-002258890910 | CC1=NN(C(C)=C1C(=O)NC=2C=CC(O)=C(C2)S(=O)(=O)N)C=3C=CC(Br)=CC3 | 4.52 | 30000 |
| compound_74 | PV-002267465517 | CC=1ON=C(C1C(=O)NC=2C=C(C=CC2O)S(=O)(=O)N)C=3C=CC(=CC3)C=4C=CC=CC4 | 4.52 | 30000 |
| compound_75 | PV-002287251748 | CC1CCCN(C1)C(=O)C=2C=CC(NC(=O)C3CN(C(=O)C3)C=4C=CC=C(C4)S(=O)(=O)N)=C(C)C2 | 4.52 | 30000 |
| compound_76 | PV-002288126501 | CC(C)OC=1C=CC(=CC1)C=2C=CC=C(C2)C(C)N(C)C(=O)C=3C=C(C=CC3O)S(=O)(=O)N | 8.22 | 6 |
| compound_77 | PV-002293358502 | CN(CC=1C=CC=CC1S(=O)(=O)N)C(=O)C=2C=CC(Cl)=CC2NC(=O)C=3C=COC3C | 4.52 | 30000 |
| compound_78 | PV-002294768980 | CC=1C=C(C=CC1S(=O)(=O)N)C(=O)N2CCCCC32CCN(C3)C(=O)OC(C)(C)C | 7.64 | 23 |
| compound_79 | PV-002301934044 | NS(=O)(=O)C=1C=CC(Br)=C(C1)C(=O)NC=2C=CC=C(NC(=O)C3CCCCC3)C2 | 7.28 | 53 |

| Compound_id | Example_id | SMILES | pIC$_{50}$ | IC$_{50}$ Value [nM] |
|---|---|---|---|---|
| compound_80 | PV-002346857595 | NS(=O)(=O)C=1C=C(NC(=O)C2CCC(CC2)NC(=O)CC3CCCC3)C=CC1O | 4.52 | 30000 |
| compound_81 | PV-002363796301 | CC=1C=CC(=CC1NC(=O)C2=CC=CO2)C(=O)N3CCC(CC3)C=4C=CC=C(C4)S(=O)(=O)N | 7.43 | 37 |
| compound_82 | PV-002378801870 | NS(=O)(=O)C=1C=CC(=CC1Br)C(=O)N2CCN(CC2)C=3C=CC=C4NC=CC34 | 7.55 | 28 |
| compound_83 | PV-002379117148 | CC=1C(NC(=O)C=2C=C(C=CC2O)S(=O)(=O)N)=CC=CC1C(=O)N3CCC=4SC=CC4C3 | 4.52 | 30000 |
| compound_84 | PV-002379570479 | COC(=O)C=1C=CC=2NC=C(C3CCN(CC3)C(=O)C=4C=C(C)C(C)=C(C4)S(=O)(=O)N)C2C1 | 6.82 | 152 |
| compound_85 | PV-002390375586 | CC=1C=C(C=CC1CNC(=O)NC=2C=CC(Cl)=C(C2)C(=O)NCC3=CC=CS3)S(=O)(=O)N | 7.07 | 86 |
| compound_86 | PV-002398818574 | NS(=O)(=O)C=1C=CC(=CC1Cl)C(=O)N2CCCC(CNC=3C=CC(Cl)=CC3)C2 | 6.69 | 206 |
| compound_87 | PV-002413919204 | CC(NC(=O)C(C)N1CCN(CC1)C(=O)C=2C=CC=C(O)C2)C=3C=CC(=CC3)S(=O)(=O)N | 6.12 | 750 |
| compound_88 | PV-002428685102 | NS(=O)(=O)C=1C=CC=C(NC(=O)CCN2CCC(CC2)C=3C=CC=C4NC=CC34)C1 | 4.96 | 10961 |
| compound_89 | PV-002439016629 | NS(=O)(=O)C=1C=CC=2N(CCC2C1)C(=O)CN3CCC(CC3)C(=O)NC=4C=C(Cl)C=CC4O | 4.52 | 30000 |

| Compound_id | Example_id | SMILES | $pIC_{50}$ | $IC_{50}$ Value [nM] |
|---|---|---|---|---|
| compound_90 | PV-002453933936 | NS(=O)(=O)C=1C=CC(=CC1[N+](=O)[O-])C(=O)N2CCC=3SC=CC3C2C=4C=CC(Cl)=CC4 | 7.34 | 46 |
| compound_91 | PV-002456829604 | CN(CC(=O)NC=1C=CC(=CC1)N2CCOCC2)C(=O)C=3C=CC(=C(Cl)C3)S(=O)(=O)N | 7.10 | 80 |
| compound_92 | PV-002458634382 | NS(=O)(=O)C=1C=C(CNC(=O)N(CC2CC2)CC=3C=CC(O)=CC3)C=CC1F | 5.92 | 1200 |
| compound_93 | PV-002459616211 | CN(CCN(C)C=1C=CC(=CC1)S(=O)(=O)N)C(=O)NC=2C=CC(=CC2C)C(=O)N3CCOCC3 | 4.52 | 30000 |
| compound_94 | PV-002465199505 | CC1=CC=C(CN(C(=O)C=2C=C(C=CC2O)S(=O)(=O)N)C=3C=CC=4OCCOC4C3)S1 | 6.35 | 442 |
| compound_95 | PV-002470670048 | NS(=O)(=O)C=1C=C(NC(=O)CC=2C=CC(NC(=O)C3=CC=C(Br)O3)=CC2)C=CC1F | 6.74 | 180 |
| compound_96 | PV-002515614235 | CN(CC1=CN(N=C1C2=CC=CS2)C=3C=CC=CC3)C(=O)C=4C=C(C=CC4O)S(=O)(=O)N | 7.00 | 100 |
| compound_97 | PV-002549663326 | CC1CCCCN1C(=O)C=2C=CC(NC(=O)C=3N=CC=C4C(=CC=CC34)S(=O)(=O)N)=C(C)C2 | 7.07 | 85 |
| compound_98 | PV-002563033473 | NS(=O)(=O)C=1C=C(C=C2CCCC21)C(=O)NCC=3C=CC(NC(=O)C=4C=CC=CC4O)=CC3 | 4.52 | 30000 |

| Compound_id | Example_id | SMILES | $pIC_{50}$ | $IC_{50}$ Value [nM] |
|---|---|---|---|---|
| compound_99 | PV-002568996599 | NS(=O)(=O)C=1C=C(C=CC1Cl)C(=O)NC=2C=CC(NC(=O)CCN3CCCCC3)=CC2 | 6.80 | 160 |
| compound_100 | PV-002575793063 | CC=1C(F)=CC(=CC1C(=O)N[C@@H](CC=2C=CC(O)=CC2)C(=O)OC(C)(C)C)S(=O)(=O)N | 7.46 | 35 |
| compound_101 | PV-002576867821 | CC(C)(C)OC(=O)NCC=1C=CC(=CC1)C=2C=CC=C(C2)C(=O)NC=3C=CC(=CC3)S(=O)(=O)N | 7.12 | 75 |
| compound_102 | PV-002584767299 | CC(C)C(NC(=O)CC=1C=CC=2C=CC=CC2C1)C(=O)NC=3C=CC(O)=C(C3)S(=O)(=O)N | 4.52 | 30000 |
| compound_103 | PV-002585030171 | CC=1C=CC(=CC1)N2C(C(CC2=O)C(=O)NC=3C=C(C=CC3O)S(=O)(=O)N)C4=CC=CS4 | 4.52 | 30000 |
| compound_104 | Z1003640198 | CC(NCC1(CCOCC1)C=2C=CC(Br)=CC2)C(=O)NC=3C=CC(=CC3)S(=O)(=O)N | 7.72 | 19 |
| compound_105 | Z1010591842 | CC(OC(=O)C=1C=C(CC=2C=CC=CC2)C=CC1O)C(=O)NC=3C=CC(=CC3)S(=O)(=O)N | 6.52 | 305 |
| compound_106 | Z1021718994 | NS(=O)(=O)C=1C=CC(CNC(=S)N(CC=2C=CC(=CC2)S(=O)(=O)N)CC(F)(F)F)=CC1 | 7.72 | 19 |
| compound_107 | Z1021812420 | CC=1C=CC(NC(=S)NCC=2C=CC(=CC2)S(=O)(=O)N)=CC1NC(=O)CN3CCCCC3 | 6.90 | 127 |
| compound_108 | Z1098824442 | NS(=O)(=O)C=1C=CC(NC(=O)CN(CC2=CC=CS2)C=3C=CC=C(C#C)C3)=CC1 | 7.02 | 95 |

| Compound_id | Example_id | SMILES | $pIC_{50}$ | $IC_{50}$ Value [nM] |
|---|---|---|---|---|
| compound_109 | Z1102054364 | CC1CC=2C=CC=CC2N1CC=3C=COC3C(=O)NC=4C=CC(=CC4)S(=O)(=O)N | 7.92 | 12 |
| compound_110 | Z1127103284 | CC(C(=O)NC=1C=CC(NC(=O)CC=2C=CC(=CC2)S(=O)(=O)N)=CC1Cl)N3N=C(C)C=C3C | 6.93 | 118 |
| compound_111 | Z1128050495 | NS(=O)(=O)C=1C=CC=C(CNC(=O)C=2C=CSC2NC(=O)C=3C=CC=4CCCCC4C3)C1 | 4.52 | 30000 |
| compound_112 | Z1128567557 | CCC=1C=CC(NC(=O)C2CSCN2C(=O)C=3C=CC(Cl)=CC3)=CC1S(=O)(=O)N | 4.52 | 30000 |
| compound_113 | Z1128797112 | NS(=O)(=O)C=1C=C(C=CC1F)C(=O)N2CCCCC2C(=O)NC=3C=C(Cl)C=CC3Cl | 7.52 | 30 |
| compound_114 | Z1132186624 | CCC(NC(=O)NC=1C=CC(NC(=O)C2CCCO2)=CC1)C=3C=CC=C(C3)S(=O)(=O)N | 6.21 | 615 |
| compound_115 | Z1213737360 | CC=1C=CC(C(=O)NCCC(=O)NCC=2C=C(C=CC2F)S(=O)(=O)N)=C(O)C1 | 5.78 | 1660 |
| compound_116 | Z1233687572 | CC=1C=CSC1C(=O)NC=2C=CC(OC(=O)C=3C=CC(=C(C)C3)S(=O)(=O)N)=CC2 | 6.62 | 238 |
| compound_117 | Z1270771584 | CC(C)CC(=O)NC=1C=CC(NC2CC(=O)N(C2=O)C=3C=CC=C(C3)S(=O)(=O)N)=CC1 | 7.39 | 41 |
| compound_118 | Z131023474 | CC1CCN(CC1)C(CNC(=O)NC=2C=CC(=CC2)S(=O)(=O)N)C3=CC=CS3 | 6.91 | 123 |

| Compound_id | Example_id | SMILES | $pIC_{50}$ | $IC_{50}$ Value [nM] |
|---|---|---|---|---|
| compound_119 | Z1411351789 | CC=1C=C(C=CC1S(=O)(=O)N)C(=O)N2CCOCC2CC(=O)C3=CC=CS3 | 7.09 | 82 |
| compound_120 | Z1466710623 | CN(C)C(=O)C1CN(CCN1C(=O)C=2C=CC=3NC=CC3C2)C(=O)C=4C=CC=5NC=CC5C4 | 4.52 | 30000 |
| compound_121 | Z1498230862 | CC1=CSC(NC=2C=CC(NC(=O)C=3C=C(C=C(C)C3C)S(=O)(=O)N)=CC2)=N1 | 6.76 | 174 |
| compound_122 | Z1525016622 | CC=1C=CC(=C(C1)C(=O)NC=2C=CC(NC(=O)NCC3CCCO3)=CC2)S(=O)(=O)N | 5.10 | 7882 |
| compound_123 | Z1576709178 | CC(OC(=O)C1(CC=2C=CC(Cl)=CC2)CC1)C(=O)NC=3C=CC=C(C3)S(=O)(=O)N | 7.42 | 38 |
| compound_124 | Z1894623041 | CC=1C=C(CNC(=O)[C@@H](NC(=O)OC(C)(C)C)C(C)(C)C)C=CC1S(=O)(=O)N | 5.86 | 1372 |
| compound_125 | Z194572336 | CC=1C=CC(NC=2C=CC=CC2C(=O)OCC(=O)NC=3C=CC(=CC3)S(=O)(=O)N)=CC1 | 7.11 | 77 |
| compound_126 | Z1982849991 | CN(C)CC=1C=CC(=NC1)N2CCN(CC2)C(=O)C=3C=CC(Cl)=C(C3)S(=O)(=O)N | 5.24 | 5704 |
| compound_127 | Z2014120909 | COC=1C=CC(=CC1C(=O)NC2CCN(C2)C(=O)NC=3C=CC=CC3)S(=O)(=O)N | 4.52 | 30000 |
| compound_128 | Z2185821639 | CC1CCCN(C1)C=2C=C(C=CN2)C(=O)N3CCN(CC3)C=4C=CC(=CC4)S(=O)(=O)N | 7.08 | 84 |

| Compound_id | Example_id | SMILES | $pIC_{50}$ | $IC_{50}$ Value [nM] |
|---|---|---|---|---|
| compound_129 | Z2230474937 | CC(C)(CCCNC(=O)C=1C=CC=2NC=CC2C1)NC(=O)C=3C=CC=4NC=CC4C3 | 4.52 | 30000 |
| compound_130 | Z228627984 | NS(=O)(=O)C=1C=CC(NC(=O)C=2C=3CCCC(=CC4=CC=CO4)C3N=C5C=CC=CC25)=CC1 | 6.71 | 197 |
| compound_131 | Z236079254 | CC1=CC(C(=O)NC=2C=CC(NC(=O)C3=CC=CO3)=CC2)=C(C)N1CC=4C=CC(=CC4)S(=O)(=O)N | 7.85 | 14 |
| compound_132 | Z2474885859 | NS(=O)(=O)C=1C=CC(NC(=S)N(CC2CC2)CC=3C=CC=C(O)C3)=CC1 | 7.68 | 21 |
| compound_133 | Z254592874 | CC=1C=C(C=C(C1C)S(=O)(=O)N)C(=O)NC=2C=CC(NC(=O)NC=3C=CC=CC3)=CC2 | 6.61 | 248 |
| compound_134 | Z263202688 | CC(OC(=O)C=1C=CC(Cl)=C(C1)S(=O)(=O)N)C(=O)NCCC2=CNC=3C=CC=CC23 | 6.53 | 293 |
| compound_135 | Z295990146 | NS(=O)(=O)C=1C=CC=C(NC(=O)C2CC=3C=CC=CC3CN2C(=O)C4=CC=CO4)C1 | 5.75 | 1785 |
| compound_136 | Z3289831644 | NS(=O)(=O)C=1C=CC(F)=C(CNC=2C(=CC=C3C=CC=CC23)S(=O)(=O)N)C1 | 7.30 | 50 |
| compound_137 | Z362005216 | NS(=O)(=O)C=1C=CC(NC(=O)N2CCN(CC3=CC=C(S3)C=4C=CC=C(Cl)C4)CC2)=CC1 | 7.28 | 53 |
| compound_138 | Z365446158 | CC(C)C(=O)NC=1C=CC(=CC1)C(=O)N2CCC(CC2)C(=O)NC=3C=CC(=CC3)S(=O)(=O)N | 7.43 | 37 |

| Compound_id | Example_id | SMILES | $pIC_{50}$ | $IC_{50}$ Value [nM] |
|---|---|---|---|---|
| compound_139 | Z367584278 | NS(=O)(=O)C=1C=CC=C(CNC(=O)C=2C=C(C3=CC=CO3)N(N2)C=4C=CC(Cl)=CC4)C1 | 4.52 | 30000 |
| compound_140 | Z423226006 | O=C(N1CCN(CC1)C(=O)C2=CC=C(S2)C(=O)N3CCN(CC3)C(=O)C4=CC=CO4)C5=CC=CO5 | 4.52 | 30000 |
| compound_141 | Z458308386 | CC(N(C)C(=O)C1CCCN(C1)C(=O)C=2C=C(C)C=C(C)C2)C=3C=CC(=CC3)S(=O)(=O)N | 7.22 | 60 |
| compound_142 | Z644056738 | CC(N(C)CC(=O)NC(C1=CC=CS1)C(C)(C)C)C=2C=CC(=CC2)S(=O)(=O)N | 7.12 | 75 |
| compound_143 | Z646387096 | COC(=O)C=1C=C(C=CC1NCC=2C=CC=C(NC(=O)NC=3C=CC=C(C)C3)C2)S(=O)(=O)N | 6.09 | 810 |
| compound_144 | Z794444524 | CC=1C=CC(NC(=O)COC(=O)C=2C=CC(O)=C(C2)C=3C=CC=CC3)=CC1S(=O)(=O)N | 6.55 | 279 |
| compound_145 | Z803847706 | CC(C)(C)OC(=O)N1CCC(CC1)(NC=2C=CC=CC2)C(=O)NC=3C=CC(=CC3)S(=O)(=O)N | 4.52 | 30000 |
| compound_146 | Z807338068 | NS(=O)(=O)C=1C=CC(CCNC(=O)NC=2C=CC(NC(=O)C=3C=CC(Cl)=CC3)=CC2)=CC1 | 7.02 | 96 |
| compound_147 | Z838613390 | CC(NC(=O)NCC=1C=CC(=CC1)C(=O)N2CC(C)OC(C)C2)C=3C=CC=C(C3)S(=O)(=O)N | 4.52 | 30000 |
| compound_148 | Z847412398 | NS(=O)(=O)C=1C=C(C=CC1Cl)C(=O)OCC2=CSC(CC(=O)NC=3C=CC=CC3)=N2 | 6.84 | 144 |

| Compound_id | Example_id | SMILES | $pIC_{50}$ | $IC_{50}$ Value [nM] |
|---|---|---|---|---|
| compound_149 | Z912254624 | CC=1C=CN(N1)C=2C=CC=C(C2)C(=O)OCC(=O)C=3C=C(C)N(C3C)C=4C=CC(=CC4)S(=O)(=O)N | 6.82 | 151 |
| compound_150 | Z954558418 | CC=1C=C(C=CC1NC(=O)COC(=O)C=2C=C(C=CC2O)S(=O)(=O)N)N3CCCC3 | 6.13 | 746 |
| compound_151 | Z977212584 | NS(=O)(=O)C=1C=CC(N2CCOCC2)=C(C1)C(=O)NC=3C=CC(NC(=O)C4=CC=CS4)=CC3 | 7.39 | 41 |
| compound_152 | Z994785762 | CC=1C=CC(NC(=O)C2CCCN2C(=O)C=3C=CC=4C=CC=CC4C3O)=CC1S(=O)(=O)N | 4.52 | 30000 |

**Table S4. Compound derivatives equipped with reaction handles (capped) for *in vitro* and *in vivo* validation.**

| Capped_compound_id | Capped_SMILES | Capped_pIC$_{50}$ |
|---|---|---|
| capped_compound_131c-i | CC1=CC(C(=O)NC=2C=CC=CC2C(=O)NCC3=CN(C)N=N3)=C(C)N1CC=4C=CC(=CC4)S(=O)(=O)N | 7.41 |
| capped_compound_131c-ii | CC1=CC(C(=O)NC=2C=CC(=CC2)C3=CN(C)N=N3)=C(C)N1CC=4C=CC(=CC4)S(=O)(=O)N | 7.74 |
| capped_compound_68c-i | CC1=NN(C(C)=C1CCC(=O)NC=2C=CC=CC2C(=O)NCC3=CN(C)N=N3)C=4C=CC(=CC4)S(=O)(=O)N | 7.41 |
| capped_compound_68c-ii | CC1=NN(C(C)=C1CCC(=O)NC=2C=CC(=CC2)C3=CN(C)N=N3)C=4C=CC(=CC4)S(=O)(=O)N | 6.33 |
| capped_compound_68c-iii | CC1=NN(C(C)=C1CCC(=O)NC=2C=CC=C(OCC3=CN(C)N=N3)C2)C=4C=CC(=CC4)S(=O)(=O)N | 7.93 |
| capped_compound_68c-iv | CC1=NN(C(C)=C1CCC(=O)NC=2C=CC(OCC3=CN(C)N=N3)=CC2)C=4C=CC(=CC4)S(=O)(=O)N | 7.62 |
| capped_compound_68c-v | CC1=NN(C(C)=C1CCC(=O)NC=2C=CC=C(C2)C3=CN(C)N=N3)C=4C=CC(=CC4)S(=O)(=O)N | 7.57 |
| capped_compound_12a | CC(=O)Nc1ccc(NC(=O)c2ccc(Cl)c(S(N)(=O)=O)c2)cc1 | 6.06 |
| capped_compound_109b | CNC(=O)C=1C=CC=C2N(CC=3C=COC3C(=O)NC=4C=CC(=CC4)S(=O)(=O)N)C(C)CC21 | 7.38 |
| capped_compound_109b-ii | CNC(=O)C=1C=CC=2N(CC=3C=COC3C(=O)NC=4C=CC(=CC4)S(=O)(=O)N)C(C)CC2C1 | 7.70 |
| capped_compound_109a-ii | CC1CC=2C=C(NC(=O)C)C=CC2N1CC=3C=COC3C(=O)NC=4C=CC(=CC4)S(=O)(=O)N | 7.32 |
| capped_compound_109a | CC1CC=2C(=CC=CC2NC(=O)C)N1CC=3C=COC3C(=O)NC=4C=CC(=CC4)S(=O)(=O)N | 7.43 |
| capped_compound_109c | CC1CC=2C(=CC=CC2C3=CN(C)N=N3)N1CC=4C=COC4C(=O)NC=5C=CC(=CC5)S(=O)(=O)N | 7.95 |
| capped_compound_109c-ii | CC1CC=2C=C(C=CC2N1CC=3C=COC3C(=O)NC=4C=CC(=CC4)S(=O)(=O)N)C5=CN(C)N=N5 | 6.88 |

| Capped_compound_id | Capped_SMILES | Capped_pIC$_{50}$ |
|---|---|---|
| capped_compound_104c | CC(NCC1(CCOCC1)C=2C=CC(=CC2)C3=CN(C)N=N3)C(=O)NC=4C=CC(=CC4)S(=O)(=O)N | 7.00 |
| capped_compound_104a | CC(NCC1(CCOCC1)C=2C=CC(NC(=O)C)=CC2)C(=O)NC=3C=CC(=CC3)S(=O)(=O)N | 7.07 |
| capped_compound_104a-ii | CC(NCC1(CCOCC1)C=2C=CC=C(NC(=O)C)C2)C(=O)NC=3C=CC(=CC3)S(=O)(=O)N | 6.94 |
| capped_compound_104c-ii | CC(NCC1(CCOCC1)C=2C=CC=C(C2)C3=CN(C)N=N3)C(=O)NC=4C=CC(=CC4)S(=O)(=O)N | 6.93 |
| capped_compound_82b | CNC(=O)C=1C=CC(N2CCN(CC2)C(=O)C=3C=CC(=C(Br)C3)S(=O)(=O)N)=C4C=CNC14 | 7.41 |
| capped_compound_82c | CN1C=C(N=N1)C=2C=CC(N3CCN(CC3)C(=O)C=4C=CC(=C(Br)C4)S(=O)(=O)N)=C5C=CNC25 | 7.89 |
| capped_compound_76a | CC(N(C)C(=O)C=1C=C(C=CC1O)S(=O)(=O)N)C=2C=CC=C(C2)C=3C=CC(NC(=O)C)=CC3 | 7.05 |
| capped_compound_76c | CC(N(C)C(=O)C=1C=C(C=CC1O)S(=O)(=O)N)C=2C=CC=C(C2)C=3C=CC(=CC3)C4=CN(C)N=N4 | 7.66 |
| capped_compound_76b | CNC(=O)C=1C=CC(=CC1)C=2C=CC=C(C2)C(C)N(C)C(=O)C=3C=C(C=CC3O)S(=O)(=O)N | 7.24 |
| capped_compound_61a-ii | CC(=O)NC=1C=CC(NC(=O)CN(CC=2C=CC(Cl)=CC2)C(=O)C=3C=CC(=C(C)C3)S(=O)(=O)N)=C(F)C1 | 7.82 |
| capped_compound_61c-ii | CC=1C=C(C=CC1S(=O)(=O)N)C(=O)N(CC(=O)NC=2C=CC(=CC2F)C3=CN(C)N=N3)CC=4C=CC(Cl)=CC4 | 7.92 |
| capped_compound_61b-ii | CNC(=O)C=1C=CC(NC(=O)CN(CC=2C=CC(Cl)=CC2)C(=O)C=3C=CC(=C(C)C3)S(=O)(=O)N)=C(F)C1 | 7.54 |
| capped_compound_61a | CC(=O)NC=1C=CC(CN(CC(=O)NC=2C=CC=CC2F)C(=O)C=3C=CC(=C(C)C3)S(=O)(=O)N)=CC1 | 7.38 |

| Capped_compound_id | Capped_SMILES | Capped_pIC$_{50}$ |
|---|---|---|
| capped_compound_61c | CC=1C=C(C=CC1S(=O)(=O)N)C(=O)N(CC(=O)NC=2C=CC=CC2F)CC=3C=CC(=CC3)C4=CN(C)N=N4 | 7.57 |
| capped_compound_53b | CNC(=O)C=1C=CC=2OC(=CC2C1)C(=O)NC3CCCN(CC(=O)NC=4C=CC(=CC4)S(=O)(=O)N)C3 | 7.02 |
| capped_compound_53c-ii | CN1C=C(N=N1)C=2C=CC=C3OC(=CC23)C(=O)NC4CCCN(CC(=O)NC=5C=CC(=CC5)S(=O)(=O)N)C4 | 7.52 |
| capped_compound_53b-ii | CNC(=O)C=1C=CC=C2OC(=CC12)C(=O)NC3CCCN(CC(=O)NC=4C=CC(=CC4)S(=O)(=O)N)C3 | 7.15 |
| capped_compound_53a-ii | CC(=O)NC=1C=CC=C2OC(=CC12)C(=O)NC3CCCN(CC(=O)NC=4C=CC(=CC4)S(=O)(=O)N)C3 | 7.37 |
| capped_compound_53c | CN1C=C(N=N1)C=2C=CC=3OC(=CC3C2)C(=O)NC4CCCN(CC(=O)NC=5C=CC(=CC5)S(=O)(=O)N)C4 | 7.48 |
| capped_compound_53a | CC(=O)NC=1C=CC=2OC(=CC2C1)C(=O)NC3CCCN(CC(=O)NC=4C=CC(=CC4)S(=O)(=O)N)C3 | 7.17 |
| capped_compound_43a | CC(=O)NC=1C=CC(=CC1)N2N=C(C=C2C3=CC=CO3)C(=O)NC=4C=CC(=CC4)S(=O)(=O)N | 7.22 |
| capped_compound_43b | CNC(=O)C=1C=CC(=CC1)N2N=C(C=C2C3=CC=CO3)C(=O)NC=4C=CC(=CC4)S(=O)(=O)N | 8.00 |
| capped_compound_43c | CN1C=C(N=N1)C=2C=CC(=CC2)N3N=C(C=C3C4=CC=CO4)C(=O)NC=5C=CC(=CC5)S(=O)(=O)N | 8.85 |
| capped_compound_29c | CC1=CC(C(=O)CN2CCN(CC2)C(=O)C=3C=CC(C4=CN(C)N=N4)=C(F)C3)=C(C)N1C=5C=CC(=CC5)S(=O)(=O)N | 7.46 |
| capped_compound_29a | CC(=O)NC=1C=CC(=CC1F)C(=O)N2CCN(CC(=O)C=3C=C(C)N(C3C)C=4C=CC(=CC4)S(=O)(=O)N)CC2 | 7.29 |

| Capped_compound_id | Capped_SMILES | Capped_pIC$_{50}$ |
| --- | --- | --- |
| capped_compound_29b | CNC(=O)C=1C=CC(=CC1F)C(=O)N2CCN(CC(=O)C=3C=C(C)N(C3C)C=4C=CC(=CC4)S(=O)(=O)N)CC2 | 7.17 |
| capped_compound_26a | CC(=O)NC=1C=CC(CN2C(=CC=3SC=CC23)C(=O)NC=4C=CC(=CC4)S(=O)(=O)N)=CC1 | 8.13 |
| capped_compound_26c | CN1C=C(N=N1)C=2C=CC(CN3C(=CC=4SC=CC34)C(=O)NC=5C=CC(=CC5)S(=O)(=O)N)=CC2 | 8.94 |
| capped_compound_19a | CC(=O)NC=1C=CC(NC(=O)NC2=NC(CC(=O)NC=3C=CC(=CC3)S(=O)(=O)N)=CS2)=CC1 | 7.52 |
| capped_compound_19c | CN1C=C(N=N1)C=2C=CC(NC(=O)NC3=NC(CC(=O)NC=4C=CC(=CC4)S(=O)(=O)N)=CS3)=CC2 | 7.53 |
| capped_compound_19b | CNC(=O)C=1C=CC(NC(=O)NC2=NC(CC(=O)NC=3C=CC(=CC3)S(=O)(=O)N)=CS2)=CC1 | 7.53 |
| capped_compound_17c | CN1C=C(N=N1)C=2C=CC(=NN2)N3CCCC(C3)C(=O)NC=4C=CC(=CC4)S(=O)(=O)N | 7.56 |

**Table S5. Structures and characterization of lead compounds by CAIX inhibition assay and fluorescence polarization measurements.**

| Compound_id | SMILES | pIC$_{50}$ | K$_D$ (n=5) [M] |
|---|---|---|---|
| compound_68 | CC1=NN(C(C)=C1CCC(=O)NC=2C=CC(=CC2)C3=CC=CN3)C=4C=CC(=CC4)S(=O)(=O)N | 7.82E-09 | |
| compound_109 | CC1CC=2C=CC=CC2N1CC=3C=COC3C(=O)NC=4C=CC(=CC4)S(=O)(=O)N | 7.92E-09 | |
| compound_61 | CC=1C=C(C=CC1S(=O)(=O)N)C(=O)N(CC(=O)NC=2C=CC=CC2F)CC=3C=CC(Cl)=CC3 | 7.89E-09 | |
| compound_53 | NS(=O)(=O)C=1C=CC(NC(=O)CN2CCCC(C2)NC(=O)C3=CC=4C=CC=CC4O3)=CC1 | 7.59E-09 | |
| compound_43 | NS(C1=CC=C(NC(C2=NN(C3=CC=C(Cl)C=C3)C(C3OC=CC=3)=C2)=O)C=C1)(=O)=O | 8.52E-09 | |
| compound_26 | N1(CC2=CC=C(C=C2)Cl)C2C=CSC=2C=C1C(=O)NC1=CC=C(C=C1)S(N)(=O)=O | 7.74E-09 | |
| capped_compound_68c-iii | CC1=NN(C(C)=C1CCC(=O)NC=2C=CC=C(OCC3=CN(C)N=N3)C2)C=4C=CC(=CC4)S(=O)(=O)N | 7.93E-09 | |
| capped_compound_68c-v | CC1=NN(C(C)=C1CCC(=O)NC=2C=CC=C(C2)C3=CN(C)N=N3)C=4C=CC(=CC4)S(=O)(=O)N | 7.57E-09 | |
| capped_compound_109b-ii | CNC(=O)C=1C=CC=2N(CC=3C=COC3C(=O)NC=4C=CC(=CC4)S(=O)(=O)N)C(C)CC2C1 | 7.70E-09 | |
| capped_compound_61c-ii | CC=1C=C(C=CC1S(=O)(=O)N)C(=O)N(CC(=O)NC=2C=CC(=CC2F)C3=CN(C)N=N3)CC=4C=CC(Cl)=CC4 | 7.92E-09 | |
| capped_compound_61a | CC(=O)NC=1C=CC(CN(CC(=O)NC=2C=CC=CC2F)C(=O)C=3C=CC(=C(C)C3)S(=O)(=O)N)=CC1 | 7.38E-09 | |

| Compound_id | SMILES | pIC$_{50}$ | K$_D$ (n=5) [M] |
|---|---|---|---|
| capped_compound_53a | CC(=O)NC=1C=CC=2OC(=CC2C1)C(=O)NC3CCCN(CC(=O)NC=4C=CC(=CC4)S(=O)(=O)N)C3 | 7.17E-09 | |
| capped_compound_43b | CNC(=O)C=1C=CC(=CC1)N2N=C(C=C2C3=CC=CO3)C(=O)NC=4C=CC(=CC4)S(=O)(=O)N | 8.00E-09 | |
| capped_compound_26c | CN1C=C(N=N1)C=2C=CC(CN3C(=CC=4SC=CC34)C(=O)NC=5C=CC(=CC5)S(=O)(=O)N)=CC2 | 8.94E-09 | |
| compound_68c-iii-FITC | C#CCOc1cccc(NC(=O)CCc2c(C)nn(-c3ccc(S(N)(=O)=O)cc3)c2C)c1 | | 5.68E-09 |
| compound_68c-v-FITC | C#Cc1cccc(NC(=O)CCc2c(C)nn(-c3ccc(S(N)(=O)=O)cc3)c2C)c1 | | 5.66E-09 |
| compound_109b-ii-FITC | CC1CC=2C=C(C=CC2N1CC=3C=COC3C(=O)NC=4C=CC(=CC4)S(=O)(=O)N)C(=O)O | | 4.41E-08 |
| compound_61c-ii-FITC | CC=1C=C(C=CC1S(=O)(=O)N)C(=O)N(CC(=O)NC=2C=CC(C#C)=CC2F)CC=3C=CC(Cl)=CC3 | | 1.13E-08 |
| compound_61a-FITC | CC=1C=C(C=CC1S(=O)(=O)N)C(=O)N(CC(=O)NC=2C=CC=CC2F)CC=3C=CC(N)=CC3 | | 1.25E-08 |
| compound_53a-FITC | NC=1C=CC=2OC(=CC2C1)C(=O)NC3CCCN(CC(=O)NC=4C=CC(=CC4)S(=O)(=O)N)C3 | | 9.63E-08 |
| compound_43b-FITC | NS(=O)(=O)C=1C=CC(NC(=O)C=2C=C(C3=CC=CO3)N(N2)C=4C=CC(=CC4)C(=O)O)=CC1 | | 6.48E-09 |
| compound_26c-FITC | NS(=O)(=O)C=1C=CC(NC(=O)C2=CC=3SC=CC3N2CC=4C=CC(C#C)=CC4)=CC1 | | 9.53E-09 |

### Supporting Information Figures

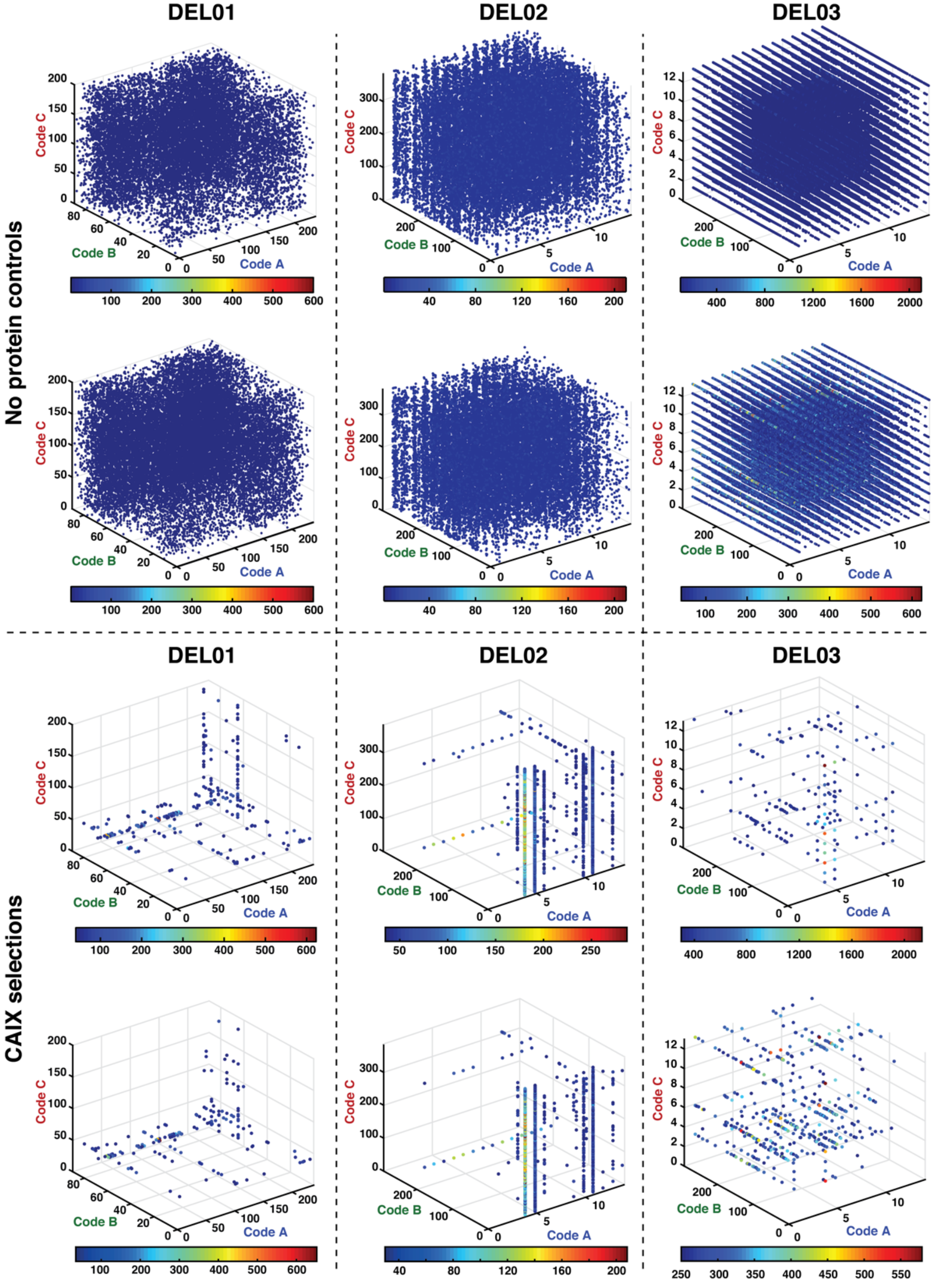


**Figure S1. Selection fingerprints of DEL01, DEL02, DEL03 screened against beads without immobilized protein (upper panel) and against Carbonic Anhydrase IX (lower panel) in duplicates. The *x*, *y* and *z*-coordinates correspond to code A, B, and C, respectively. DNA sequence counts are indicated by the colored jet scale. Cut-off = 4, 4, 40 and 40 (DEL01); 4, 4, 30 and 30 (DEL02), 100, 50, 300 and 250 (DEL03). Total counts = 1,551,875, 1,621714,**

**2,837,745 and 2,071,139 (DEL01); 1,817,134, 1,671,433, 1,964,585 and 2,159,325 (DEL02); 1,566,016, 2,010,344, 2,533,971 and 2,669,343 (DEL03).**

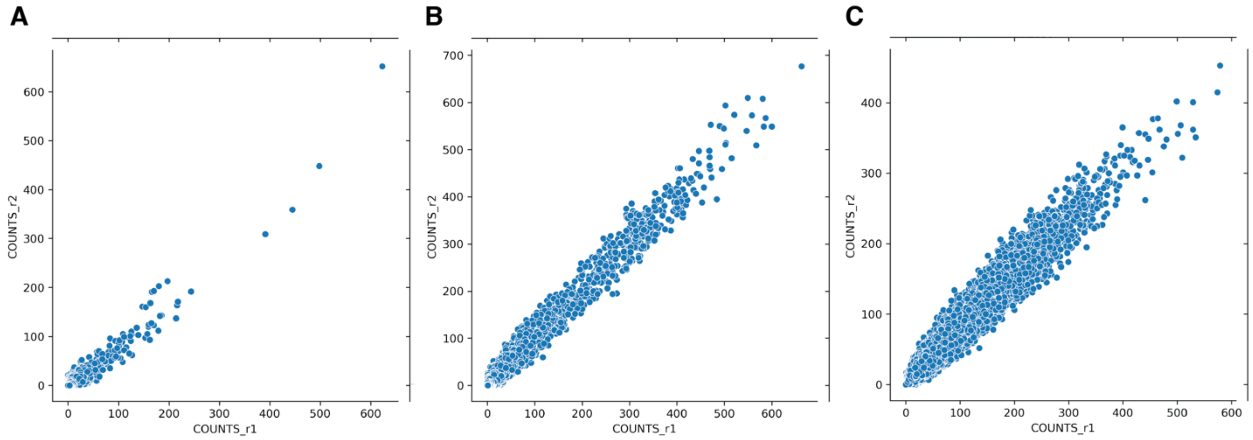


**Figure S2. Scatter plots between sequencing counts from two experimental replicates for each library.** (A) DEL01 (B) DEL02 (C) DEL03.

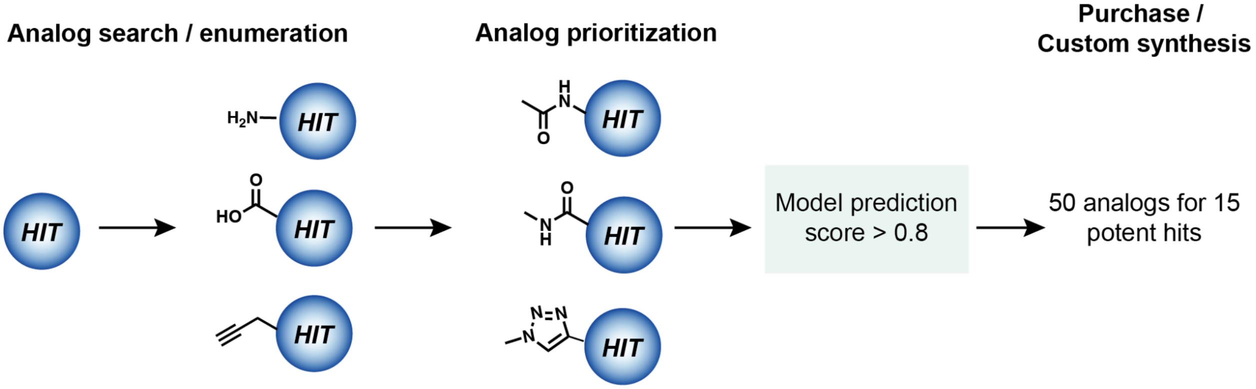


**Figure S3. Analog search and model prioritization.** For each HIT, analogs (ECFP6 Tanimoto similarity > 0.5) with primary amines, carboxylic acids, and alkynes are identified from Enamine REAL or computationally enumerated. Since the reaction handles are expected to undergo chemical changes after conjugation, surrogate compounds where the reaction handles are capped are created and re-scored by our trained models. Only compounds whose surrogate form scored above 0.8 were selected for experimental validation. A total of 50 analogs were selected for 15 potent hits.

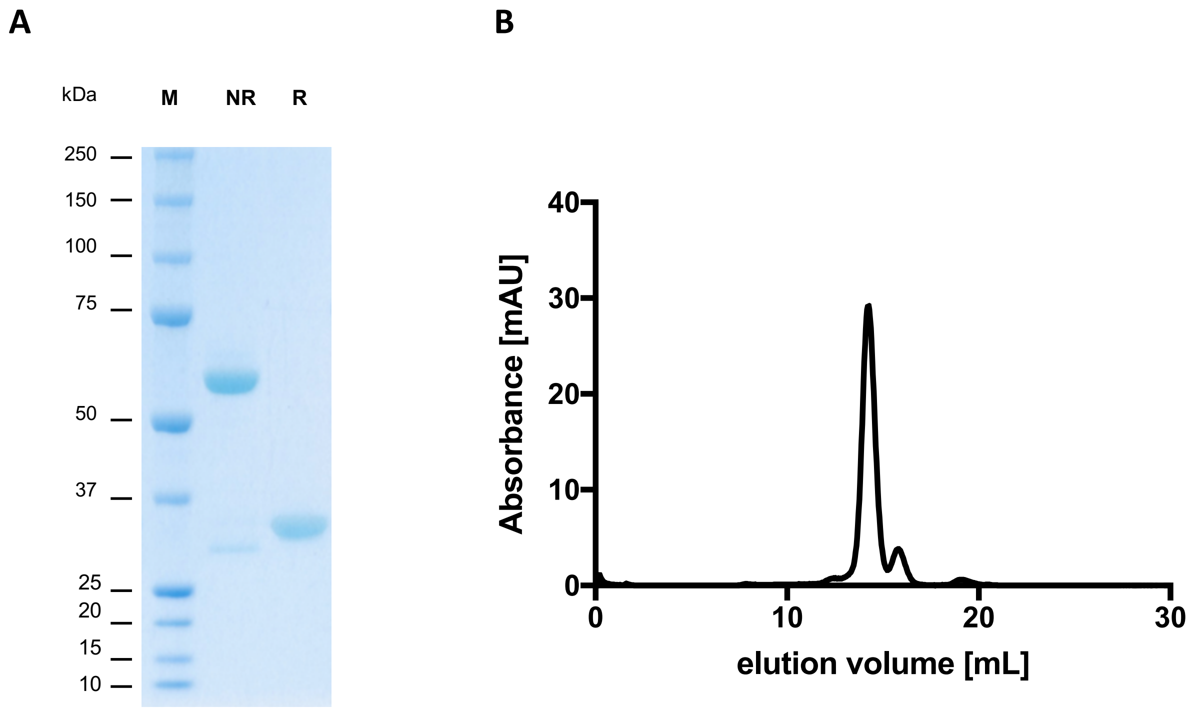


**Figure S4. Characterization of recombinantly produced human Carbonic Anhydrase IX.** Recombinant CAIX was characterized by SDS-page (A) and size-exclusion chromatography (B). SDS-PAGE was performed with 10% gels (Invitrogen, NP0302BOX) under reducing (R) and non-reducing (NR) conditions. For size-exclusion chromatography a Superdex 200 increase 10/300 GL column was used in combination with an ÄKTA PURE chromatography system (Cytiva).

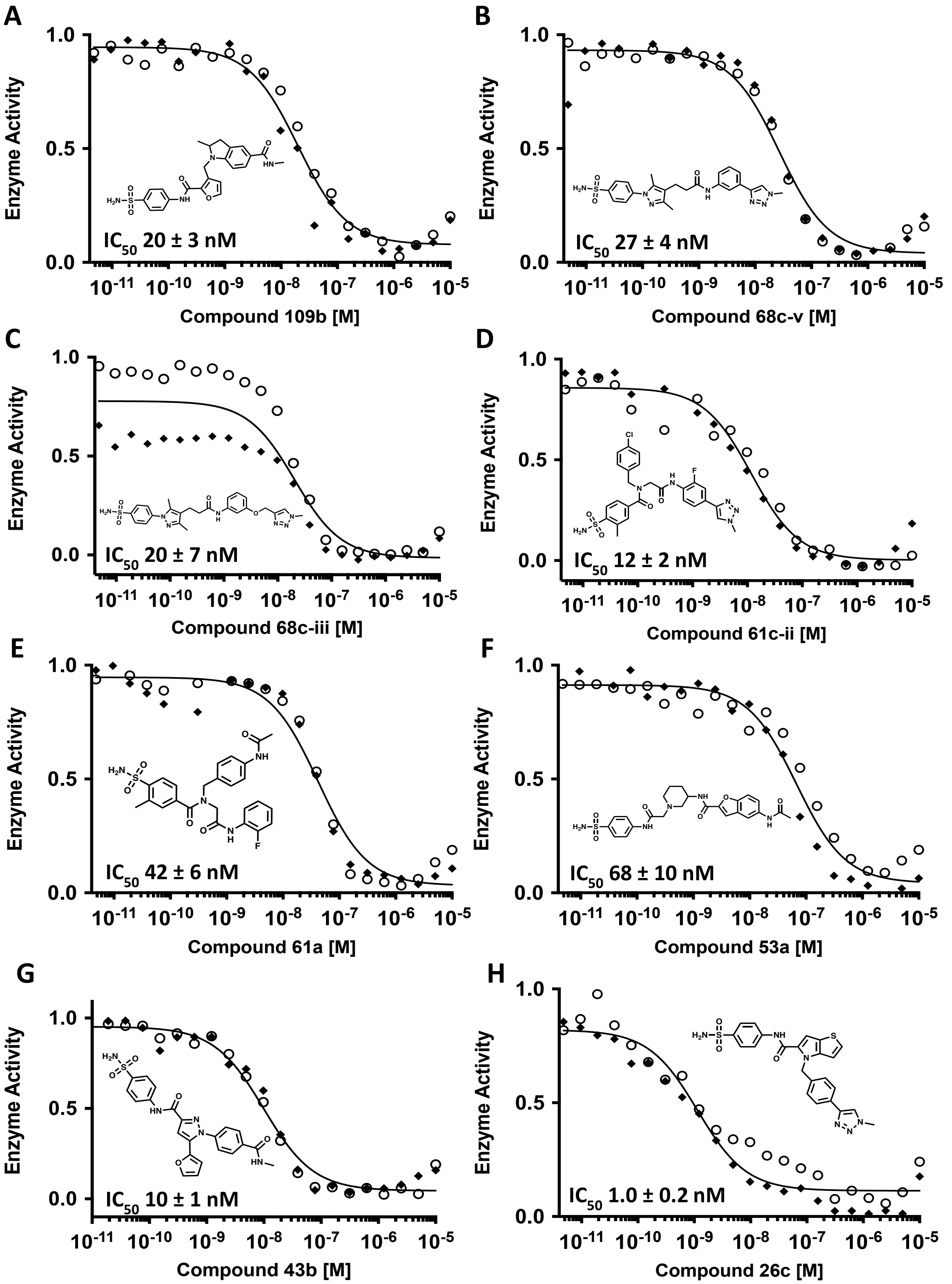


**Figure S5. CAIX inhibition assay of the 8 most potent capped candidates (see also Table S4).** The measurements have been performed in duplicates (n = 2).

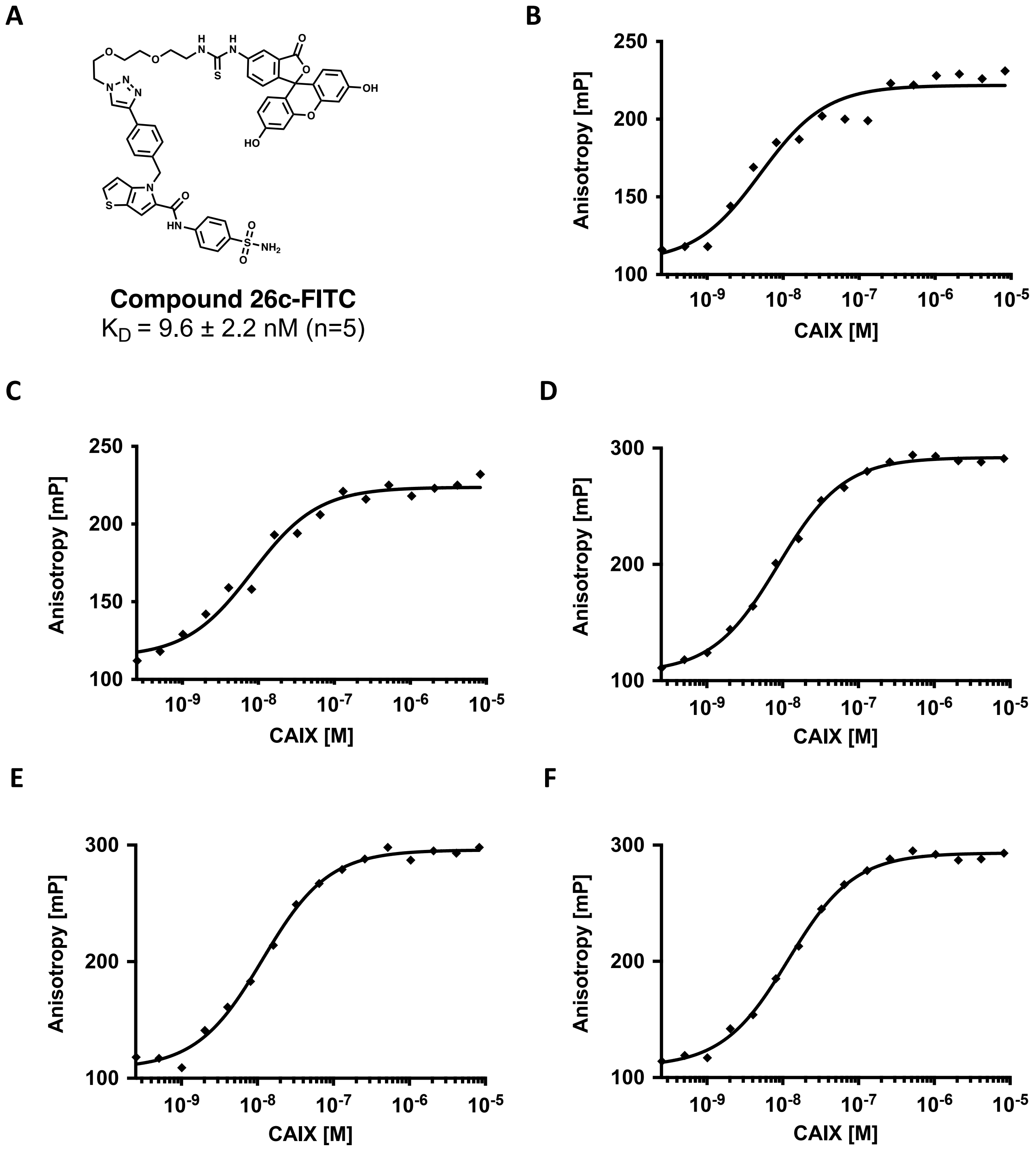


**Figure S6. Fluorescence polarization (FP) measurements of FITC-conjugated hits against carbonic anhydrase (CAIX).** Chemical structures (A) and Fluorescence Polarization (FP) measurements (B to F) of FITC-conjugate HIT 26c against carbonic anhydrase performed in replicates (n = 5).

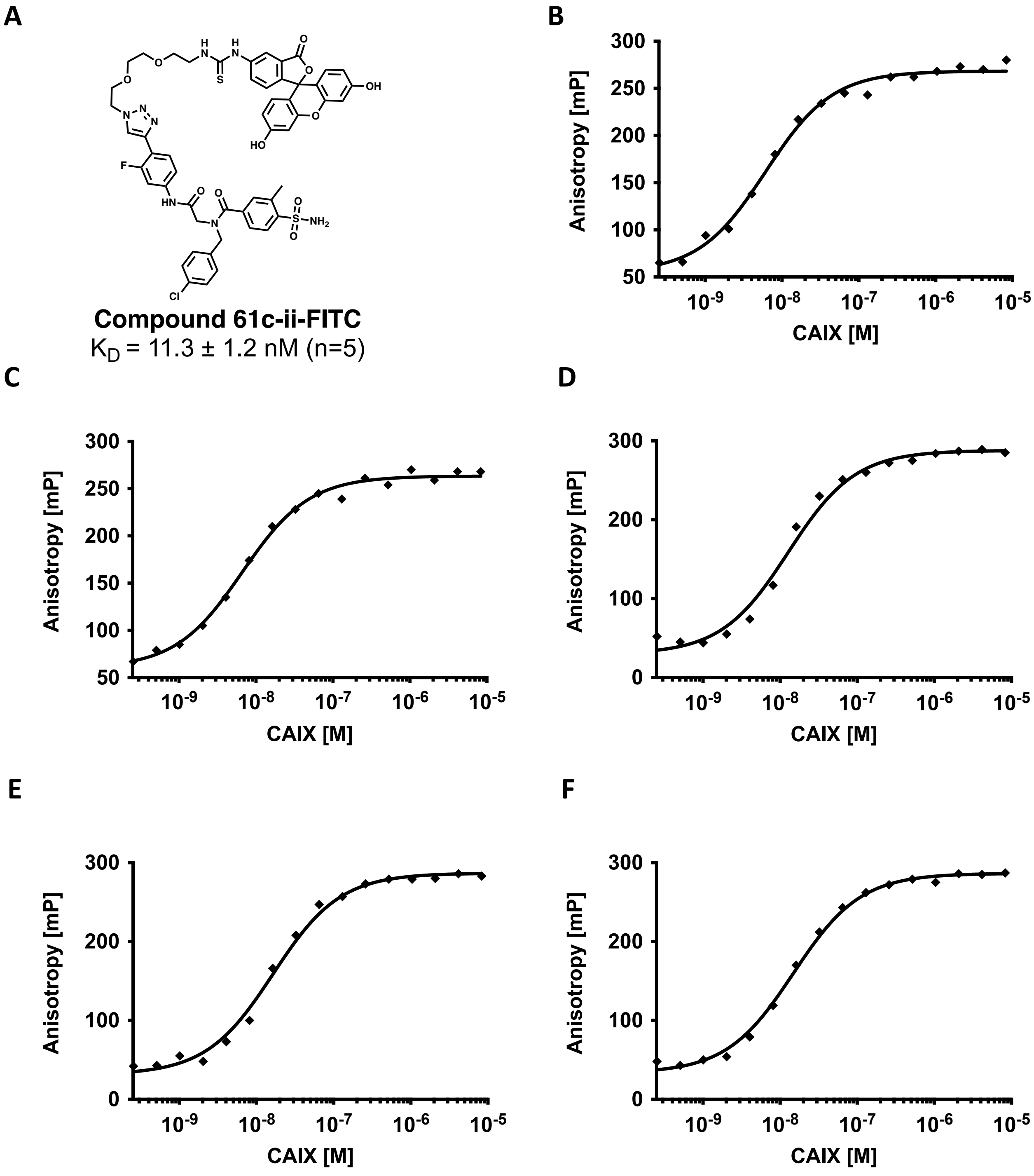


**Figure S7. Fluorescence polarization (FP) measurements of FITC-conjugated hits against carbonic anhydrase (CAIX).** Chemical structures (A) and Fluorescence Polarization (FP) measurements (B to F) of FITC-conjugate HIT 61c-ii against carbonic anhydrase performed in replicates (n = 5).

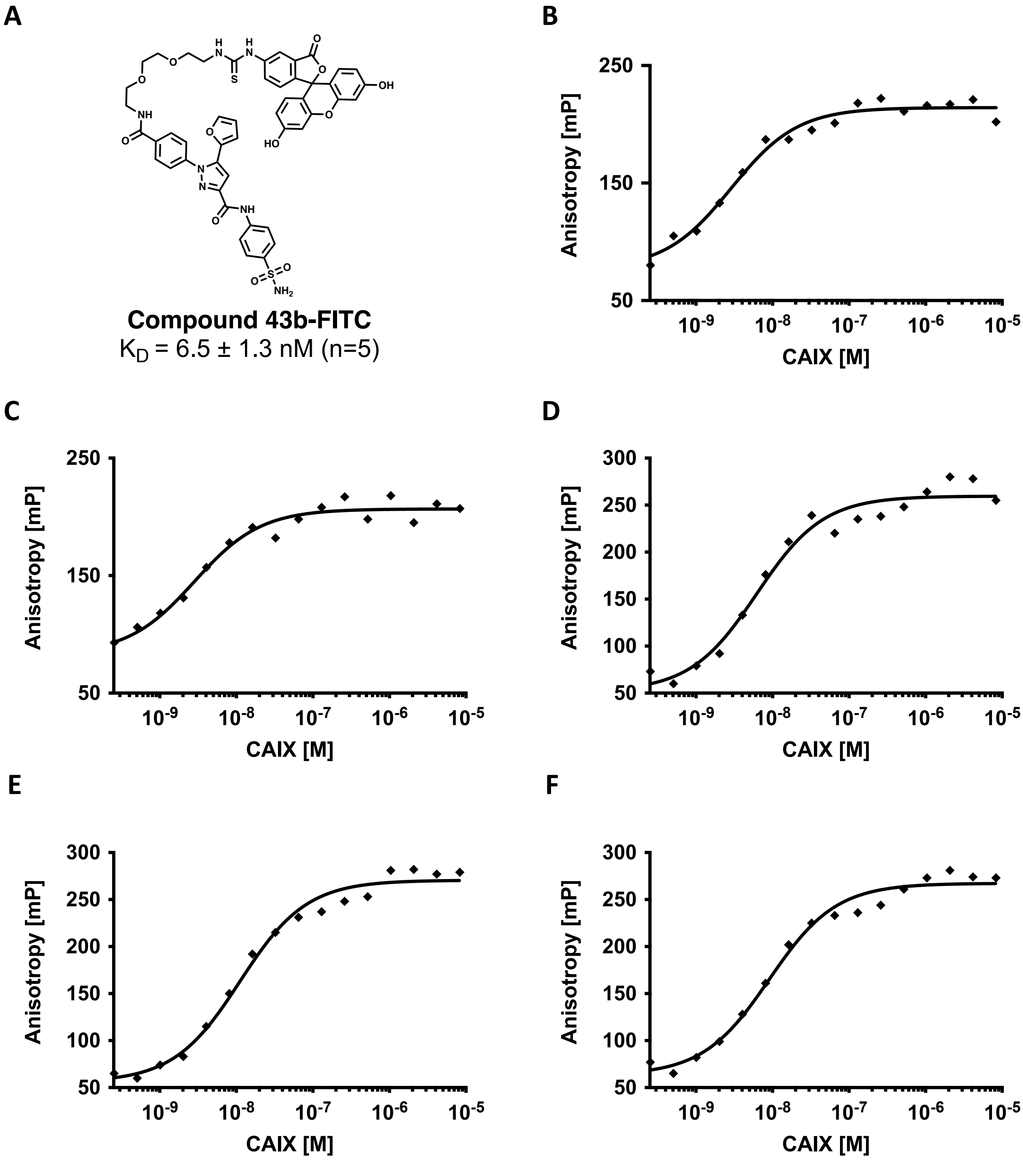


**Figure S8. Fluorescence polarization (FP) measurements of FITC-conjugated hits against carbonic anhydrase (CAIX).** Chemical structures (A.) and Fluorescence Polarization (FP) measurements (B. to F.) of FITC-conjugate HIT 43b against carbonic anhydrase performed in replicates (n = 5).

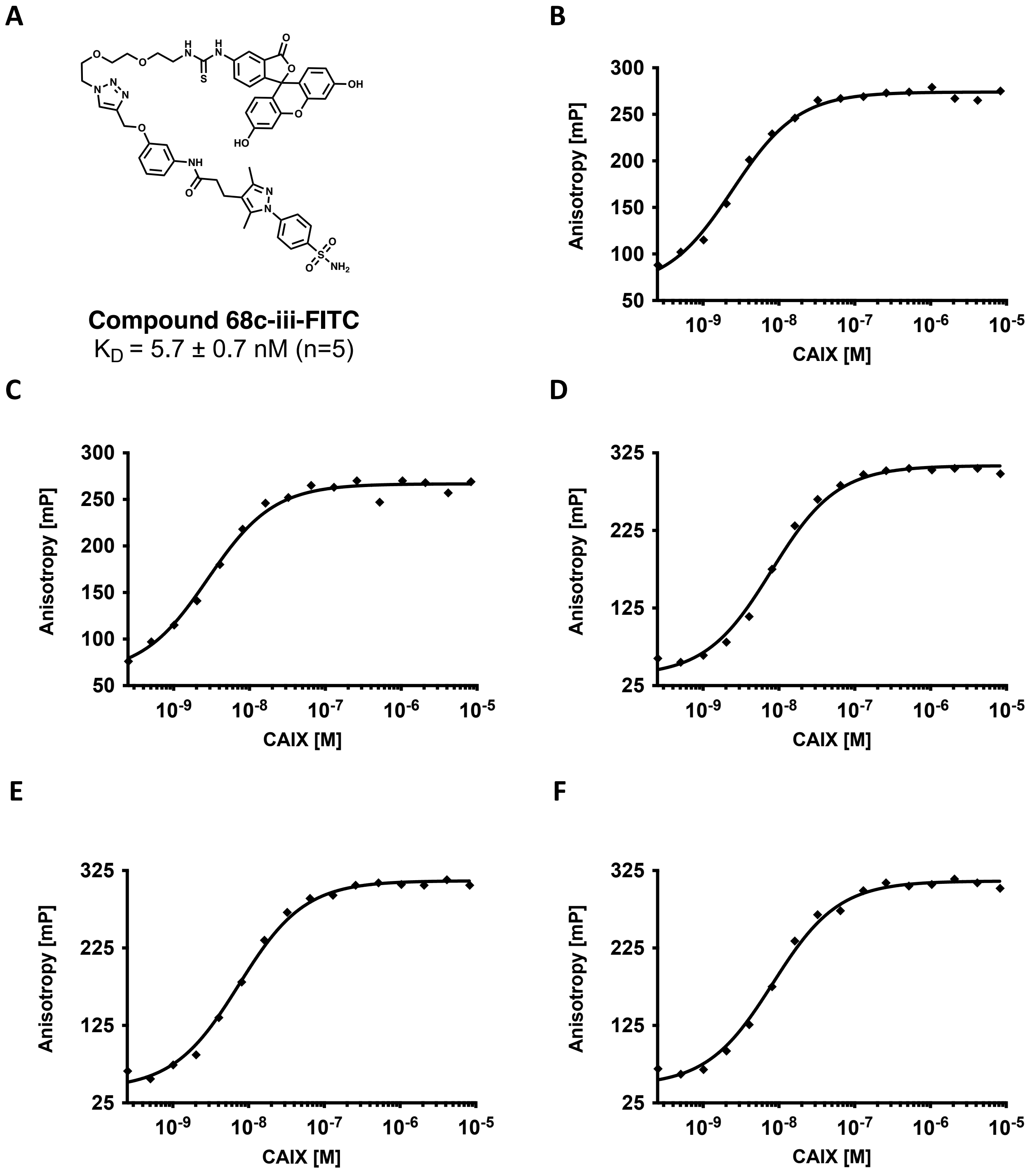


**Figure S9. Fluorescence polarization (FP) measurements of FITC-conjugated hits against carbonic anhydrase (CAIX).** Chemical structures (A.) and Fluorescence Polarization (FP) measurements (B. to F.) of FITC-conjugate HIT 68c-iii against carbonic anhydrase performed in replicates (n = 5).

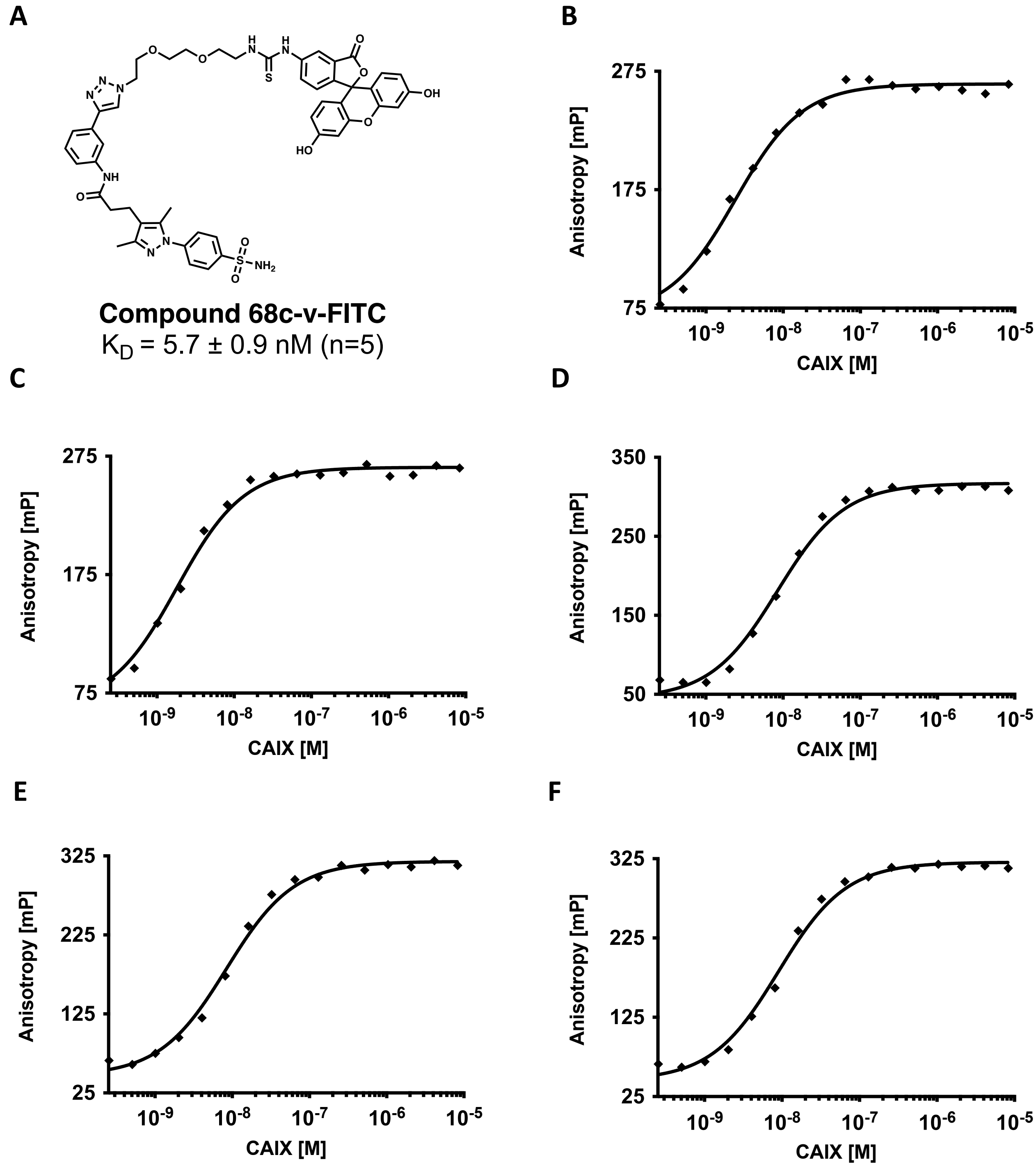


**Figure S10. Fluorescence polarization (FP) measurements of FITC-conjugated hits against carbonic anhydrase (CAIX).** Chemical structures (A.) and Fluorescence Polarization (FP) measurements (B. to F.) of FITC-conjugate HIT 68c-v against carbonic anhydrase performed in replicates (n = 5).

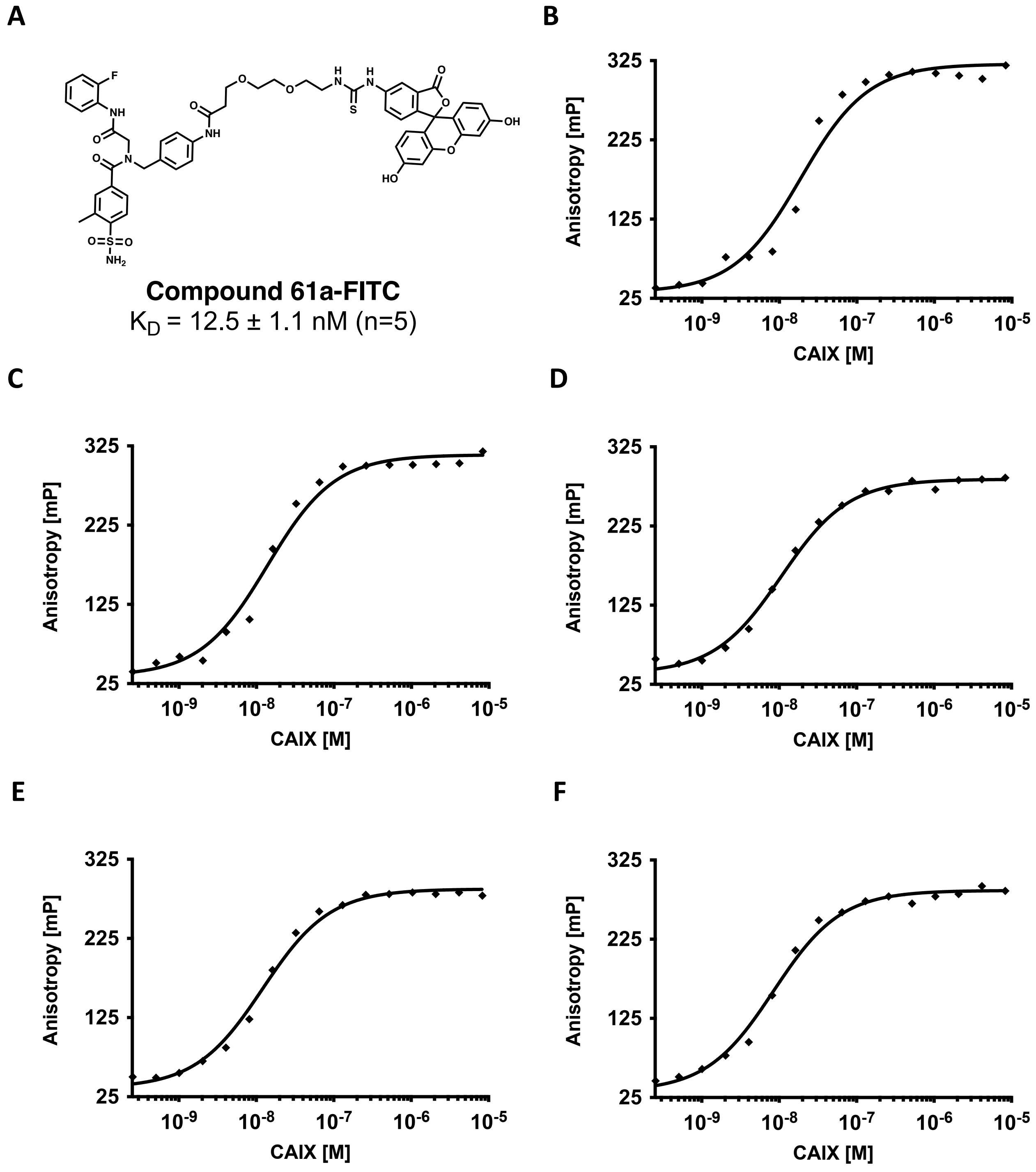


**Figure S11. Fluorescence polarization (FP) measurements of FITC-conjugated hits against carbonic anhydrase (CAIX).** Chemical structures (A.) and Fluorescence Polarization (FP) measurements (B. to F.) of FITC-conjugate HIT 61a against carbonic anhydrase performed in replicates (n = 5).

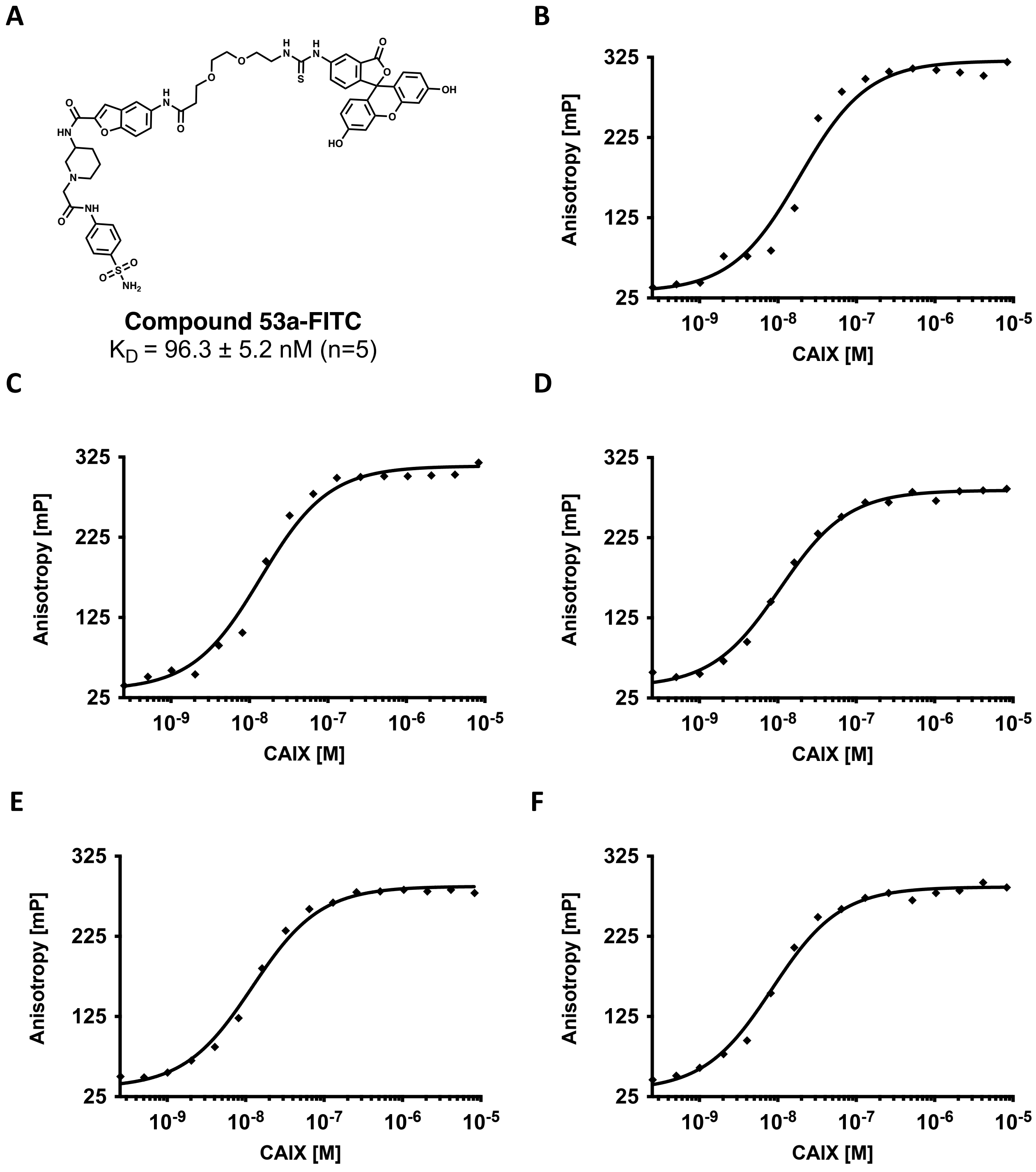


**Figure S12. Fluorescence polarization (FP) measurements of FITC-conjugated hits against carbonic anhydrase (CAIX).** Chemical structures (A.) and Fluorescence Polarization (FP) measurements (B. to F.) of FITC-conjugate HIT 53a against carbonic anhydrase performed in replicates (n = 5).

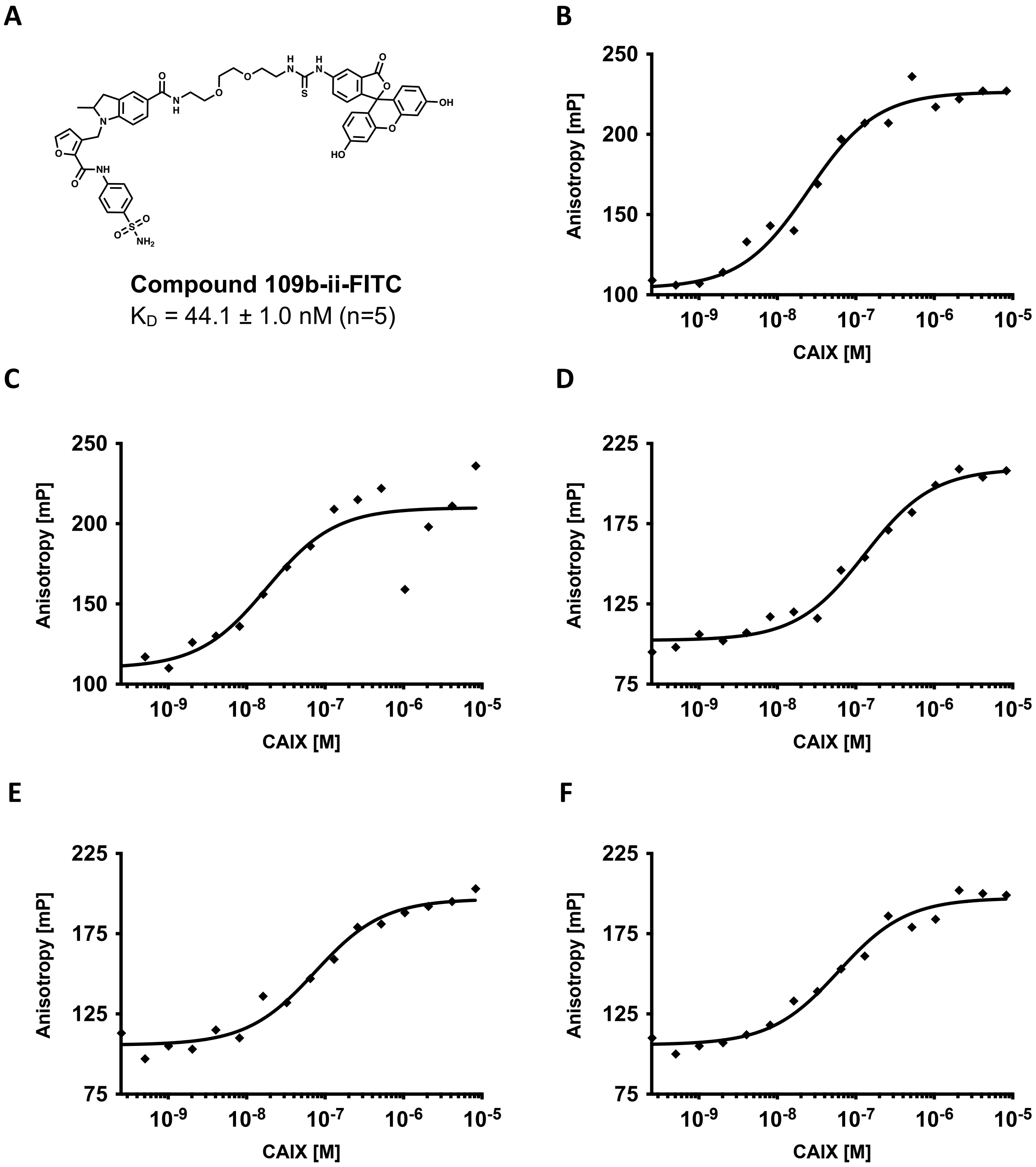


**Figure S13. Fluorescence polarization (FP) measurements of FITC-conjugated hits against carbonic anhydrase (CAIX).** Chemical structures (A.) and Fluorescence Polarization (FP) measurements (B. to F.) of FITC-conjugate HIT 109b-ii against carbonic anhydrase performed in replicates (n = 5).

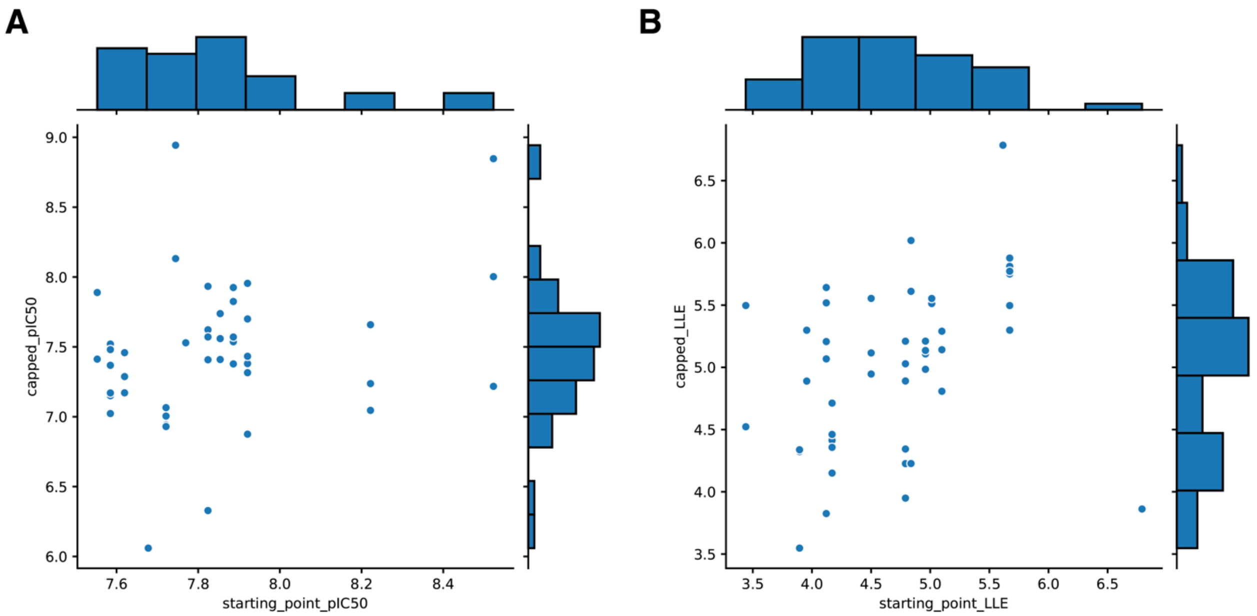


**Figure S14. pIC$_{50}$ and LLE of starting point and capped analogs (surrogate compounds).** 46 surrogate compounds were received and experimentally validated in enzymatic assays. (A) pIC50 of starting points (mean pIC50=7.841) and their capped analogs (mean pIC50=7.458). (B) LLE (pIC50 - clogP) of the starting points (mean LLE=4.7) and the surrogate compounds (mean LLE=5.0).

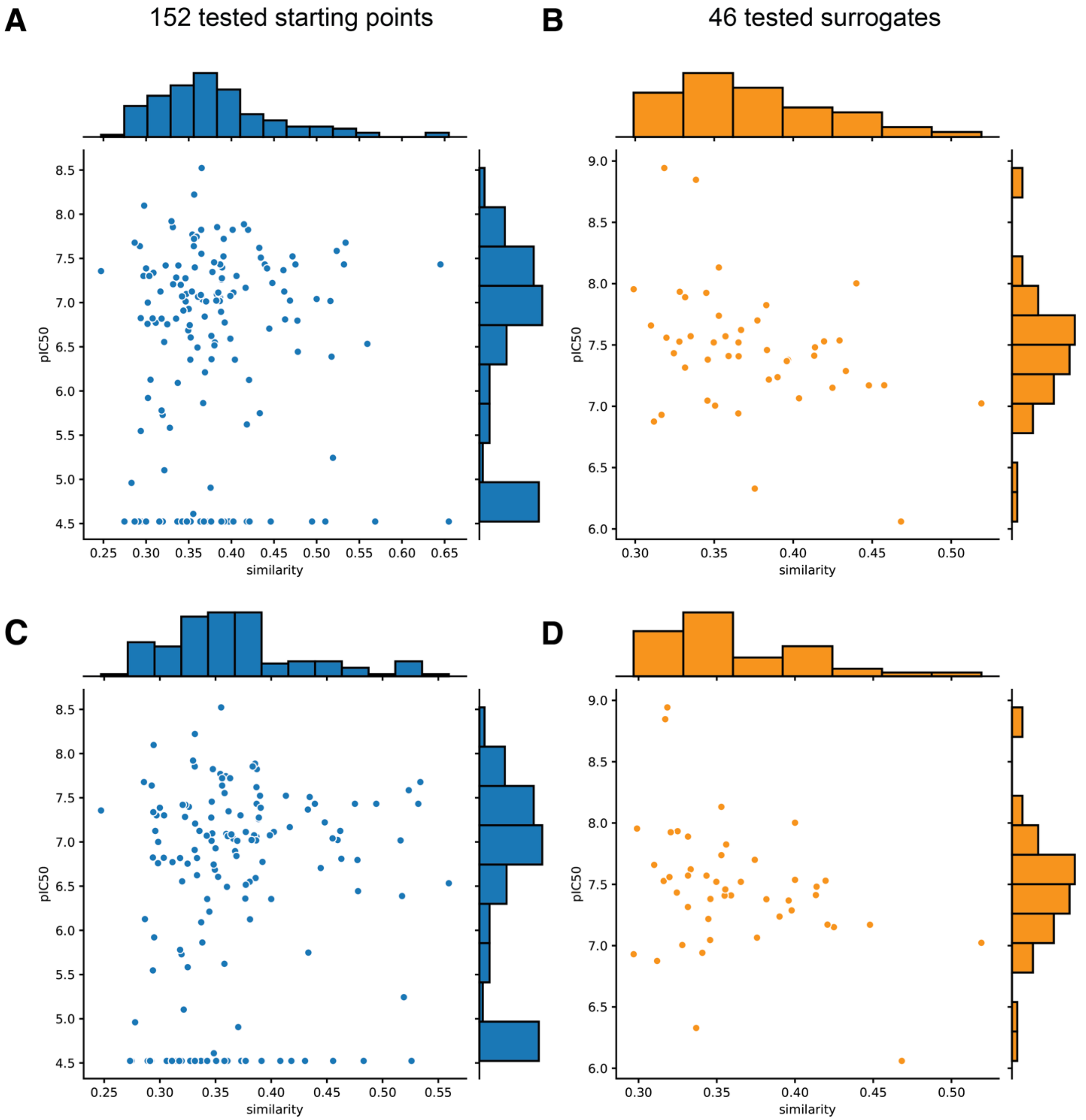


**Figure S15. Correlation between $pIC_{50}$ of tested compounds and ECFP6 (Morgan Fingerprint, radius=3) Tanimoto similarity to DEL training data.** For each subplot, the x-axis represents the nearest neighbor similarity of the tested compound to DEL training set and the y-axis represents the biochemical pIC50. (A) 152 tested compounds to all training DEL data (Spearman correlation = 0.0994) (B) 46 tested surrogate compounds to all training DEL data (Spearman correlation = -0.335) (C) 152 tested compounds to positive training examples (PTE) only (Spearman correlation = 0.1616) (D) 46 tested surrogate compounds to PTE only (Spearman correlation = -0.287).

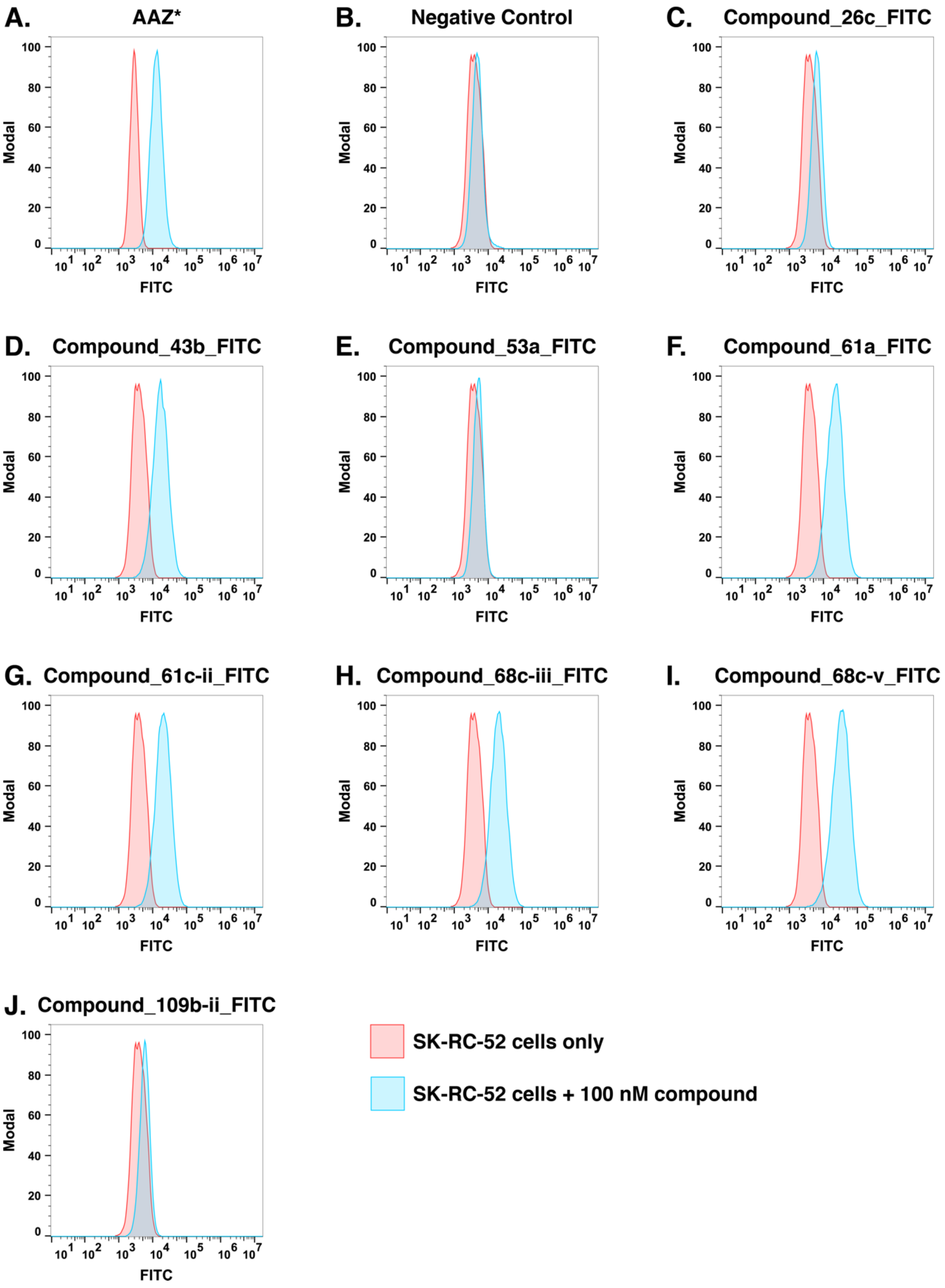


**Figure S16. Flow cytometry analysis of CAIX-positive SK-RC-52 cells in absence (red) and presence (blue) of FITC-conjugated compounds.**

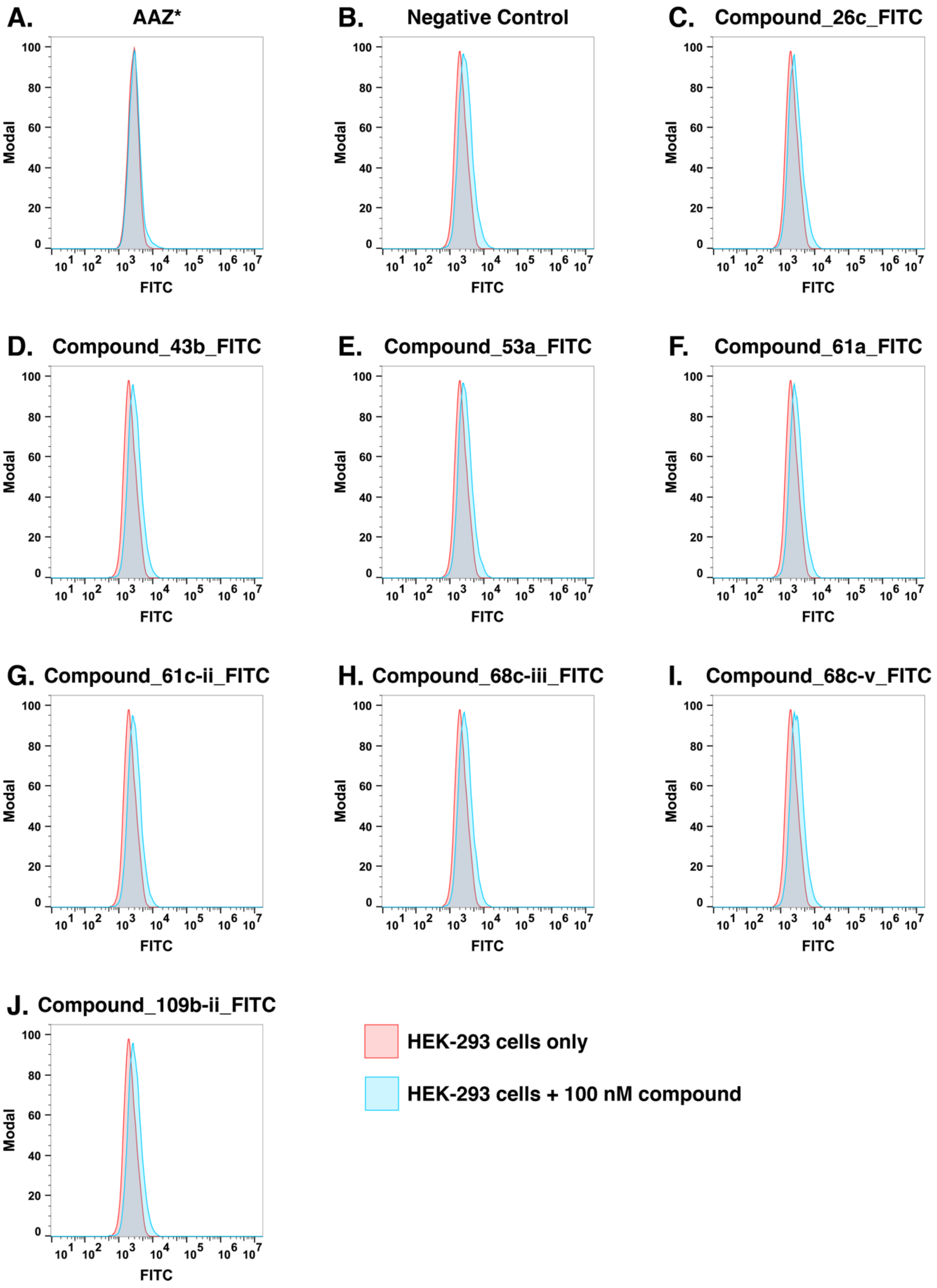


**Figure S17. Flow cytometry analysis of CAIX-negative HEK-293 cells in absence (red) and presence (blue) of FITC-conjugated compounds.**

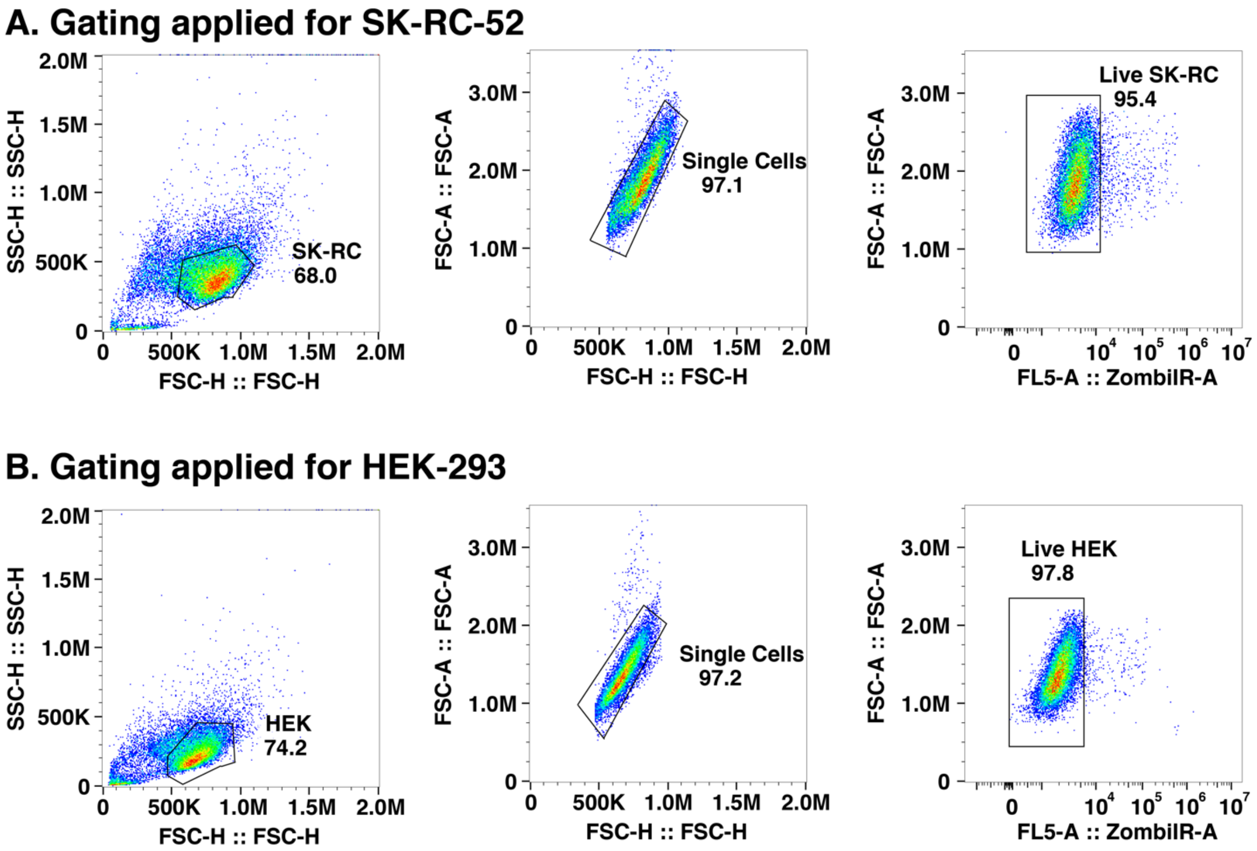


**Figure S18. Gating applied for the flow cytometry analysis of CAIX-positive SK-RC-52 cells (A.) and CAIX-negative HEK-293 cells (B).**

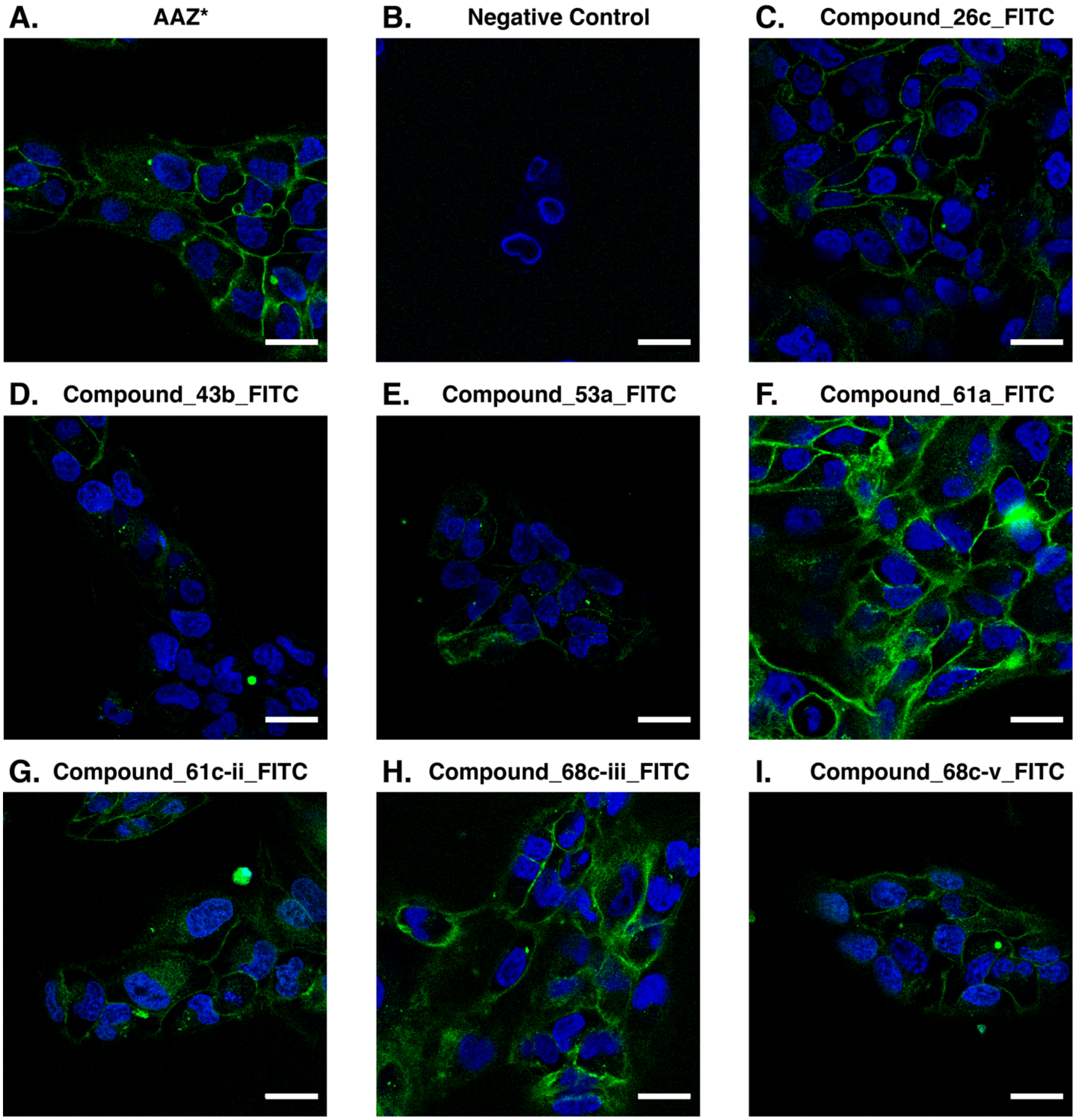


**Figure S19. Confocal fluorescence microscopy images of CAIX-positive SK-RC-52 cells after incubation with FITC conjugated compounds. GREEN = compounds (Fluorescein), BLUE = cell nuclei (DAPI staining). Scale bar = 30 μm.**

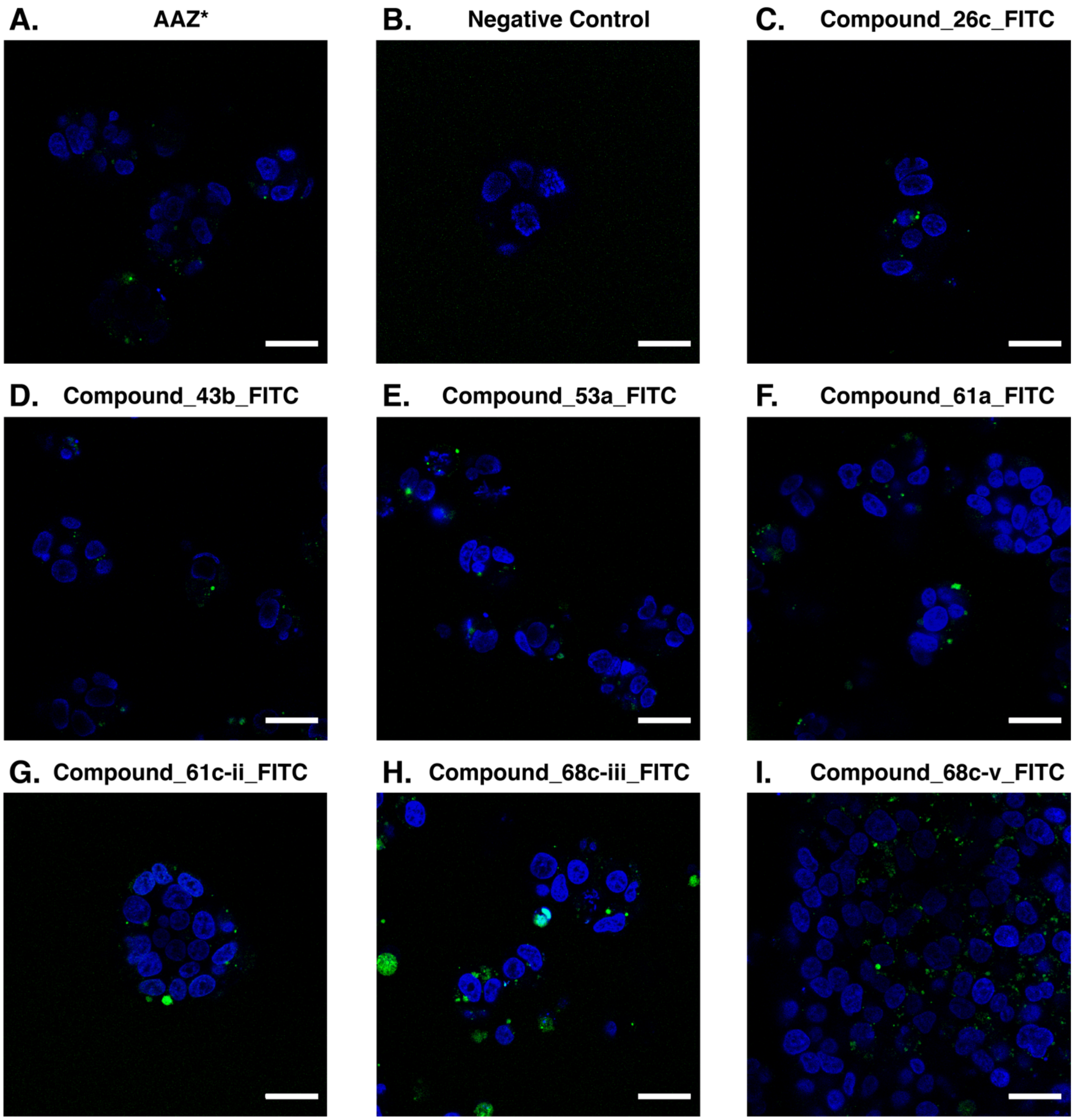


**Figure S20. Confocal fluorescence microscopy images of CAIX-negative HEL-293 cells after incubation with FITC conjugated compounds. GREEN = compounds (Fluorescein), BLUE = cell nuclei (DAPI staining). Scale bar = 30 µm.**

### Appendix I – HR-MS spectra

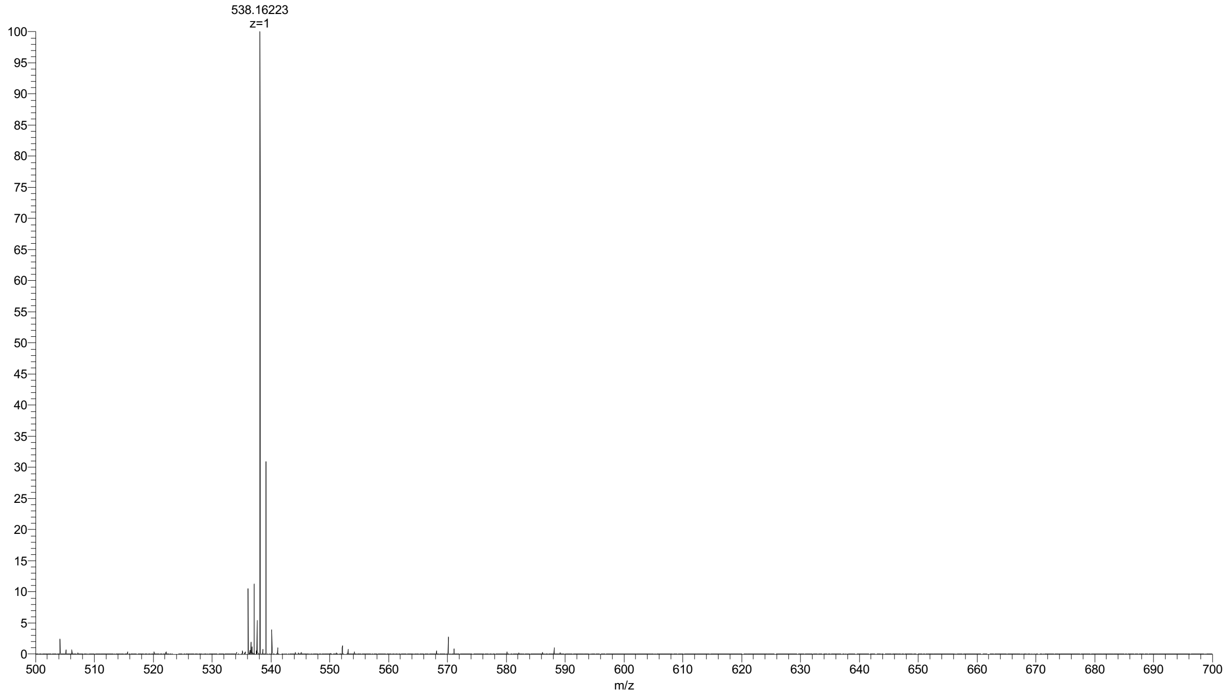


**Figure S21. HR-MS of intermediate 1.**

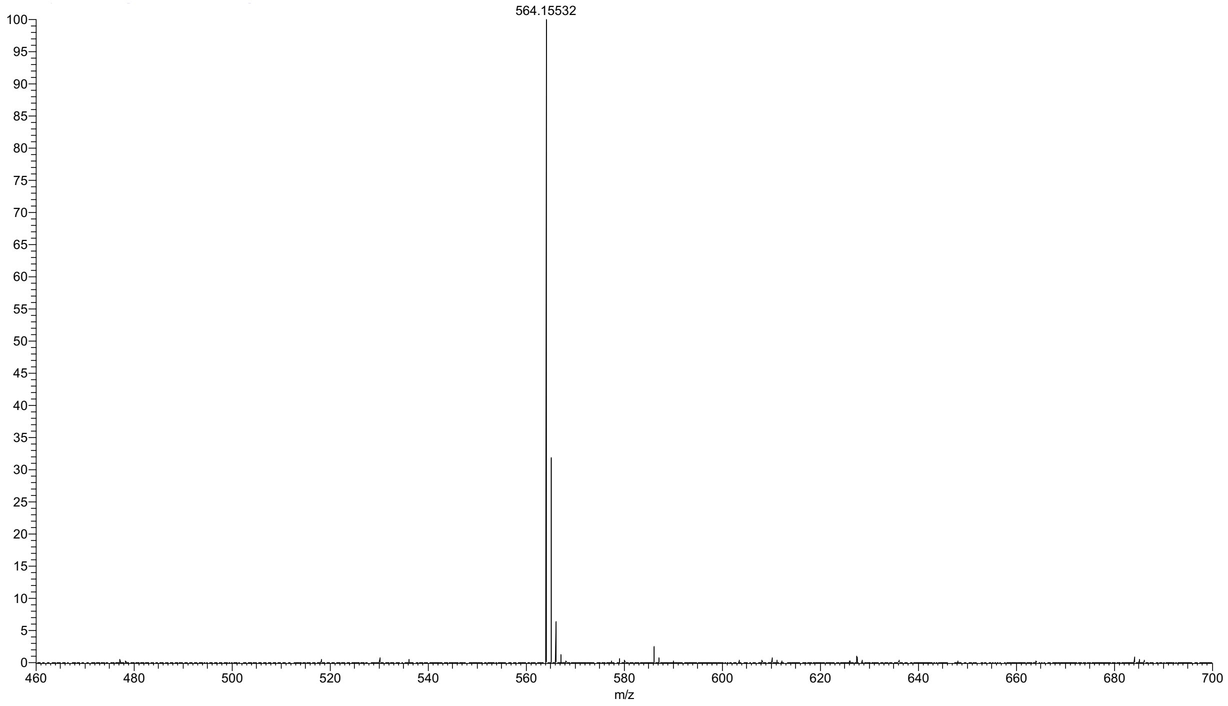


**Figure S22. MS of intermediate 2.**

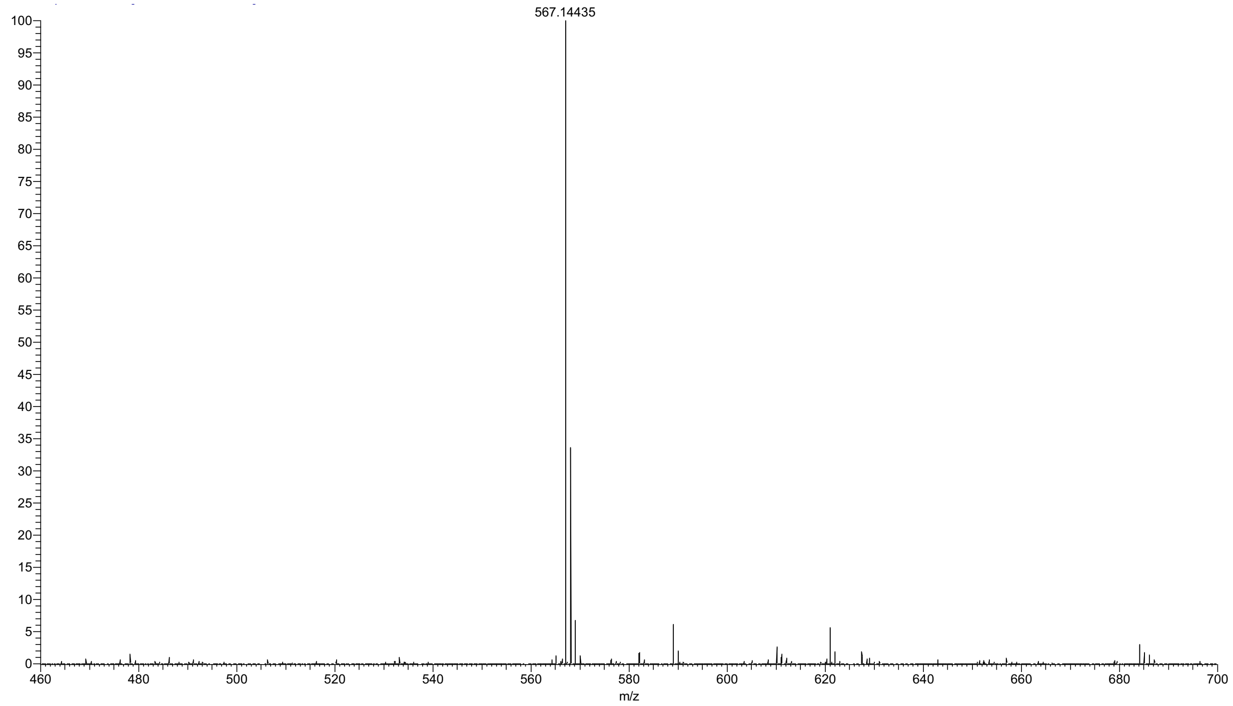


**Figure S23. MS of intermediate 3.**

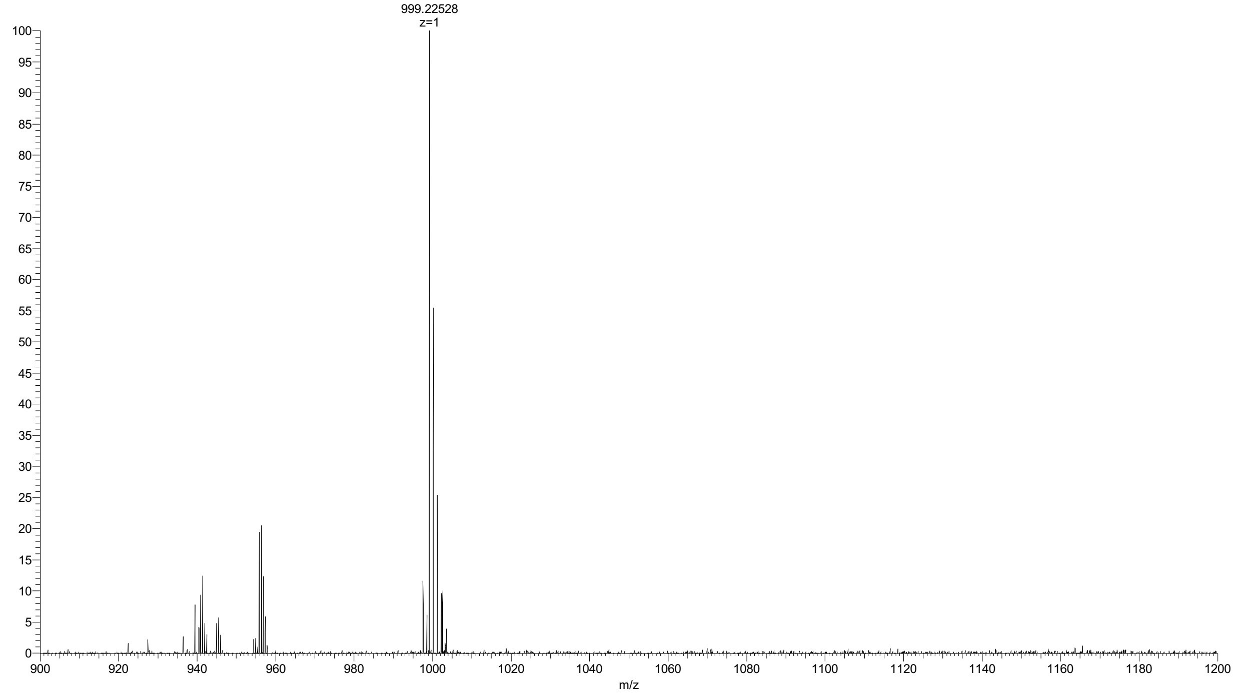


**Figure S24. HR-MS of compound_26c_FITC.**

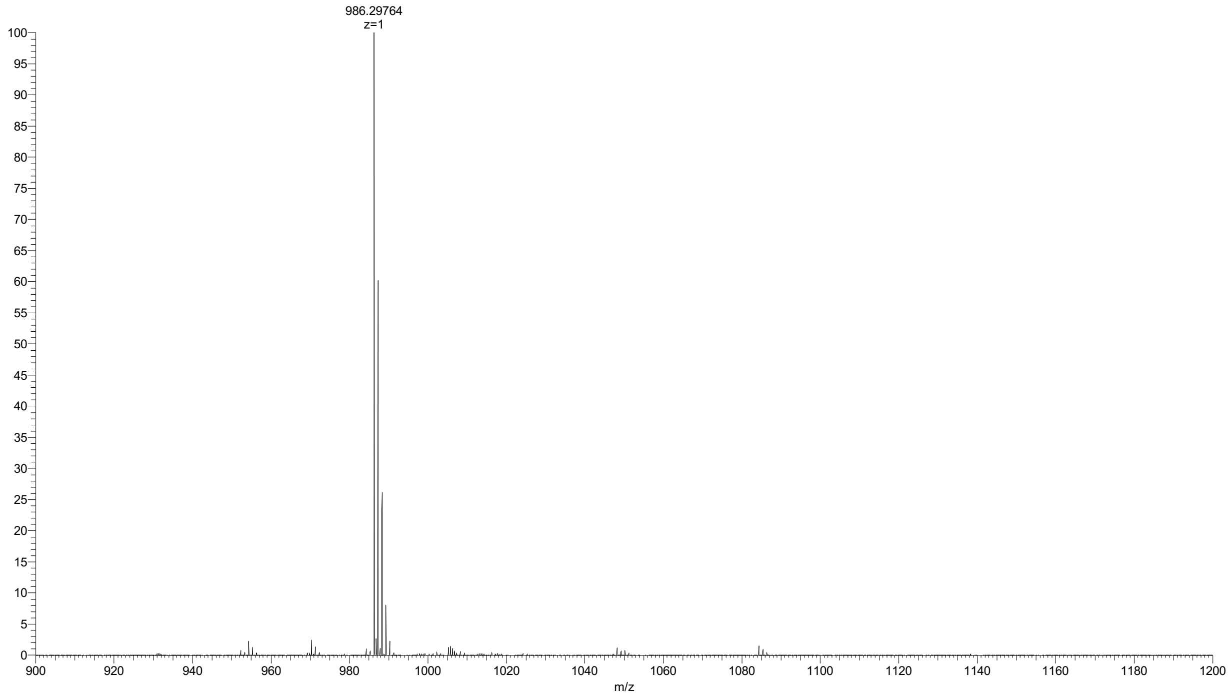


**Figure S25. HR-MS of compound_68c-v_FITC.**

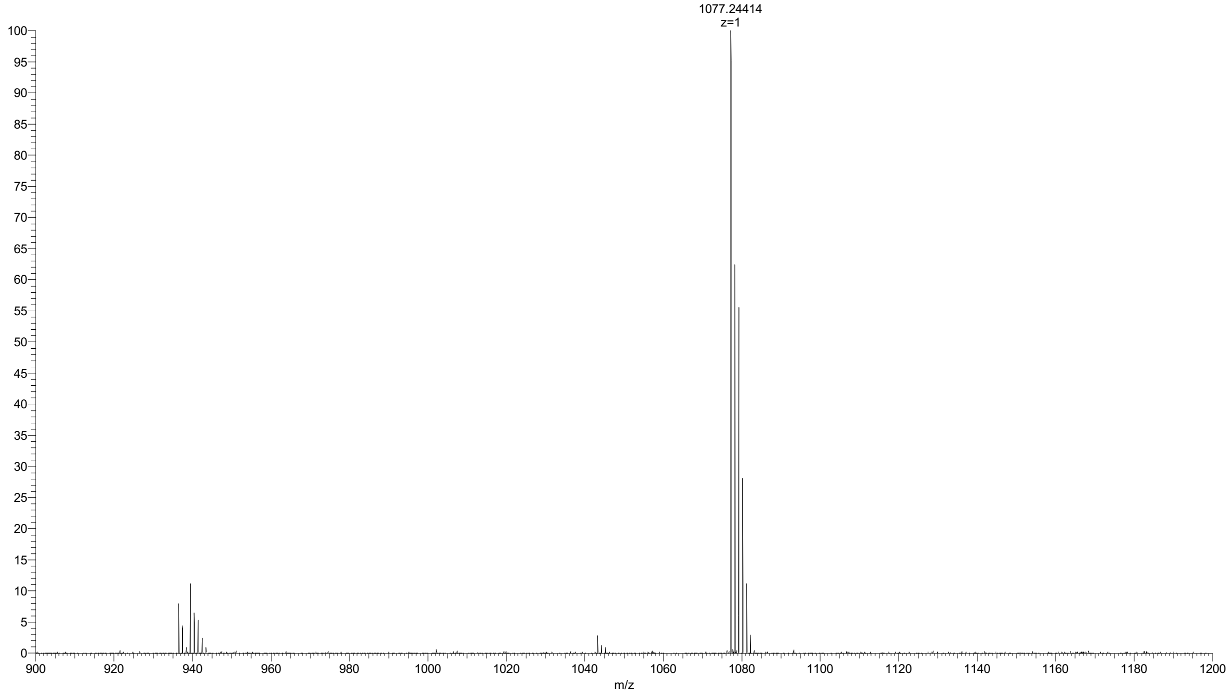


**Figure S26. HR-MS of compound_61c-ii_FITC.**

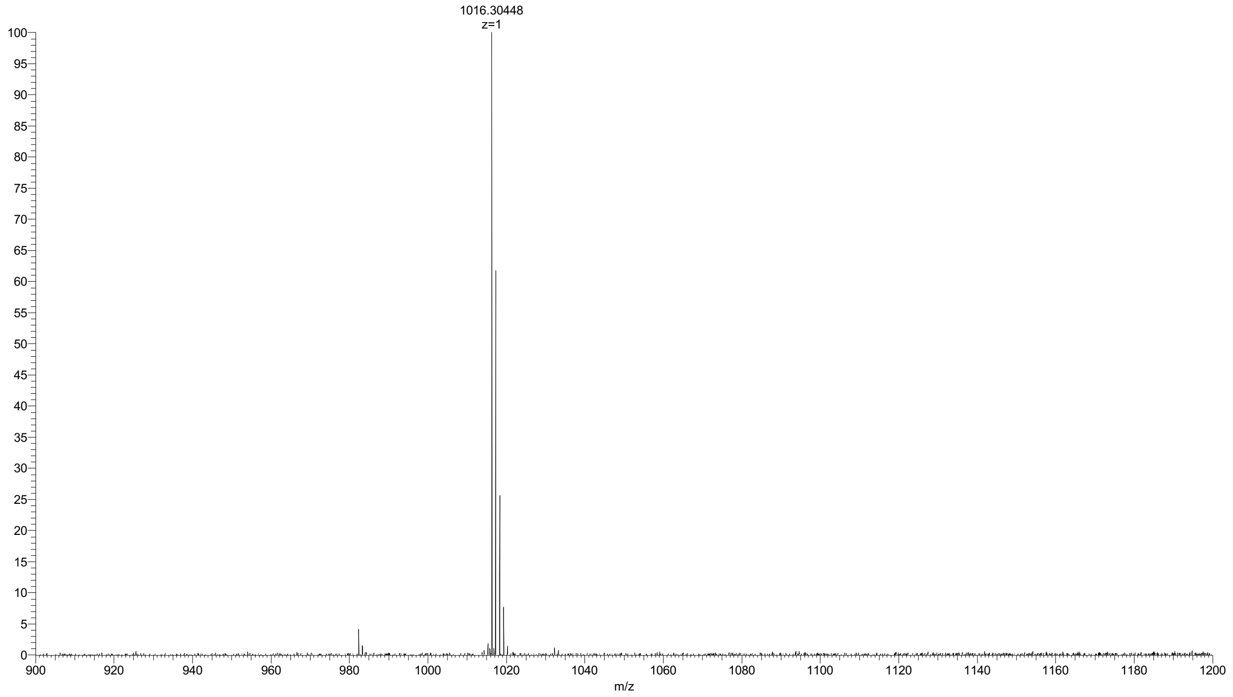


**Figure S27. HR-MS of compound_68c-iii_FITC.**

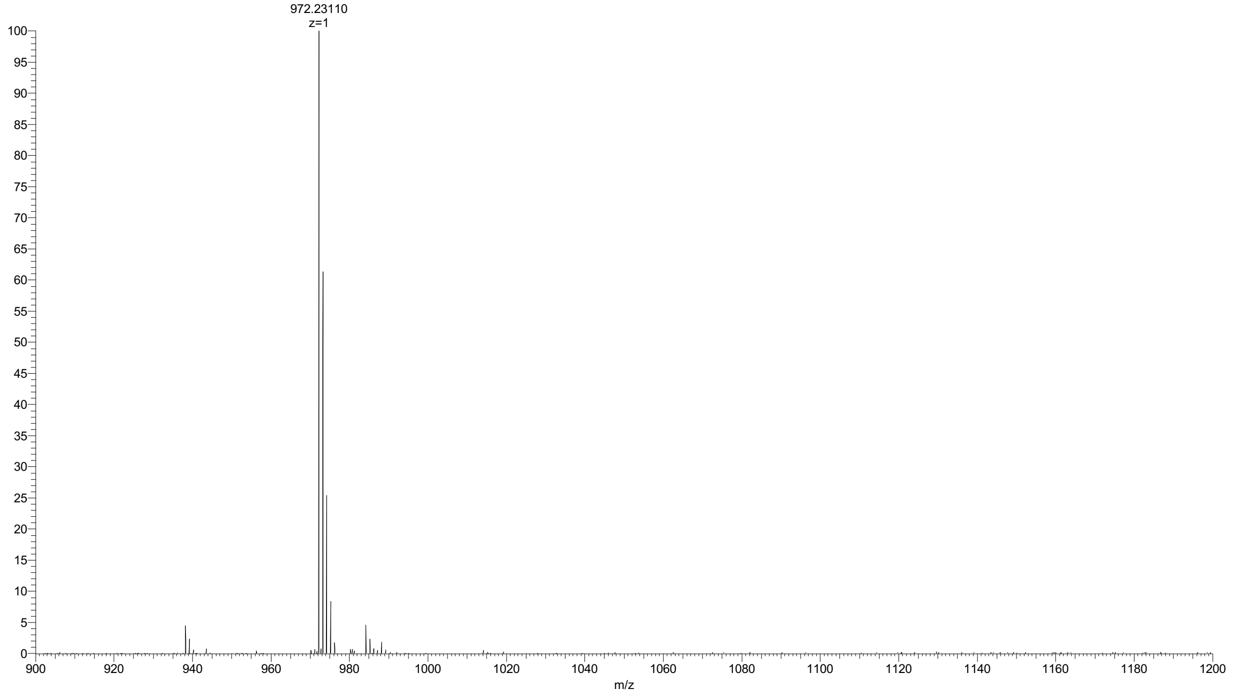


**Figure S28. HR-MS of compound_43b_FITC.**

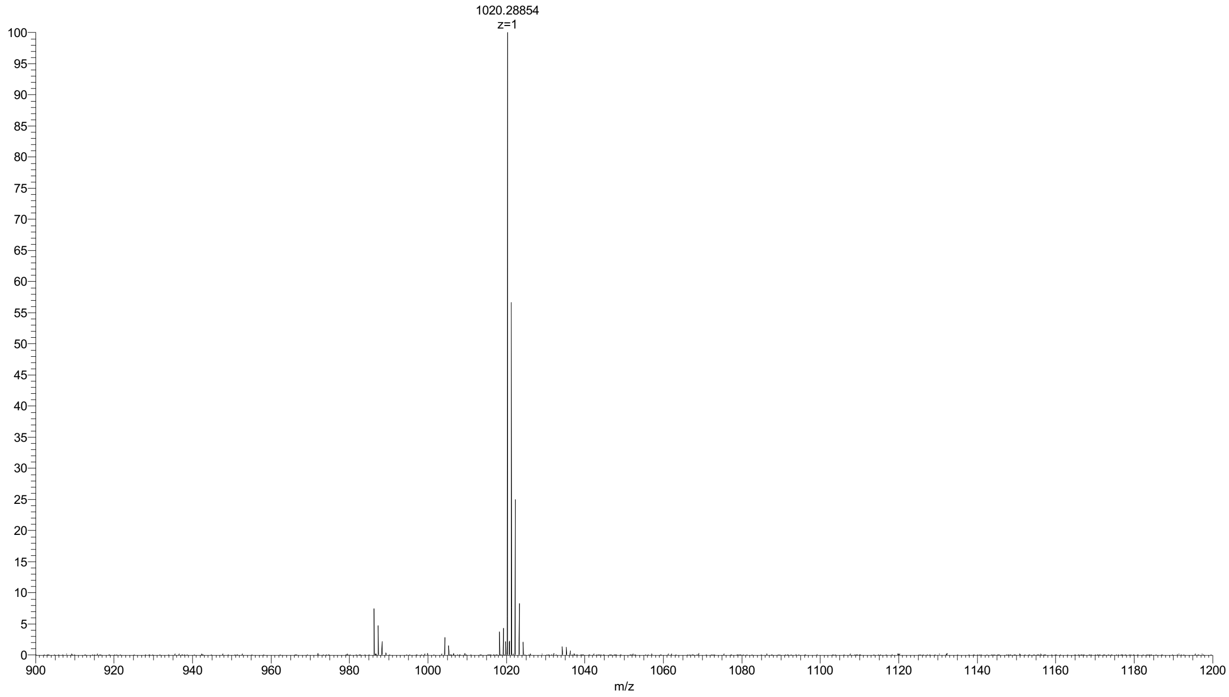


**Figure S29. HR-MS of compound_53a_FITC.**

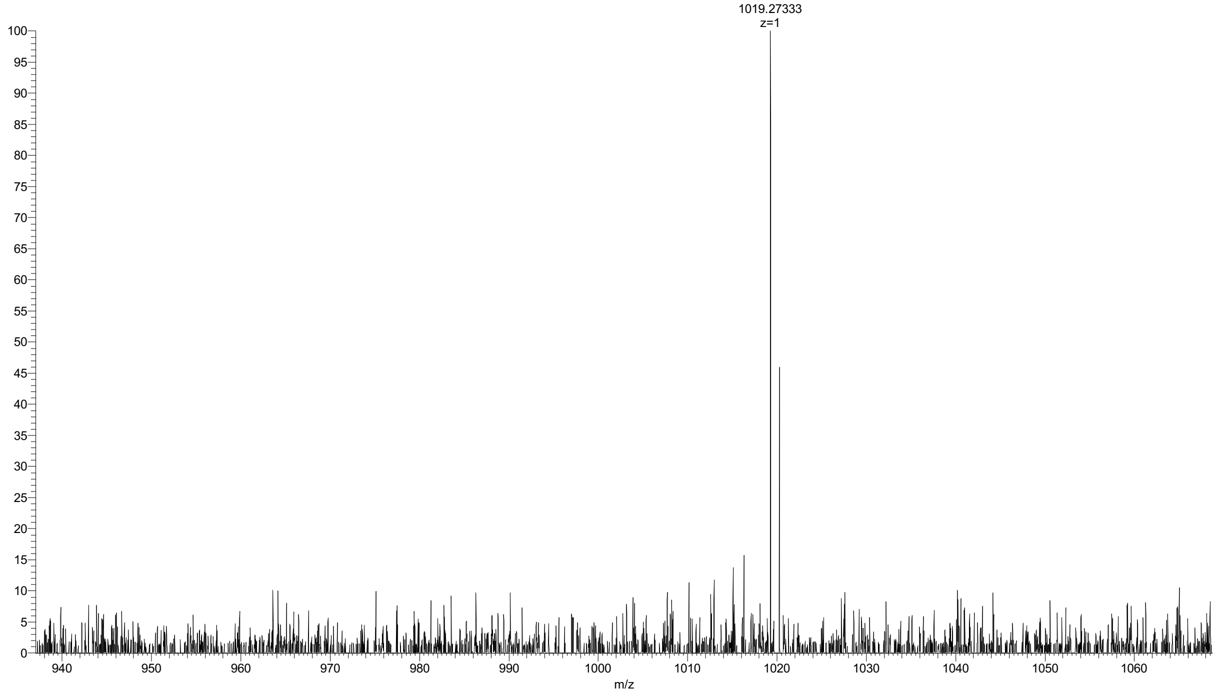


**Figure S30. HR-MS of compound_61a_FITC.**

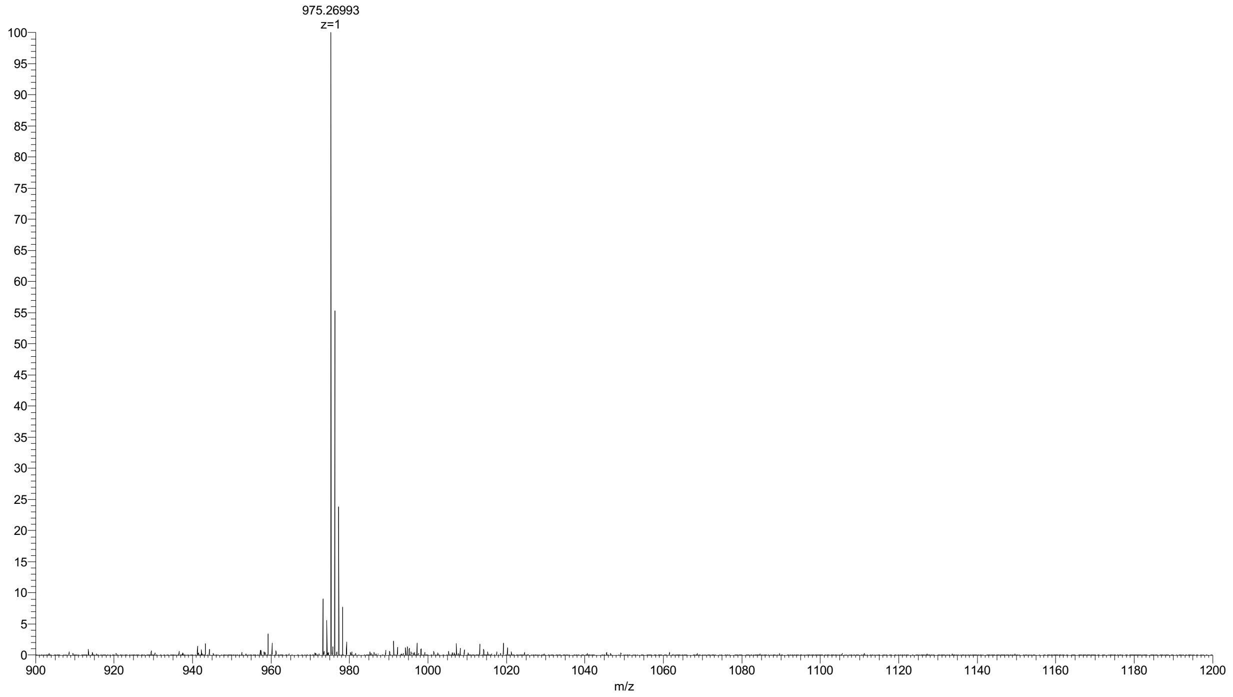


**Figure S31. HR-MS of compound_109b-ii_FITC.**

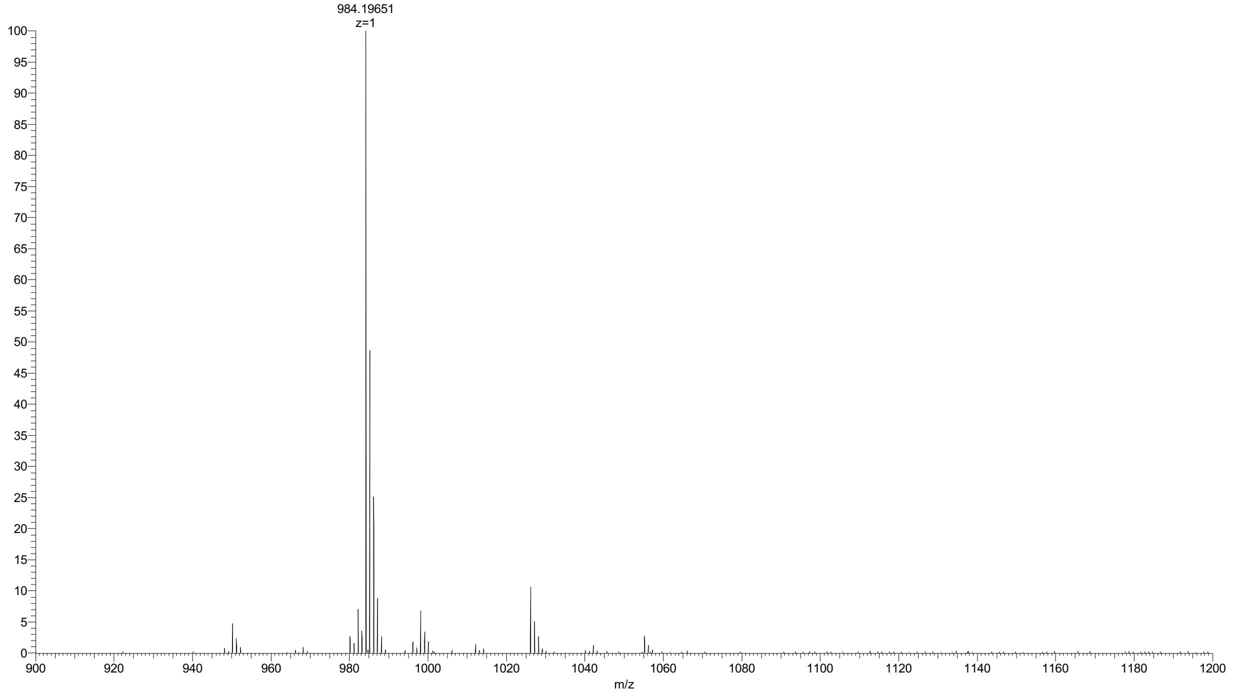


**Figure S32. HR-MS of AAZ*.**

### Appendix 2 – HPLC spectra

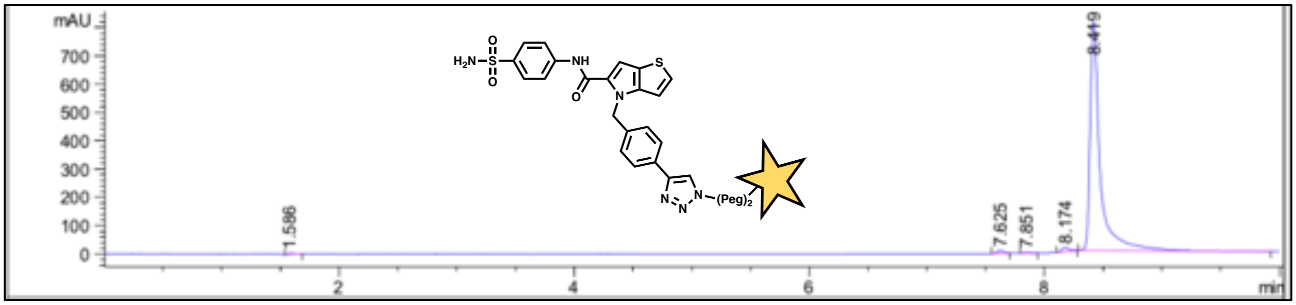


Figure S33. HPLC trace of compound_26c_FITC.

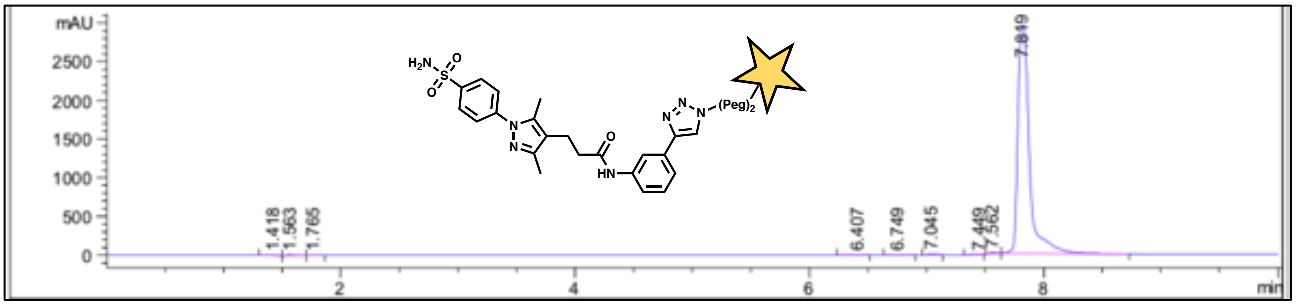


Figure S34. HPLC trace of compound_68c-v_FITC.

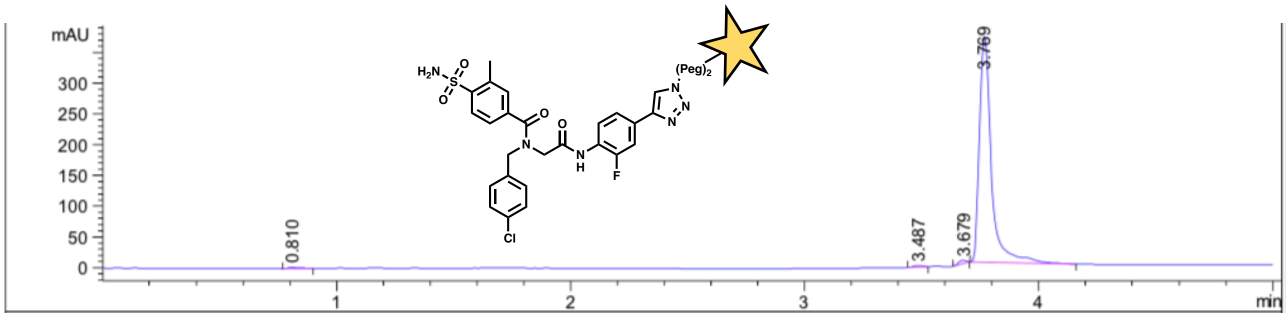


Figure S35. HPLC trace of compound_61c-ii_FITC.

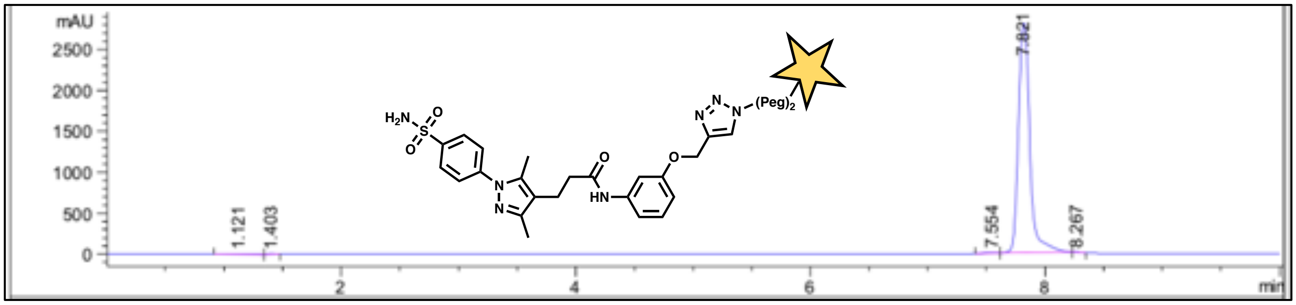


Figure S36. HPLC trace of compound_68c-iii_FITC.

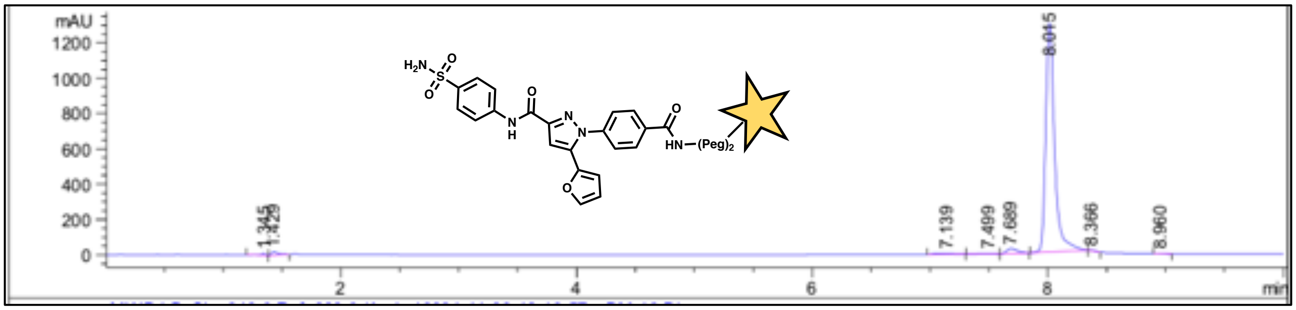

**Figure S37. HPLC trace of compound_43b_FITC.**

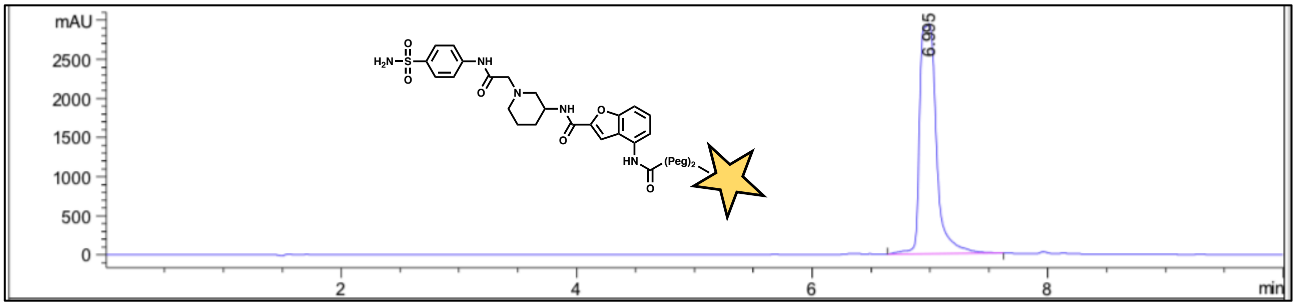

**Figure S38. HPLC trace of compound_53a_FITC.**

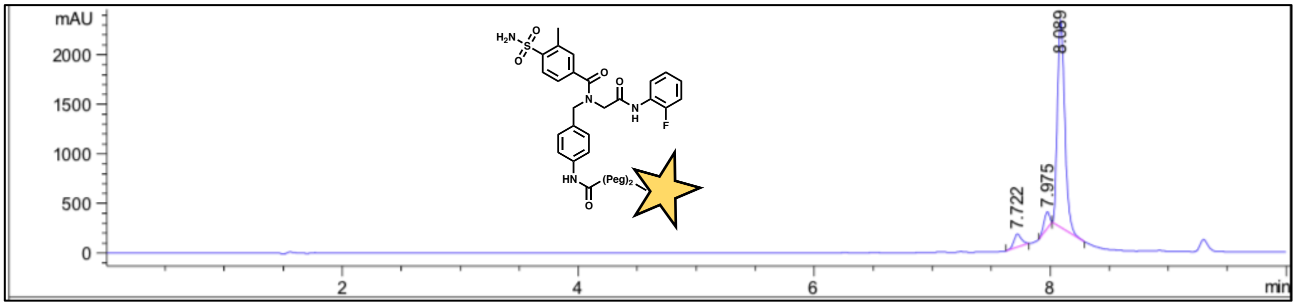

**Figure S39. HPLC trace of compound_61a_FITC.**

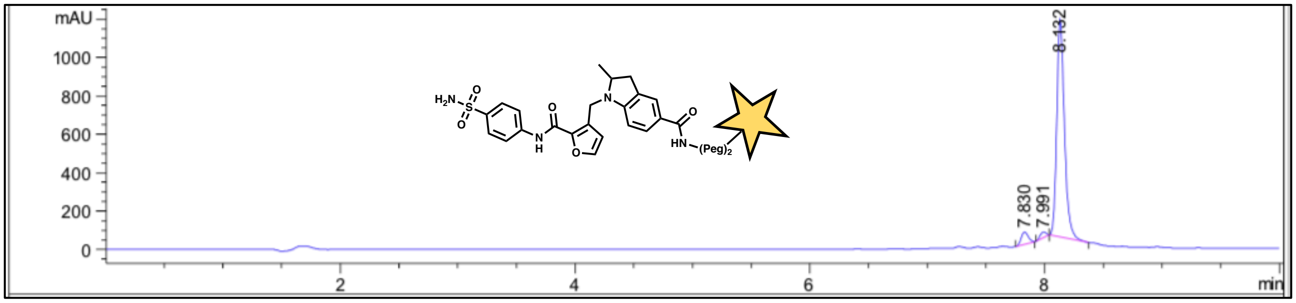

**Figure S40. HPLC trace of compound_109b-ii_FITC.**
